# Design, Construction, and Validation of a Yeast-Displayed Chemically Expanded Antibody Library

**DOI:** 10.1101/2024.05.29.596443

**Authors:** Arlinda Rezhdo, Rebecca L. Hershman, James A. Van Deventer

## Abstract

*In vitro* display technologies, exemplified by phage and yeast display, have emerged as powerful platforms for antibody discovery and engineering. However, the identification of antibodies that disrupt target functions beyond binding remains a challenge. In particular, there are very few strategies that support identification and engineering of either protein-based irreversible binders or inhibitory enzyme binders. Expanding the range of chemistries in antibody libraries has the potential to lead to efficient discovery of function-disrupting antibodies. In this work, we describe a yeast display-based platform for the discovery of chemically diversified antibodies. We constructed a billion-member antibody library that supports the presentation of a range of chemistries within antibody variable domains via noncanonical amino acid (ncAA) incorporation and subsequent bioorthogonal click chemistry conjugations. Use of a polyspecific orthogonal translation system enables introduction of chemical groups with various properties, including photo-reactive, proximity-reactive, and click chemistry-enabled functional groups for library screening. We established conjugation conditions that facilitate modification of the full library, demonstrating the feasibility of sorting the full billion-member library in “protein-small molecule hybrid” format in future work. Here, we conducted initial library screens after introducing *O*-(2-bromoethyl)tyrosine (OBeY), a weakly electrophilic ncAA capable of undergoing proximity-induced crosslinking to a target. Enrichments against donkey IgG and protein tyrosine phosphatase 1B (PTP1B) each led to the identification of several OBeY-substituted clones that bind to the targets of interest. Flow cytometry analysis on the yeast surface confirmed higher retention of binding for OBeY-substituted clones compared to clones substituted with ncAAs lacking electrophilic side chains after denaturation. However, subsequent crosslinking experiments in solution with ncAA-substituted clones yielded inconclusive results, suggesting that weakly reactive OBeY side chain is not sufficient to drive robust crosslinking in the clones isolated here. Nonetheless, this work establishes a multi-modal, chemically expanded antibody library and demonstrates the feasibility of conducting discovery campaigns in chemically expanded format. This versatile platform offers new opportunities for identifying and characterizing antibodies with properties beyond what is accessible with the canonical amino acids, potentially enabling discovery of new classes of reagents, diagnostics, and even therapeutic leads.

**Table of Contents Figure:** 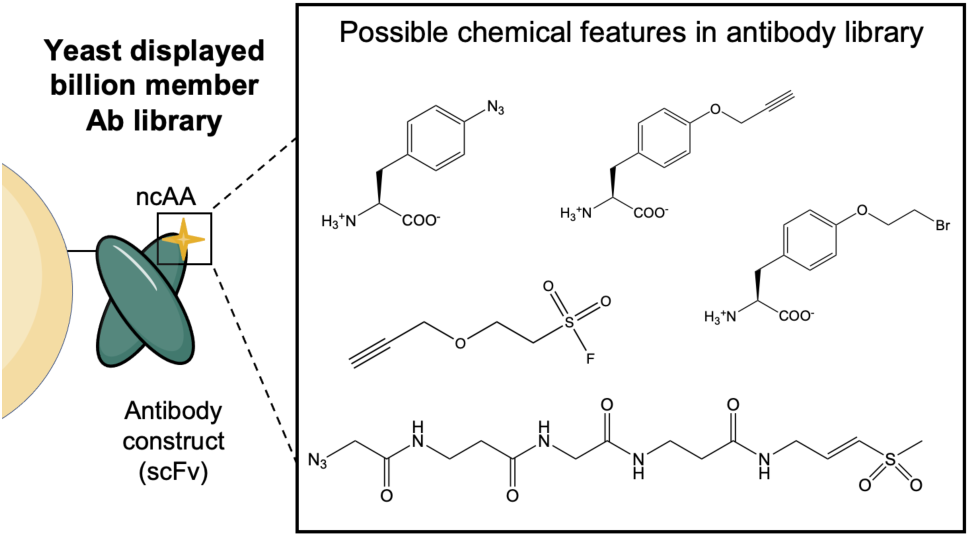

## Introduction

Modern medicine and biotechnology rely heavily on antibodies as an important class of therapeutics, as key tools for basic research, and as critical components of diagnostics^1–3^. Yeast display, phage display, and additional display technologies have emerged as powerful *in vitro* discovery platforms that enable the discovery of antibodies that would be challenging or impossible to discover with immunization and hybridoma-based strategies^4–8^. Importantly, *in vitro* platforms enable construction and screening of synthetic antibody libraries^9^, which encode sequences beyond those generated in the human immune system. Strategic use of display technologies enables the discovery and engineering of antibody variants with combinations of desirable properties (high affinity, low polyspecificity, low self-association, etc.) that can be difficult to identify from natural repertoires^10–12^. In addition, combinatorial screens with synthetic diversities have enabled researchers to gain powerful insights into the roles of different frameworks, complementarity determining regions (CDRs), and amino acid diversities in dictating binding interactions and other antibody properties^4–6,9,13,14^. These *in vitro* screening platforms have greatly enhanced antibody discovery and engineering.

In parallel with advances in display technologies, there are rapidly expanding advances in the integration of chemical functionalities into antibodies to establish innovative classes of antibody-based therapeutics. Bioconjugate chemistry^2,15–17^, glycoengineering^18,19^, and genetically encoded noncanonical amino acids (ncAAs)^11,20,21^ all enable routes to produce “chemically diversified” antibodies *in vitro*. Advances in the field of antibody-drug conjugates (ADCs)^15,22,23^ serve as a compelling testament to the deliberate integration of chemical entities into antibody frameworks. Further, recent reports have demonstrated approaches to presenting added chemical functionality within antibody binding domains to induce covalent target engagement^24^. Several groups have reported strategies for engineering covalent antibodies with envisioned applications such as radiopharmaceuticals with enhanced potency^25,26^. However, numerous questions remain regarding how best to “chemically diversify” antibodies.

*In vitro* display technologies provide opportunities to conduct high throughput investigations to address these outstanding questions. Early work with phage display and *E. coli* display platforms demonstrated the technical feasibility of engineering ncAA-containing antibodies with expanded chemical functionalities in high throughput^13,27,28^. In addition, our lab has integrated yeast display with ncAAs and demonstrated semi-rational strategies for endowing antibodies and other binding proteins with bioorthogonal chemistries^29^, photocrosslinking^30,31^, and spontaneous crosslinking capabilities^31^. The power of expanding chemical functionality into display technologies has been demonstrated in myriad ways using peptide-based screens in mRNA display^32–34^ and phage display formats^35^. Yet, there remain extensive opportunities to discover antibody hybrids with unique properties through the generation and screening of chemically diversified antibodies. Such synergistic properties are largely not accessible in either proteins or small molecules alone.

In this work, we report the first chemically diversified antibody library in yeast display format. By leveraging genetic code expansion and a promiscuous orthogonal translation system, we successfully constructed a multi-modal, billion-member synthetic antibody library capable of encoding one of several ncAA side chain functionalities. The identity of the 21^st^ amino acid can be precisely controlled during the library induction process. This library can be further modified using click chemistry, enhancing its potential to generate chemically diverse antibody variants. By incorporating an azide- or alkyne-containing ncAA, the library can be easily modified using azide-alkyne cycloadditions, enabling the construction of one billion unique “protein-small molecule hybrids” covering a wide range of chemistries. As a proof-of-concept for high-throughput discovery with this library, we prepared the library with a weakly electrophilic ncAA and screened for clones capable of binding to a model antigen, donkey IgG, as well as a therapeutically relevant target, protein tyrosine phosphatase 1B (PTP1B). In both cases, we identified several ncAA-containing clones capable of binding to the targets. Individual clones contained ncAA substitutions at one of several different positions and exhibited moderate binding affinities. We observed some evidence for possible spontaneous crosslinking on the yeast surface, but experiments in solution were inconclusive, suggesting that the ncAA used may not be reactive enough to drive rapid spontaneous crosslinking in the clones isolated here. Nonetheless, these efforts validate discovery of chemically expanded antibodies in yeast display format, with the library design providing a range of opportunities to pursue chemistry-driven antibody discovery. We expect this unique, chemically expanded library to enable novel approaches for antibody discovery in high throughput.

## Results

### Multi-modal library design and construction

We designed a “multi-modal” library by combining simple synthetic antibody diversity with several potential positions for ncAA incorporation (Figure 1). The single chain variable fragment scaffold contains framework regions found in the therapeutic antibody trastuzumab (Herceptin)^5,9^, and complementarity determining region H3 (CDR-H3) diversity includes loop lengths and amino acid diversities that capture important features of naturally occurring antibody diversity^30^ (Figure 1A). Using degenerate “XYZ” codons with customized oligonucleotide frequencies (Figure 1A), clones have 5-13 positions randomized with “antibody-like” diversity followed by one position encoding A/G and one encoding F/I/L/M, all flanked between K and D residues, resulting in CDR-H3 loop lengths ranging from 9-17. For each CDR-H3 loop length, six locations were chosen as possible ncAA substitution sites in response to a TAG codon (Figure 1A). These were selected to position the ncAA in a surface accessible configuration in or near the CDRs (other than CDR-H3) in the light chain and heavy chain of the scFv. Four positions were selected in the light chain for TAG substitution: (1) position L1 located at the N-terminus of the scFv; (2) position L28 located in CDR-L1; (3) position L50 located in CDR-L2; and (4) position L93 located in CDR-L3. Two positions were selected in the heavy chain for substitution: (1) position H31 located in CDR-H1; and (2) position H54 located in CDR-H2. Since amino acids in CDRs frequently participate in antigen binding, positioning ncAAs within surface-accessible regions of these loops increases the chances that these residues can contribute to target binding either directly or via moieties installed via bioorthogonal conjugation.

**Figure 1.**
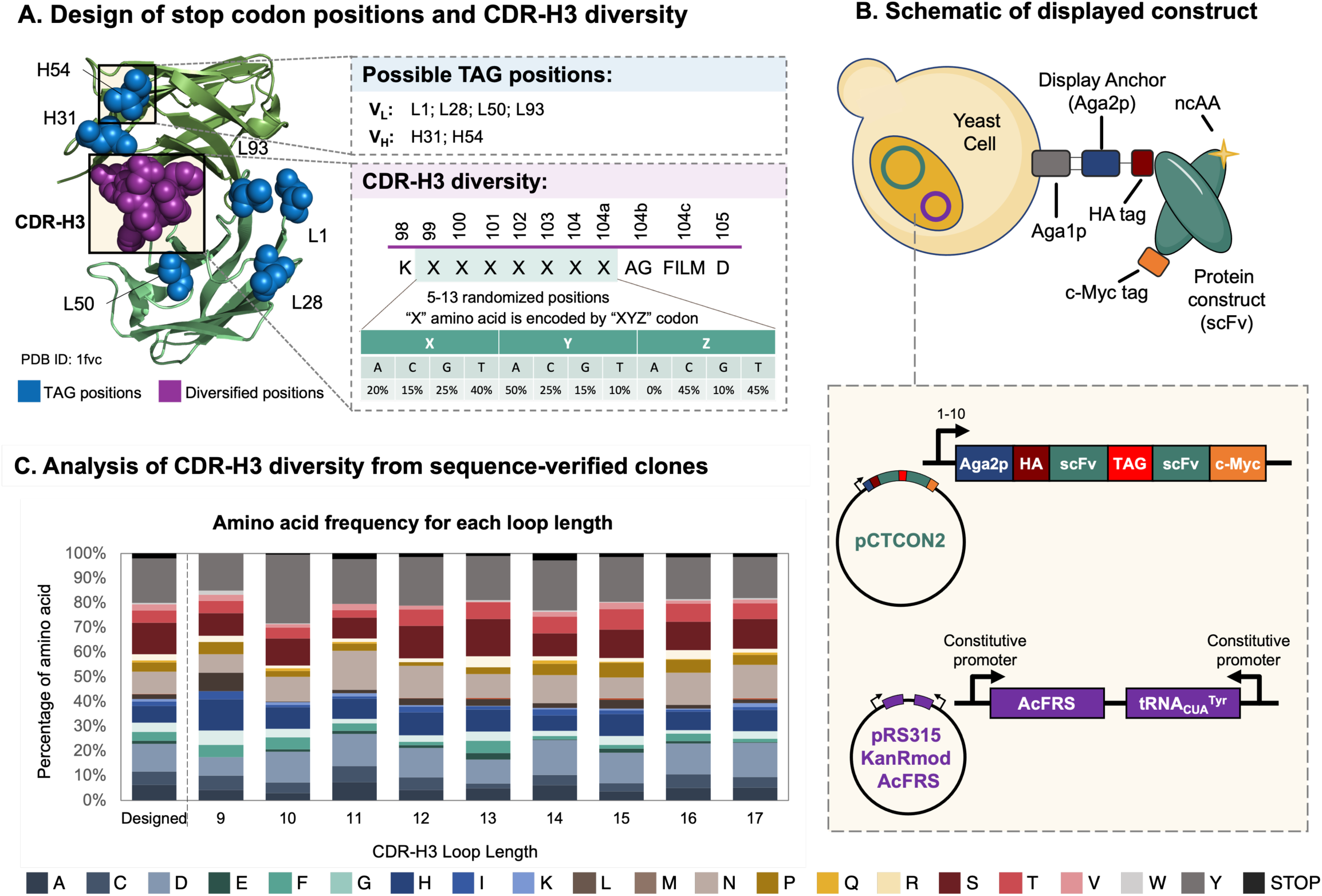
Design and construction of “multi-modal” library. A) Design of CDR-H3 diversification scheme and TAG substitutions in the light and heavy chains for ncAA positioning (Protein Data Bank ID: 1fvc). B) Schematic overview of library in yeast display format. pCTCON2 backbone encodes antibody single-chain variable fragment library under control of the Gal1-10 promotor; inducible sequence includes Aga2p display anchor, HA epitope tag, diversified scFv with ncAA incorporation site, and c-Myc epitope tag. A second plasmid encodes the orthogonal translation system (AcFRS and tRNA_CUA_), both under constitutive promotors, which will incorporate ncAAs in reponse to a TAG codon. C) Amino acid frequency in the diversified CDR-H3 from Sanger sequencing data.

We prepared the library by constructing a series of 54 sublibraries, each with a distinct CDR-H3 loop length and ncAA position pairing (9 loop lengths each paired with 6 individual ncAA positions, resulting in 54 unique framework-loop combinations). To obtain such pairings, stop codons were introduced into the scFv sequence via point mutations and amplifications with primers encoding the CDR-H3 diversity. The resulting PCR products and linearized yeast display plasmid pCTCON2 were transformed via homologous recombination into *Saccharomyces cerevisiae* RJY100 cells. These cells were previously transformed with a plasmid encoding the *p*-acetylphenylalanyl-tRNA synthetase/tRNA^CUA^ orthogonal translation system (OTS; AcFRS^36,37^), which supports a range of aromatic ncAA incorporations in response to the amber stop codon at moderate efficiencies (approximately 15% efficiency of wild-type protein translation)^37^ (Figure 1B). Following transformation, each sublibrary was evaluated for (1) number of transformants via plating and colony counting (SI Figure 1); (2) full-length display levels in the absence and presence of a ncAA during induction of yeast display (SI Figure 2, SI Figure 3); (3) fraction of truncated clones (defined here as detection of HA+, c-Myc– clones following induction in the presence of a ncAA; SI Figure 4); and (4) sequence validation of individual clones (see detailed criteria for sublibrary quality control in Supporting Information). Sequence validation of each sublibrary was performed via Sanger sequencing of 5-10 clones (SI Table 1). The observed frequencies of each amino acid and nucleotide in the region encoding CDR-H3 were comparable to the designed frequencies^38^ (Figure 1C, SI Figure 5), and 92.2% of the sequenced clones were unique. These data indicate that the constructed library is consistent with the designed diversity. Although deep sequencing could give a more thorough picture of library quality, Sanger sequencing was sufficient to verify key elements of the design. Out of 54 initial sublibraries, 49 were deemed to be of sufficiently high quality to be included in the final, pooled library. However, one of the 49 did not regrow from glycerol stocks. Thus, 48 sublibraries were pooled for a total of 1.1×10^9^ transformants (SI Figure 1). Overall, the billion-member scFv library contains amino acid, loop length, and TAG codon position diversities consistent with the initial library design. This set the stage for (1) investigating the range of chemistries accessible through ncAA incorporation and bioorthogonal conjugations; and (2) high throughput antibody discovery with the chemically diversified library.

### Evaluation of library chemical versatility

To determine the range of chemistries that can be integrated within the library, we sought to evaluate (1) display levels after incorporation of a range of ncAAs; and (2) conjugation efficiencies with a range of small molecules. Covalent target engagement and efficient bioorthogonal conjugation both require amino acid side chain functionalities that are not available within the conventional genetic code. We evaluated our ability to genetically encode a set of functional groups that enable covalent binding or efficient conjugations by inducing the library in the presence of each of six ncAAs. Analysis of N-terminal and C-terminal display levels via flow cytometry indicated that AcFRS supports full-length scFv display following induction of the library in the presence of each of the following ncAAs: *p*-azido-L-phenylalanine (AzF; **1**), *p*-propargyloxy-L-phenylalanine (OPG; **2**), *p*-acetyl-L-phenylalanine (AcF; **3**), *O*-methyl-L-tyrosine (OmeY; **4**), 4-iodo-L-phenylalanine (IPhe; **5**) and *O*-(2-bromoethyl)tyrosine (OBeY; **6**) (Figure 2A). The fractions of full-length and truncated clones calculated for each ncAA further demonstrate that the suppression machinery supports the incorporation of all six ncAAs in this system (SI Figure 6). This data confirms the library versatility at the level of ncAA side chains: the six translationally active ncAAs include crosslinkable side chains as well as groups that can undergo efficient conjugation reactions^15,27,39–41^. For example, covalent antibody discovery could be conducted using either the photocrosslinkable ncAA AzF (**1**)^30,31^ or the spontaneously crosslinkable ncAA OBeY (**6**)^31^. Additionally, both the azide side chain of AzF (**1**) and alkyne functionality of OPG (**2**) enable bioorthogonal click chemistry reactions^29,30^.

**Figure 2.**
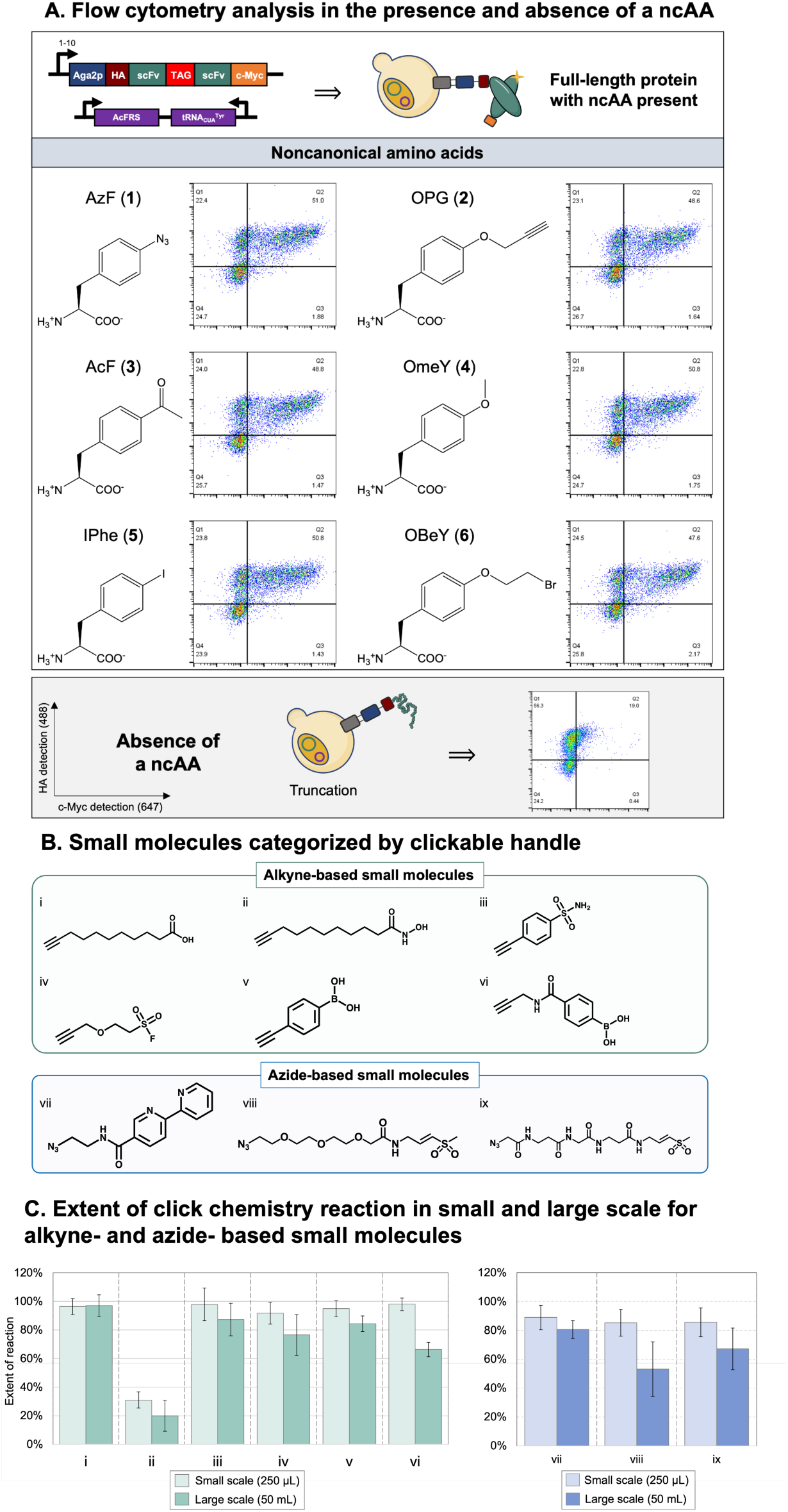
Chemically diversified antibody library via ncAA incorporation and CuAAC. A) Flow cytometry dot plots (HA and c-Myc detection) of displayed antibodies following induction in the presence of one of six ncAAs (1 mM concentration): AzF (**1**), OPG (**2**), AcF (**3**), OmeY (**4**), IPhe (**5**) and OBeY (**6**), as well as a flow cytometry dot plot (HA and c-Myc detection) following induction in the absence of ncAAs. B) Six alkyne-functionalized pharmacophores (i) 10-undecynoic acid, (ii) N-hydroxy-10-undecynamide, (iii) 4-ethynylbenzene-1-sulfonamide, (iv) 2-(2-propyn-1-yloxy)ethane-sulfonyl fluoride, (v) 4-(dihydroxy-borophenyl) acetylene, (vi) 4-(propargyl-aminocarbonyl)phenylboronic acid and three azide-functionalized pharmacophores (vii) N-(2-azidoethyl)-[2,2’-bipyridine]-5-carboxamide, (viii) N_3_-PEG_4_-VS, (ix) N_3_-bAGbA-VS used in this work to evaluate chemical reactivity. C) Extent of click chemistry reactions at small and large scale (small scale: 2×10^6^ cells in 250 μL; large scale: 1×10^10^ cells in 50 mL) with the naïve library induced with either AzF (compatible for CuAAC with pharmacophores i-vi) or OPG (compatible for CuAAC with pharmacophores vii-ix) and the small molecules in panel B.

We also investigated the use of copper-catalyzed azide-alkyne cycloadditions (CuAAC) at large scale as a way to further modify the full library prior to screening. While in previous work we have routinely conducted CuAAC at small scale on the yeast surface (2×10^6^ cells in 250 µL reaction)^29,42^, chemistry-driven sorts require modification of 1×10^10^ cells to ensure tenfold oversampling of a billion-member library during the initial round of screening. Because CuAAC reaction efficiency can change as a function of cell density^29,43–45^, we evaluated CuAAC conjugations at small and large scales for a range of small molecules (Figure 2B and C). Reactions with 9 small molecules containing a range of chemical moieties (Figure 2B) were conducted on cells from the library in 250 µL total reaction volume at low cell density (2×10^6^ modified cells; 8×10^6^ cells/mL) and in 50 mL total reaction volume at high cell density (1×10^10^ modified cells; 2×10^8^ cells/mL). For both conditions, the small molecule was added to the reaction at a final concentration of 1 mM. We used “extent of reaction” calculations as a measure of how well a small molecule reacts with the ncAA-containing naïve library displayed on the yeast surface. An extent of reaction of 100% indicates that a small molecule reacts with all of the reactive ncAAs in the sample (as determined by controls with corresponding biotin probes), whereas an extent of reaction of 0% indicates undetectable levels of reaction (Further experimental details are available in *Materials and Methods*, and detailed protocols have been published elsewhere^46^). Calculated extents of reaction obtained here indicate that numerous small molecules react to a similar extent under both small- and large-scale conditions (Figure 2C). Overall, the data indicates that the library can be modified with a broad range of azide- and alkyne-based probes containing a variety of chemical warheads and linker structures. Interestingly, this even includes small molecules with metal-binding functionality that could interfere with CuAAC; this observation is consistent with other surveys of chemical reactivity performed on the yeast surface in our lab^29^. However, not all small molecules react efficiently at large scale. This reduction in reactivity may be due to increased cell density or due to increased oxygenation in 50 mL reactions; high oxygen content in solution is known to impede CuAAC reactions, even for ligands reported to be oxygen-tolerant^43^. While adjustments to reaction conditions may facilitate an even broader range of chemical group installations, the simple conditions used here already enable a versatile range of chemical groups to be efficiently installed within the library.

Overall, the range of ncAA side chains that can be incorporated biosynthetically and the range of small molecules that can be installed using bioorthogonal chemistry make this antibody library multi-modal. The reactive side chains of AzF (photo-reactive azide) and OBeY (electrophilic bromoethyl) can be used in discovery of covalent binders to targets of interests, while the clickable side chains of AzF (azide) and OPG (alkyne) can be used in conjugations to further diversify chemistries presented within displayed antibodies. Small molecules successfully conjugated in this work contain a variety of functional groups including carboxylic acid (metal chelation), boronic acid (Suzuki coupling, glycan targeting), sulfonyl fluoride (SuFEx chemistry), bipyridine (metal chelation), and vinyl sulfone (thiol binding). This has the potential to support discovery efforts for protein-small molecule hybrids that interact with targets via a variety of functionalities that are not normally accessible within antibody structures.

### Chemically diversified library screening and isolation of donkey IgG binders

To initiate library screening, we chose to substitute the library with weakly electrophilic ncAA OBeY (**6**), with the intent of discovering antibodies that bind to and undergo spontaneous crosslinking with a target (Figure 3A). We focused initial screening efforts on the model target donkey IgG. In previous work, we identified binders to this target from a synthetic antibody library containing the same CDR-H3 design^30^, but with only canonical amino acids. Additionally, work from our lab has exploited the reactivity of the bromoethyl side chain of OBeY to promote spontaneous target crosslinking in high-affinity single-domain antibodies (sdAbs; also referred to as VHHs or nanobodies)^31^. Thus, we sought to identify screening conditions that would allow us to combine these two capabilities to attempt to isolate irreversible binders from the chemically diversified, billion-member library.

**Figure 3.**
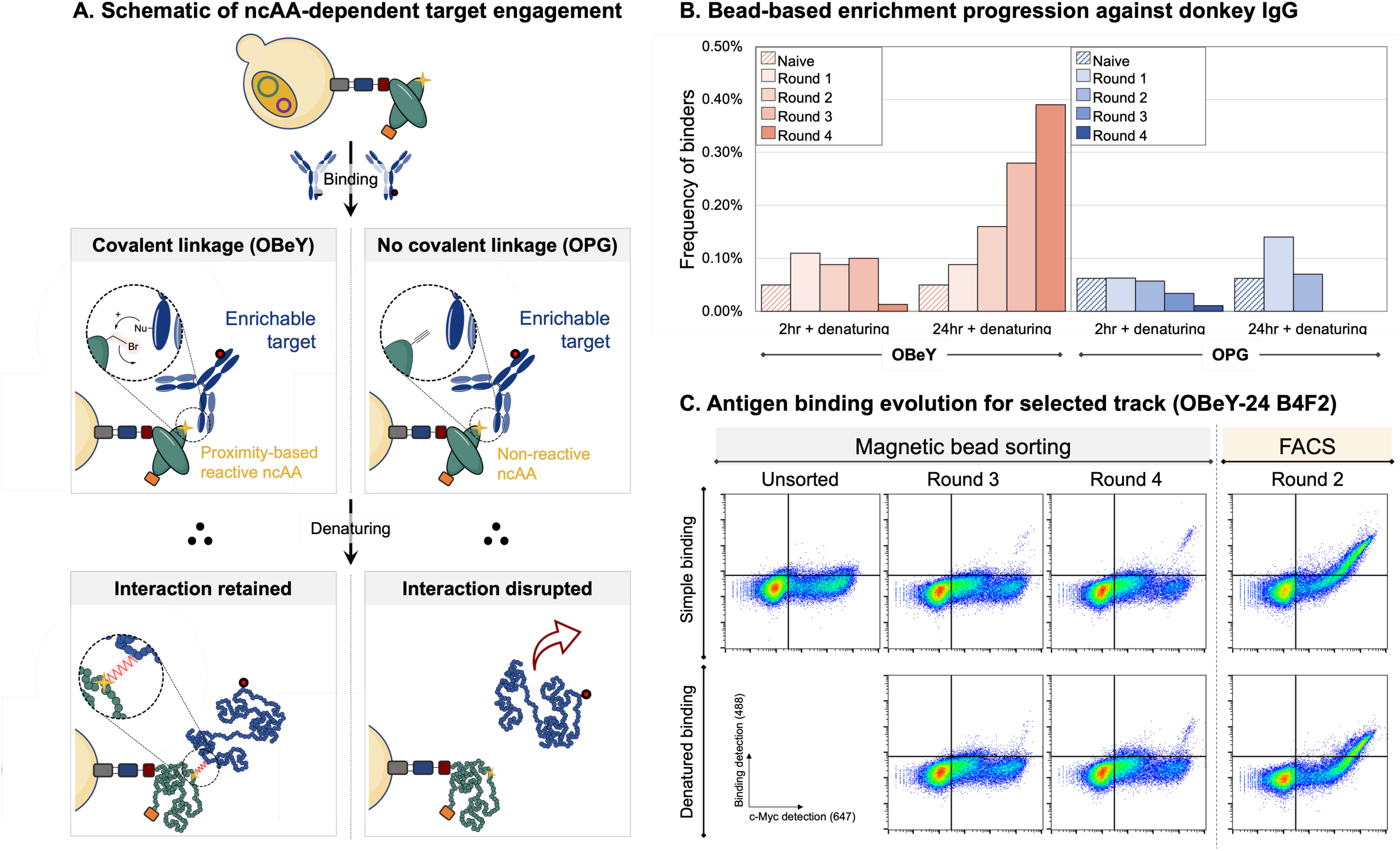
Overview of sorting progression for ncAA-containing binders to donkey IgG. A) Schematic concept of ncAA-dependent target engagement that was utilized for the sorts. B) Frequency of apparent binders to donkey IgG for four rounds of bead-based enrichments under two enrichment conditions with the OBeY-induced library and the OPG-induced library. C) Flow cytometry analysis of OBeY-substituted populations binding with donkey IgG with and without denaturing. Data was collected for a selected track (OBeY-24 B4F2) following 3 and 4 rounds of magnetic bead sorting and 2 rounds of FACS. For all cases simple binding conditions: 200 nM donkey IgG, PBSA washes; denatured binding conditions: 200 nM donkey IgG, denatured with 8 M urea, 200 mM EDTA, 25 mM tris, pH 8.4, PBSA washes.

We prepared versions of the library containing OBeY and OPG and subjected them to magnetic bead-based enrichments with antigen-coated beads (these enrichments followed an initial library depletion against non-specific binders; see *Materials and Methods* for details). While OBeY is known to undergo proximity-based crosslinking, the ncAA OPG in general does not. For both OBeY and OPG forms of the library, we used two conditions to attempt to promote enrichment of irreversible binders: (1) 2-hour antigen incubation at room temperature followed by 1 hour denaturing, (2) 24-hour antigen incubation at 4 °C followed by 1 hour denaturing. The denaturing step is intended to remove binders that that are noncovalently bound to the target and was conducted by subjecting cells to acidic conditions with 0.5 M tris-HCl at pH 4.1 for one hour (see *Materials and Methods* for details). Tris-HCl was selected as the denaturation buffer due to its efficiency in denaturing and promoting loss of antigen binding in control binders that contain only canonical amino acids (SI Figure 7).

Over four rounds of magnetic bead-based enrichments, we monitored enrichment progression using flow cytometry analysis (Figure 3B, C). Evaluation of each population revealed that for the OBeY-substituted library, a 24-hour incubation with bead-immobilized donkey IgG followed by denaturing (OBeY-24) resulted in a consistent increase in the frequency of apparent binders after each round of sorting (Figure 3B). After four rounds, the frequency of antigen-positive clones increased from 0.05% to 0.39%, diverging from the OPG control and the enriched populations that were subjected to 2-hour antigen incubations. Populations from the third and fourth round of bead-based enrichments were subsequently sorted via fluorescence activated cell sorting (FACS) using the same binding and denaturing conditions. After two rounds of FACS, the sorted populations were analyzed for retention of binding following denaturing (SI Figure 8). The population OBeY-24 B4F2 (subjected to four rounds of bead sorting followed by two rounds of FACS) was determined to be especially promising.

Single clones from OBeY-24 B4F2 were isolated, sequenced, and analyzed for binding via flow cytometry. Out of 13 sequenced clones, 8 are unique (Table 1). Given that sorts were conducted using biotinylated antigen, unique individual clones were evaluated for binding to the native, nonbiotinylated form of donkey IgG. Flow cytometry data indicated that all clones bind to nonbiotinylated IgG and that the level of antigen detection is dependent on antigen concentration (SI Figure 9, SI Figure 10). All clones possess a stop codon in the light chain, and one possesses an additional stop codon in the CDR-H3 sequence; this was consistent with the display phenotypes of the clones induced in the presence and absence of a ncAA as determined during flow cytometry analysis (SI Figure 11). The identification of binders with ncAA substitution sites only in the light chain, and no clones with ncAA substitution sites in the heavy chain (encoded within 38.2% of clones sequenced from the naïve library), suggests that the position of the ncAA may be playing a role in dictating which clones are enriched. Overall, this screening campaign demonstrated the ability to isolate unique binders to a target from a chemically expanded antibody library.

**Table 1.**
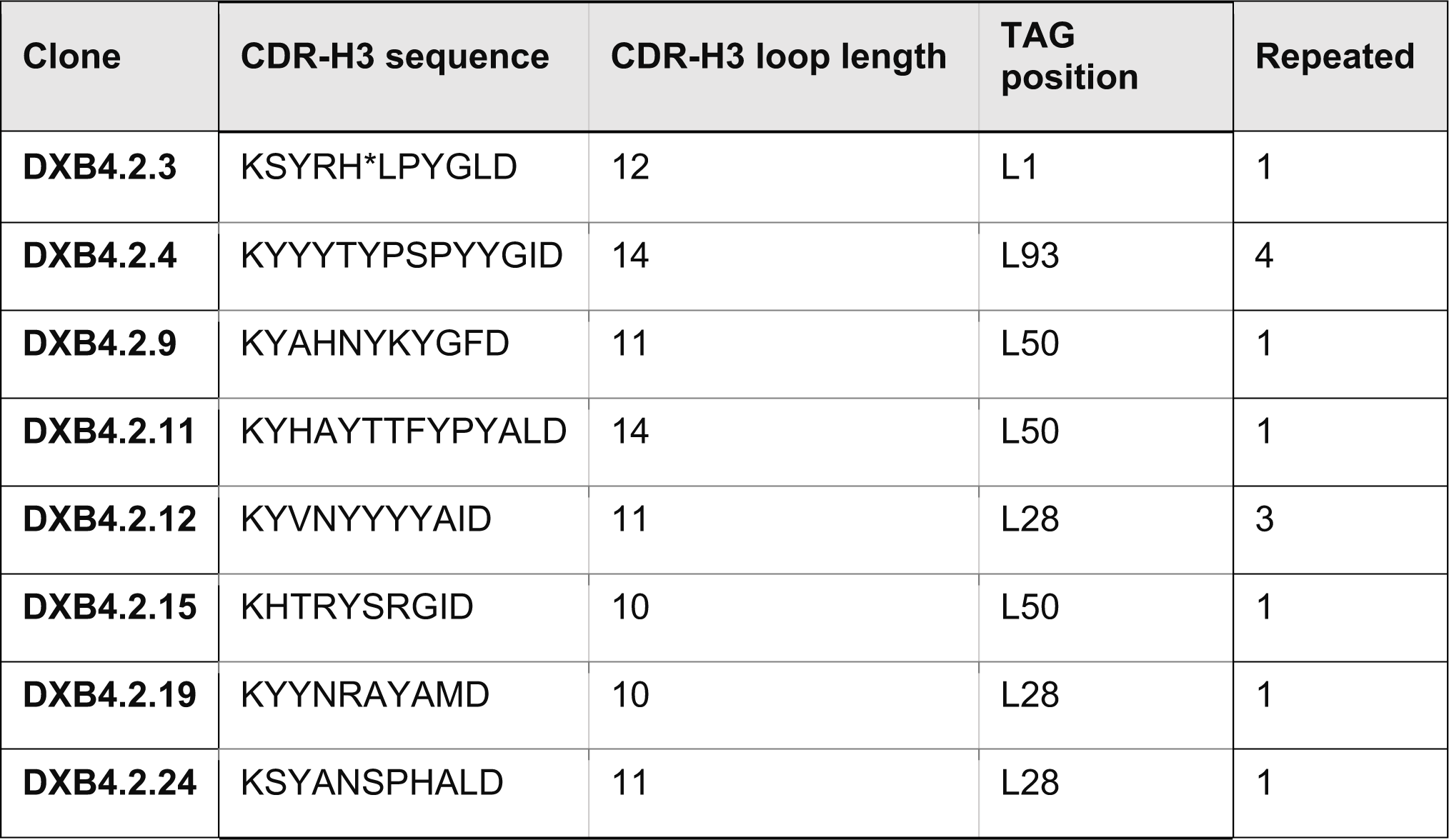
Key sequence features of donkey IgG-binding clones.

#### Characterization of donkey IgG binders on the yeast surface

Individual clones were evaluated further using flow cytometry assays to evaluate covalent antigen binding and binding affinity (Figure 4). To increase stringency during assays to evaluate possible covalent interactions, we used 8 M urea, 200 mM EDTA, 25 mM tris, pH 8.4 as the denaturant. This chemical denaturant is known to be stronger than the acid-based denaturant used during screening. Donkey1.1, a previously reported synthetic antibody clone containing only canonical amino acids^30^, was used as a noncovalent binding control to confirm that denaturation conditions led to the elimination of noncovalent binding interactions on the yeast surface (see also *Material and Methods*). We evaluated detection levels of donkey IgG binding for all OBeY- substituted and OPG-substituted single clones under binding and denaturing conditions alongside Donkey1.1 (Figure 4A, SI Figure 12). After denaturation, OBeY-substituted clones show antigen binding levels above control antigen detection levels, though these levels of detection are reduced in comparison to the levels detected in binding-only controls. This may be attributable to the low efficiency of spontaneous crosslinking of the bromoethyl side chain, which we have observed in previous work^31^. Controls in which OBeY was replaced by OPG or OmeY exhibit much lower donkey IgG detection levels after denaturation, with antigen detection levels similar to those of Donkey1.1 controls (SI Figure 13 and 14). This data provides indirect evidence that OBeY- substituted clones retain partial antigen binding following denaturation, which may point to crosslinking.

**Figure 4.**
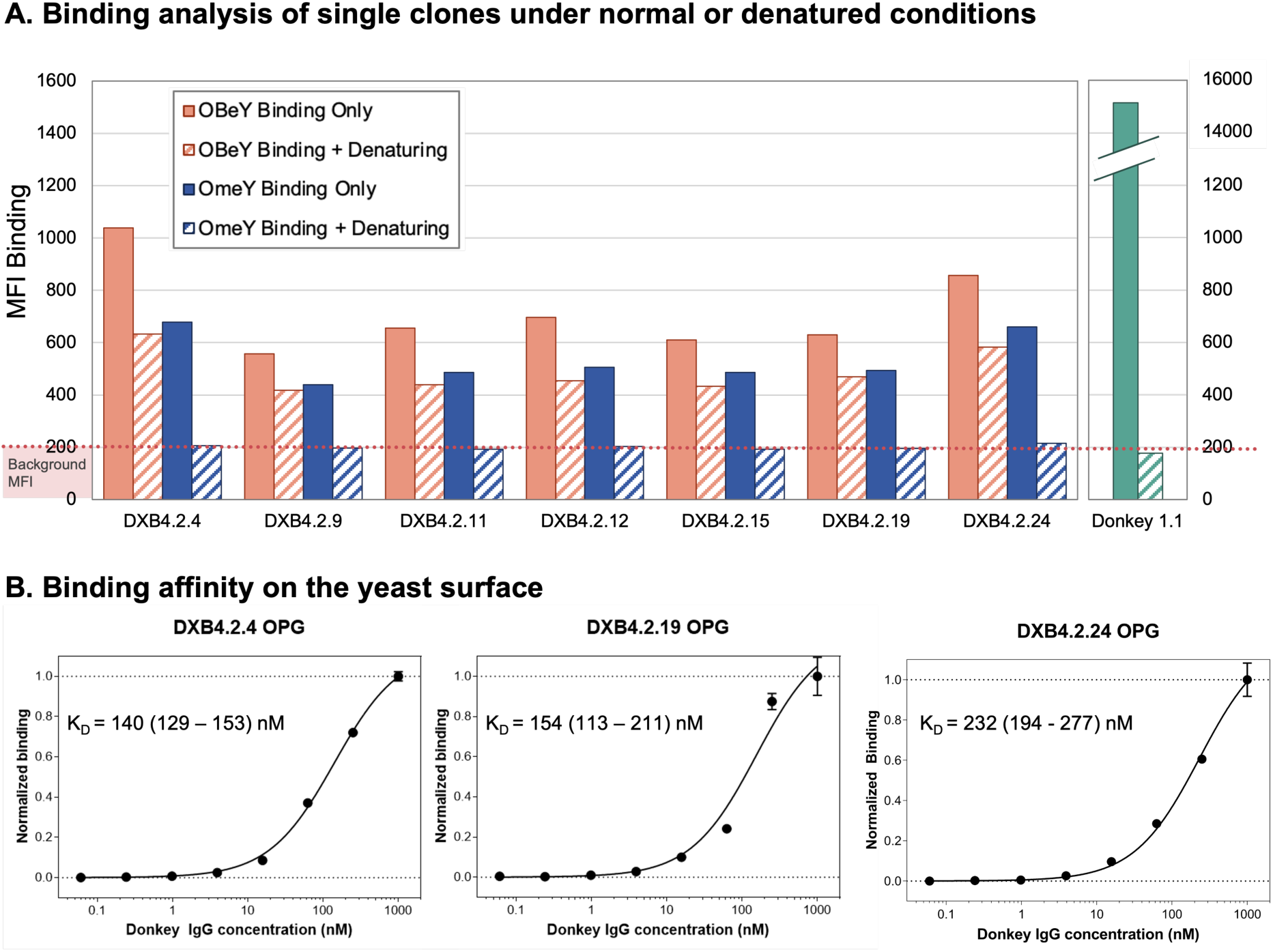
Binding and crosslinking analysis of isolated anti-donkey IgG single clones on the yeast surface. A) Donkey IgG binding of OBeY- and OPG-substituted samples under binding conditions with and without denaturation following binding. Binding only conditions: 200 nM donkey IgG (6 h at 37 °C), PBSA washes; binding + denaturing conditions: 200 nM donkey IgG (6 h at 37 °C), denatured with 8 M urea, 200 mM EDTA, 25 mM tris, pH 8.4 (24 hours), PBSA washes. B) Binding affinity evaluation of three individual clones on the yeast surface against donkey IgG.

We also investigated the binding affinities of three clones (DXB4.2.4, DXB4.2.19, DXB4.2.24). Since we observed potential evidence for crosslinking with OBeY-substituted clones, we used OPG-substituted versions of these clones to eliminate the possibility of crosslinking during K_D_ determination (equilibrium binding models assume reversible binding events). Yeast displaying OPG-substituted clones were treated with varying concentrations of donkey IgG and analyzed for antigen binding levels (Figure 4B and SI Figure 15). Apparent K_D_ values for all clones fall in the triple digit nanomolar range (Figure 4B) with DXB4.2.4 exhibiting a moderate K_D_ of 140 nM (95% confidence interval: 129-153 nM). This is approximately 5-fold lower in affinity than Donkey1.1 (27.9 nM; 95% confidence interval: 22.9-34.0 nM, SI Figure 16). However, in previous work, when Donkey1.1 was substituted with OPG or AzF at one of several positions, resulting variants exhibited apparent affinities predominantly in the triple digit nanomolar range as well^30^. This indicates that the modest affinities of newly isolated clones from the chemically expanded library are in line with the affinities of other ncAA-containing synthetic antibodies. Overall, data on single clones indicates that library screens have yielded binders against donkey IgG that exhibit moderate antigen affinity and improved antigen binding following denaturation when substituted with OBeY.

#### Characterization of donkey IgG binders in solution

Following characterization of single clones on the yeast surface, we set out to investigate the ability of isolated clones to bind and crosslink to donkey IgG in soluble form. We prepared scFvs of OBeY- and OPG- substituted DXB4.2.4 and DXB4.2.19 using secretion vectors described previously^47^ and His-tag purification with Ni-NTA resin. Following purification, SDS-PAGE analysis indicated that Ni-NTA-purified scFv constructs showed a single band at the expected molecular weight of ∼32 kDa (SI Figure 17). Since the DNA encoding these clones includes a TAG for ncAA incorporation via amber suppression, the detection of full-length protein is only possible with readthrough of the stop codon.

Once the full-length constructs were obtained and their purity was confirmed, we performed binding experiments with antigen-coated beads followed by denaturation with 8 M urea, 200 mM EDTA, 25 mM tris, pH 8.4 as before (SI Figure 18). Magnetic antigen-coated beads were incubated with 10× molar excess of OPG- or OBeY- substituted scFvs, and subjected to denaturation conditions (see *Materials and Methods*), after which samples were labelled for scFv and IgG detection and analyzed using flow cytometry to identify potential crosslinking (SI Figure 18). For all OBeY- substituted samples, the amount of scFv detected remained essentially unchanged regardless of whether the sample was subjected to denaturing conditions. In contrast, OPG-substituted samples exhibited a marked decrease in scFv binding following denaturation as evidenced by the steep drop in fluorescence (SI Figure 18). Control experiments in which IgG was directly detected after denaturation indicate no change in antigen detection levels, confirming that the donkey IgG linkage to the magnetic beads is not disrupted under the denaturing conditions used here. We note that OPG- substituted binders are not fully removed from the beads following denaturation; this is consistent with our observations on the yeast surface, and results from previous work in the lab^31^. In other studies, there is evidence that terminal alkynes can engage in covalent bond formations with cysteines in proximity-enhanced reactivity^48,49^; we cannot rule out this possibility in these experiments. Overall, in soluble form the isolated scFvs behave consistently with the yeast-displayed forms of the proteins.

### Chemically diversified library screening and isolation of PTP1B binders

To further evaluate the capabilities of the OBeY-substituted multi-modal library, we conducted screening against an additional, therapeutically relevant target: protein tyrosine phosphatase 1B (PTP1B, Figure 5A). PTP1B has been implicated in several diseases including cancer, diabetes, and obesity^50,51^. It is considered a difficult therapeutic target due to its sequence homology with other proteins in the PTP family^52^. Recently, multiple reports have described advances in targeting PTP1B^53–56^, including with OBeY substitutions in HER2 to identify proteins that covalently bind to PTP1B^57^. Thus, we sought to use the OBeY-substituted form of our library to discover antibodies that bind to, and perhaps covalently engage with, PTP1B (Figure 5).

**Figure 5.**
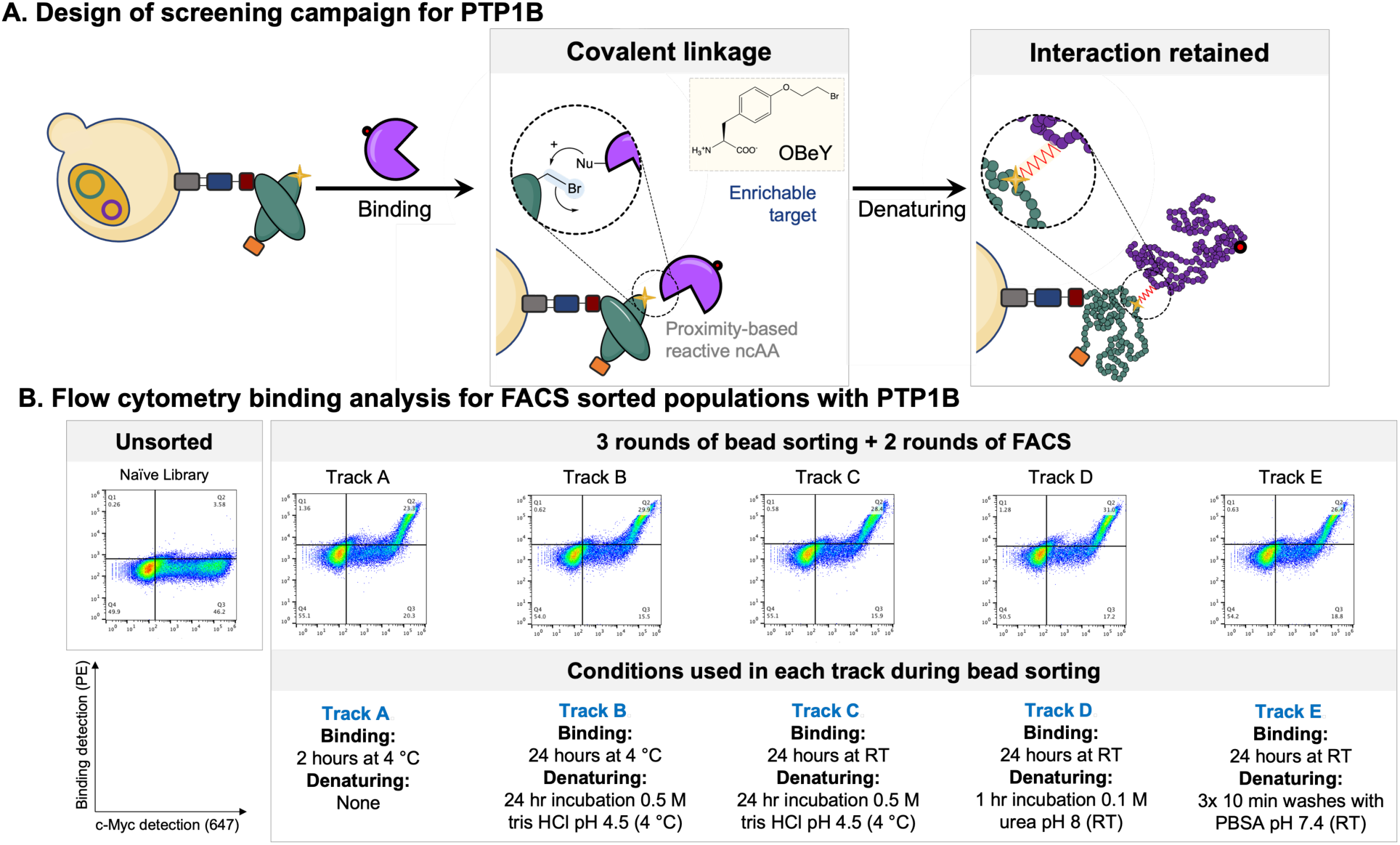
Library screening with PTP1B. A) Schematic of ncAA-dependent target engagement that was utilized for the sorts. B) Flow cytometry analysis of PTP1B binding following 3 rounds of bead sorting and 2 rounds of FACS for enrichment. Screening conditions for rounds 2 and 3 of bead sorting are listed below representative flow plots of the final enriched populations. Binding data shown was collected by incubating cells induced in the presence of OBeY with 200 nM PTP1B for 30 minutes at room temperature. Populations were subsequently labelled for biotin (PTP1B) detection as well as full length scFv detection (c-Myc).

We used several combinations of binding times, incubation temperatures, and denaturation conditions during screening against PTP1B (Figure 5B). A control track (Track A) used a set of conditions known to support enrichment of noncovalent binding proteins, and five additional tracks employed conditions of various stringency to attempt to promote the discovery of antibodies that spontaneously crosslink with PTP1B. For all tracks, the first bead-based enrichment (10 billion cells) served to enrich for clones binding to PTP1B irrespective of crosslinking. Following this initial enrichment, the recovered population was expanded, induced in the presence of OBeY, and subjected to parallel enrichments under varying levels of stringency. Flow cytometry analysis for PTP1B binding and for full-length display in the absence and presence of OBeY revealed populations exhibiting full-length display even in the absence of OBeY (SI Figure 19 and 20). This data suggests that populations lacking a TAG codon (here, called “wildtype”), though rare in the unsorted library, can be enriched. This may be due to the increased display (and thus avidity) of these clones in comparison to ncAA-containing clones. To isolate chemically diverse antibodies containing OBeY, we used FACS to remove any contaminating wildtype population (see *Materials and Methods*). Following depletion of wildtype populations from each enrichment track, we performed flow cytometry analysis on cells induced in the absence of ncAAs (SI Figure 21). This analysis shows that we effectively reduced the frequency of clones previously detected as full length in the absence of ncAAs down to 0.1%. We performed binding analysis with these populations induced in the presence of OBeY and OmeY, and we used denaturing and non-denaturing conditions to identify preferred conditions for future sorts (SI Figure 22). From this dataset, we observed that 24 hours of binding at 4 °C followed by 24 hours of denaturing leads to the most notable decrease in binding of the OmeY- substituted populations compared with the OBeY-containing populations, so we selected this condition in the subsequent two rounds of FACS. Analysis of the resulting enriched populations for PTP1B binding (Figure 5B and SI Figure 23) indicated high retention of binding in OBeY-substituted cells following denaturing as well as a notable decrease in binding with OmeY-substituted cells under the same conditions in both Tracks A and C. The data suggests these tracks are especially promising for isolating clones capable of covalent engagement, so we isolated individual clones from both Tracks A and C.

#### Characterization of PTP1B binders on the yeast surface

We identified six unique clones out of 38 clones isolated from Track A and C populations (Table 2). The sequencing data indicates that all clones possess ncAA incorporation sites in the light chain at positions L28 or L93. Consistent with the enrichments against donkey IgG (Table 1), the limited number of positions for ncAA incorporation in these clones suggests that the ncAA substitution site may play a role in driving enrichments. We next validated the binding properties of each clone with and without denaturing conditions at a variety of times and temperatures (Figure 6 and SI Figures 24-26). At each temperature (4 °C, room temperature, and 37 °C) and at the various time points, we detected high amounts of signal after incubation with 200 nM PTP1B using either OBeY or OmeY substitution. This data indicates that these clones are capable of interacting directly with PTP1B in yeast display format and support binding with multiple ncAA substitutions.

**Figure 6.**
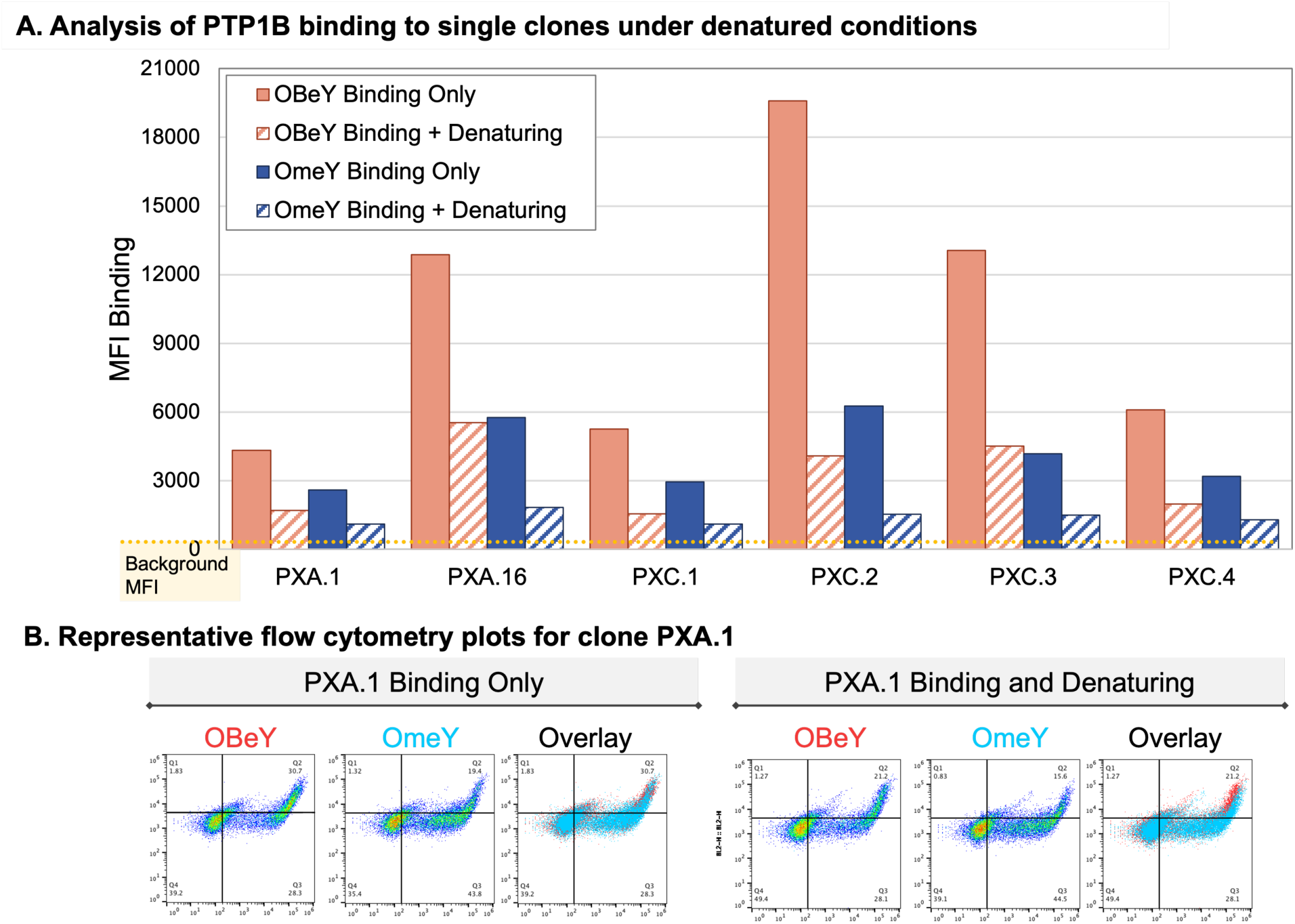
Binding analysis of isolated PTP1B-binding clones and crosslinking on the yeast surface. A) PTP1B binding of OBeY- and OmeY-substituted clones after binding with 200 nM PTP1B for 2 hours at 37 °C, with and without overnight denaturing with urea B) Representative flow cytometry plots and overlays from data shown in panel A of PXA.1.

**Table 2.**
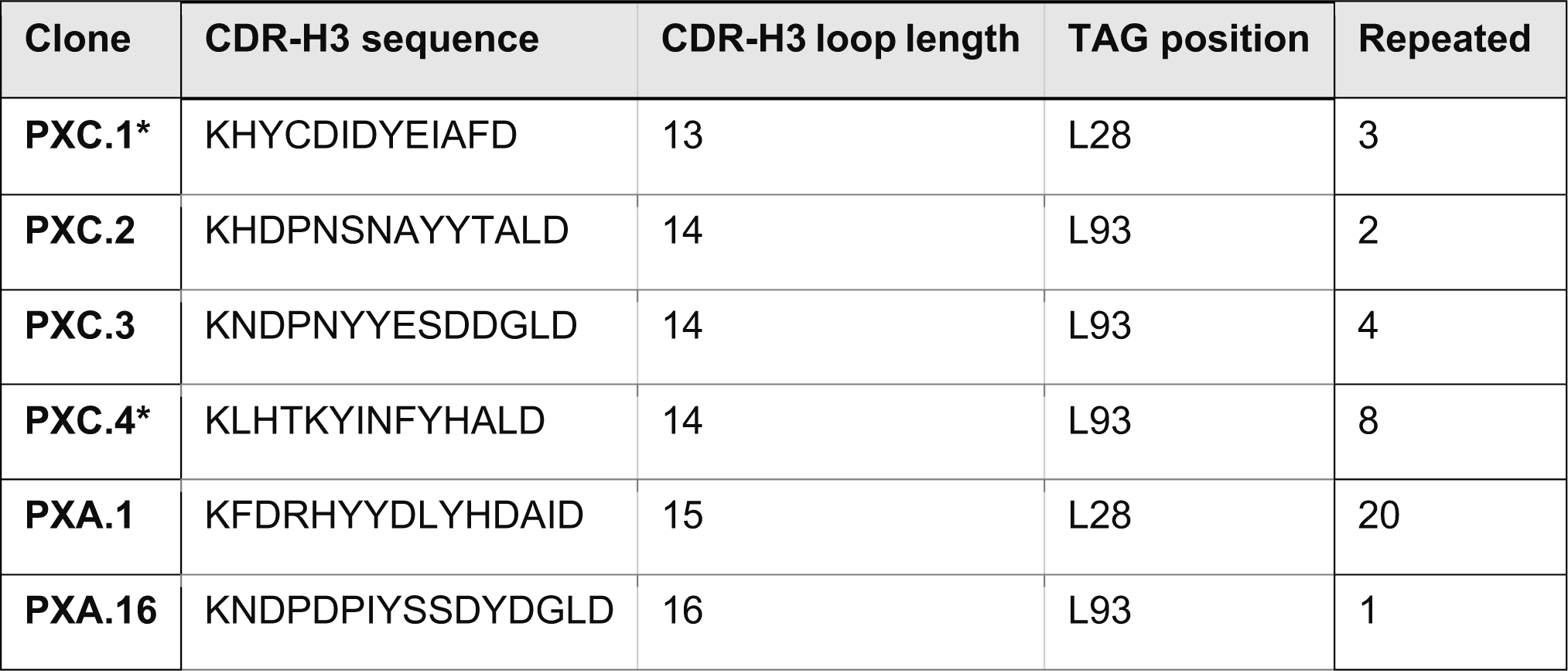
Key sequence features of PTP1B-binding clones.

To investigate whether covalent interactions with PTP1B are detectable, we performed a series of experiments with all 6 binding clones, where each was substituted with OBeY or the nonreactive analog OmeY. Over a range of incubation temperatures (4 °C, room temperature, and 37 °C) and incubation times (1–24 hours), followed by overnight denaturation (incubation with 8 M urea, 200 mM EDTA, 25 mM tris, pH 8.4), we were not able to identify binding conditions that simultaneously showed strong evidence for OBeY-substituted clones forming a covalent bond as well as full denaturing of OmeY-substituted clones (Figure 6, SI Figures 24-26). These experiments were partially confounded by what appeared to be nonspecific accumulation of PTP1B on the yeast surface even with cells displaying OmeY-substituted clones. Although the covalent binding experiments were not conclusive, the ncAA-substituted PTP1B binders we identified tolerate multiple ncAA substitutions, hinting at their potential chemical versatility.

#### Characterization of PTP1B binders in solution

To further investigate the binding properties of PTP1B binders in solution, we formatted clones PXA.1 and PXC.4 for secretion from yeast in scFv-Fc format^58^. We prepared these clones with both OBeY and OmeY substitutions and purified the proteins from culture media using Protein A resin. SDS-PAGE analysis confirmed the isolation of pure scFv-Fcs (SI Figure 27). The banding patterns observed are consistent with those in our prior preparations of scFv-Fcs and related Fc fusions from yeast^29,30,58,59^.

With the purified scFv-Fcs, we conducted bead-based assays to evaluate binding to PTP1B. Biotinylated PTP1B was coated on magnetic streptavidin beads, and aliquots of beads were incubated with 5-fold molar excess of PXA.1-OBeY, PXA.1-OmeY, PXC.4-OBeY, PXC.4-OmeY, or a nonbinding control (FAP-Fc^59^, a previously reported fibroblast-activation protein binder) for 3 hours at 4 °C. Following incubation, beads were analyzed for PTP1B and scFv-Fc detection via flow cytometry. We detected PTP1B in all samples, and scFv-Fcs were detected in each sample incubated with PTP1B-binding scFv-Fcs (SI Figure 28), indicating that PXA.1 and PXC.4 clones are capable of binding in solution as well as on the yeast surface when substituted with either with OBeY or OmeY.

Next, we tested the scFv-Fc binders for inhibition of the enzymatic activity of PTP1B. Our rationale was that OBeY may be able to interact with amino acid side chains in the catalytic active site of PTP1B and disrupt its activity. We prepared binding reactions using a 20-fold molar excess of scFv-Fcs and allowed the binding to proceed for 30 minutes at room temperature before assaying for activity. Measurement of the increase in signal corresponding to substrate hydrolysis by PTP1B (see *Materials and Methods*) was indistinguishable in samples containing PTP1B alone versus samples preincubated with high concentrations of scFv-Fcs (SI Figure 29). This outcome is unsurprising, as inhibitory antibody discovery via yeast display is extremely challenging^47,60^. Nonetheless, our confirmation of binding activities of PTP1B-binding clones in yeast display and soluble scFv-Fc formats further validates the concept of screening a library of chemically expanded antibodies. These initial studies highlight a number of opportunities for future work described below.

### Conclusions

In this work, we have established a chemically diversified antibody library in yeast display format and demonstrated the feasibility of conducting antibody discovery with this unique collection. The multi-modal library reported here contains greater than one billion synthetic antibody clones that can be prepared to present a broad range of chemistries through either different ncAA side chains or via bioorthogonal click chemistry conjugations. We can directly encode chemical groups with photo-reactive, clickable, proximity-based reactive, and additional chemical properties; CuAAC reactions with azides or alkynes substantially expands the range of accessible chemistries. Here, we conducted screens against donkey IgG and PTP1B. For both of these targets, we were able to isolate OBeY-substituted clones that bind to the target of interest. In addition, for the case of donkey IgG, parallel screens of the library in OBeY- substituted and OPG-substituted formats led to different enrichment outcomes. OBeY- substituted populations could be enriched for increasing fractions of binding clones following incubation under denaturing conditions, whereas OPG-substituted populations could not. We identified eight distinct clones that all exhibited binding to the native form of donkey-IgG and showed evidence for OBeY-dependent retention of antigen binding following denaturation on the yeast surface. Further experimentation on two clones prepared in soluble form led to outcomes that are consistent with what was observed on the yeast surface. However, we also note that we could not unambiguously observe crosslinking in solution. For PTP1B, we identified six unique clones using a wider range of screening conditions than for donkey IgG. In addition, we established procedures for eliminating clones that only contain canonical amino acids following initial sets of enrichments. The construction of this unique library, demonstrations of diverse chemistries presented therein, and clear identification of binding clones in library screens are important advances for chemically expanded antibody discovery.

We also note some limitations and the need for additional work to further advance this discovery platform. Although our lab has previously been successful in identifying spontaneously crosslinkable single-domain antibodies containing OBeY starting from existing binders^31^, we encountered some challenges in our *de novo* covalent antibody discovery efforts. We attribute some of these difficulties to the combination of OBeY’s low reactivity and the modest binding affinities of synthetic antibody clones present within this simple synthetic antibody library design (in contrast to the subnanomolar sdAbs used in other work^31^). These two factors may hinder crosslink formation, confounding our attempts to enrich and detect crosslinking events. Another challenge is that we detected antigen binding above background following denaturation in the case of OPG- and OmeY-substituted clones on the yeast surface (with donkey IgG and PTP1B binders) and on beads (for donkey IgG binders). Elevated detection even after extended incubation with 8 M urea buffer indicates aggregates may be retained on yeast and bead surfaces nonspecifically. An improved understand of what leads to these aggregates is likely to improve the performance of covalent protein discovery.

Chemically expanded antibody discovery offers numerous opportunities for powerful discovery in future screening campaigns. In the case of spontaneously crosslinkable proteins, the mechanisms dictating crosslinking efficiency and specificity remain poorly understood^31,61^. High-throughput screening has the potential to yield the depth and breadth of data needed to establish the fundamental principles underlying in on- and off-target crosslinking events. Beyond OBeY and related haloalkanes, crosslinking functionalities including fluorosulfonates and acrylamides, which are known to support improved reaction kinetics^62^, have the potential to enhance covalent antibody discovery. Beyond spontaneous crosslinking, there are numerous opportunities to conduct chemistry-driven antibody enrichments. Current studies in our laboratory with the library described here include investigations of photocrosslinkable antibody discovery and pharmacophore-driven antibody discovery. Beyond this particular library, there are opportunities to enhance chemically expanded antibody diversities further. The introduction of amino acid diversification only within the CDR-H3 region is likely limiting the affinities of clones encoded in the library (consistent with our previously reported simple synthetic antibody library containing only canonical amino acids^30^). Another limitation of the chemically expanded antibody diversity described here is that ncAAs are presented within minor CDRs that are otherwise fixed in amino acid sequence. Given the potential importance of local sequence context, library designs that present ncAAs within diversified binding loops is an important area for future library design. Finally, display levels with our current library are sufficiently high to support sorting, but ncAA incorporation efficiency with AcFRS is relatively low in comparison to other available aaRS variants in our laboratory^37,63^. More efficient ncAA incorporation may reduce the probability of enriching for clones lacking stop codons, as we observed when screening against PTP1B in this work. All of these considerations, and perhaps more, should be weighed carefully during the design and preparation of second-generation, chemically diversified antibody libraries in yeast display format. Overall, this study introduces a versatile platform capable of integrating a wide range of chemical functionalities into antibodies, facilitating high-throughput screening against desired targets. This multi-modal platform offers access to unique discovery spaces with the potential identify novel antibody-based reagents, diagnostics, and therapeutic leads.

## Materials and Methods

### Materials

The RJY100 strain *Saccharomyces cerevisiae* utilized in this work was constructed as described previously^64^. The noncanonical amino acids *p*-azido-L-phenylalanine (AzF) and *O*-methyl-L-tyrosine (OmeY) were purchased from Chem-Impex International, Inc., *p*-propargyloxy-L-phenylalanine (OPG) was purchased from Iris Biotech GmbH, *p*-acetyl-L-phenylalanine (AcF) was purchased from SynChem, Inc., 4-iodo-L-phenylalanine (IPhe) was purchased from Astatech, Inc., and *O*-(2- bromoethyl)tyrosine (OBeY) was purchased from Advanced ChemBlocks, Inc. Small molecules used in click chemistry reactions were purchased from the following locations: Chemspace US Inc. (N-hydroxy-10-undecynamide and N-(2-azidoethyl)-[2,2’- bipyridine]-5-carboxamide), Alfa Aesar (10-undecynoic acid), Sigma Aldrich (2-(2- propyn-1-yloxy)ethane-sulfonyl fluoride), Ambeed Inc. (4-ethynylbenzene-1- sulfonamide), Discovery Molecules (4-(dihydroxy-borophenyl) acetylene), and 1Click Chemistry Inc. (4-(propargyl-aminocarbonyl)phenylboronic acid). N_3_-PEG_4_-VS and N_3_- bAGbA-VS were generously gifted by the Bogyo Lab (Stanford University). Restriction enzymes used for cloning were purchased from New England Biolabs. Primary antibodies for flow cytometry labeling used in this work include chicken anti-cMyc from Exalpha, and mouse anti-HA from BioLegend. Secondary antibodies for flow cytometry labeling used in this work include goat anti-chicken Alexa Fluor 647, goat anti-mouse Alexa Fluor 488, streptavidin Alexa Fluor 488, rabbit anti-donkey DyLight 488 purchased from Thermo Fisher Scientific, and mouse anti-biotin PE from Biolegend.

New England Biolabs Q5 DNA polymerase was used for PCR amplifications. Synthetic oligonucleotide primers for cloning and sequencing were purchased from Eurofins Genomics, IDT DNA Technologies, or Azenta Life Sciences (formerly GENEWIZ). gBlocks encoding heavy and light chain frameworks were purchased from IDT DNA Technologies. All DNA sequencing was performed by Quintara Biosciences. Competent *E. coli* was prepared with Mix and Go! kits from Zymo Research. Plasmid DNA isolation from E. coli was done with Epoch Life Science GenCatch^TM^ Plasmid DNA Mini-Prep Kits. Yeast transformation and plasmid DNA isolation were done with Frozen-EZ Yeast Transformation II kits and Zymoprep DNA isolation kit from Zymo Research. Magnetic bead sorting was conducted with Dynabeads^TM^ Biotin Binder from Thermo Fisher Scientific. Penicillin-Streptomycin used was Corning brand (100× concentration; “pen-strep”). Antibodies used in this work were Chrompure Donkey IgG whole molecule (unconjugated and Biotin-SP conjugated forms) from Jackson ImmunoResearch. Reagents for antigen biotinylation: EZ-Link^TM^ NHS-LC-Biotin, Zeba^TM^ Spin Desalting Columns (7 kDa MWCO, 0.5 mL), and Pierce^TM^ Biotin Quantitation Kits were purchased from ThermoFisher Scientific. Reagents for CuAAC (copper-catalyzed azide-alkyne cycloaddition): Biotin-PEG_3_-Azide, Biotin-PEG_4_-Alkyne, and THPTA were purchased from Click Chemistry Tools; (+)-Sodium L-ascorbate, aminoguanidine hydrochloride, copper sulfate pentahydrate, EDTA, dimethylformamide, and DMSO were purchased from Sigma Aldrich. SDS-PAGE analysis was done with 4-12% Bis-Tris mini gels and SimplyBlue Safestain.

### Plasmid design

We based our chemically expanded synthetic antibody library on our previously described library^30^. The sequence of a single chain variable fragment (scFv) is encoded within pCTCON2 between the NheI and BamHI cut sites. The two regions of the scFv are linked by a peptide linker identical to the linker sequence used in Van Deventer et al^27^. The frameworks are described in detail in Islam and Kehoe et al^30^. To position stop (TAG) codons in the desired positions, primers were designed with an overlap of ∼30 nucleotides on each end of the stop codon. Six different primers with TAG codons replacing natural codons were designed for positions L1, L28, L50, L93, H31 and H54 (SI Table 2). As previously described, to mimic CDR sequences in natural antibodies, oligonucleotides with custom nucleotide frequencies and different lengths were designed and used to amplify the heavy chain of the antibody fragment^30^. Germline sequences for framework regions can be found in SI Table 3. Loop lengths of the diversified CDR-H3 region vary in length between 9 and 17 amino acids.

The orthogonal translation system consisting of a tRNA_CUA_ and *p*-acetyl-L-phenylalanyl tRNA synthetase (AcFRS) has previously been prepared in the plasmid pRS315-KanRmod-AcFRS, which encodes a kanamycin antibiotic resistance marker^30^.

### Media and reagent preparation

The RJY100 strain of *Saccharomyces cerevisiae*, used here as the host cell strain for library construction, was previously made using standard homologous recombination approaches^65^. Solid and liquid media including LB, YPD, and YPG were prepared using procedures described previously^65^. SD-SCAA and SG-SCAA (–Trp –Leu –Ura and –Leu –Ura) were prepared as described previously^37^. Procedures for preparation of electrocompetent yeast followed previously detailed protocols^65^. 8 M urea (pH 8.4, EDTA 200 mM, tris 25 mM), PBSA (pH 7.4, NaCl 137 mM, KCl 2.7 mM, Na_2_HPO_4_ 10 mM, KH_2_PO_4_ 18 mM, 0.1% w/v BSA), PBS (pH 7.4, NaCl 137 mM, KCl 2.7 mM, Na_2_HPO_4_ 10 mM, KH_2_PO_4_ 18 mM) were prepared and sterile filtered prior to use.

Terrific Broth (TB) used to produce soluble PTP1B antigen was produced by combining 24 g yeast extract and 20 g tryptone with 900 mL DI water and 4 mL glycerol. Mixture was sterilized by autoclaving. A solution of 0.17 M KH_2_PO_4_, 0.72 M K_2_HPO_4_ was prepared and sterile filtered. 100 mL phosphate solution was added to autoclaved portion of media for 1 L final volume and final concentrations of media components as follows: 0.017 M KH_2_PO_4_, 0.072 M K_2_HPO_4_, 24 g/L yeast extract, 20 g/L tryptone, and 4 g/L glycerol.

### Library construction

#### PCR Amplification of DNA fragments

The library was assembled in several steps. The gBlock encoding the first region was composed of the IGKV1-39 light chain and the JK4 J-segment. The light chain DNA sequence encoding the entire amino acid sequence of IGKV1-39, followed by the “TFGGGTKVEIK” sequence from JK4 was used as template. To amplify the light chain of the scFv region without stop codons, the primers SyntheticAmpFwd, which has a pCTCON2 overlap region, and SidLinkRev were used in a PCR reaction (SI Table 2). To amplify the light chain of the scFv region with stop codons, the primers SyntheticAmpFwd and L[*position*]TAGRev were used for the 5’ to TAG, while the primers L[*position*]TAGFwd and SidLinkRev were used for the TAG to linker (SI Table 2). The gBlock encoding the second region contained the sequence for the IGHV3-23 VH region and was used as template. To amplify the heavy chain of the scFv region without stop codons the primers SidLinkFwd and one of the primers CDR9-17Rev were used (SI Table 2). The CDR9-17Rev primers contain an overhang for the first part of the JH4 sequence. The primer SyntheticAmpRev was used to incorporate the rest of the JH4 sequence and the pCTCON2 overlap region, producing the Linker to 3’ insert. To amplify the heavy chain of the scFv region with stop codons the primers SidLinkFwd and H[*position*]TAGRev were used to obtain the linker to TAG insert, while the primers H[*position*]TAGFwd and CDR9-17Rev were used for the TAG to CDR region, which was then amplified using the SyntheticAmpRev primer to incorporate the rest of the J-segment and pCTCON2 overlap region to obtain the TAG to 3’ insert.

Amplified DNA fragments were purified via gel electrophoresis, with gels imaged using an Azure c400 gel imager. The DNA was then extracted and purified, and concentrations were measured via NanoDrop^TM^ Microvolume UV-Vis Spectrophotometer (ThermoFisher Scientific). DNA was stored in DI water at –20 °C until used further.

To digest the plasmid pCTCON2, which served as the vector backbone for the scFv assembly, the enzymes SalI, BamHI, and NheI were used for linearization along with a method described previously^65^.

#### Chemical yeast transformation

The suppression machinery-containing plasmid pRS315-KanRmod-AcFRS (Leu marker) was transformed into chemically competent RJY100 cells, plated on SD-SCAA –Leu –Ura solid media, and grown for ∼3 days at 30 °C until colonies appeared. Colonies were inoculated in 5 mL liquid media (SD-SCAA –Leu –Ura) supplemented with pen-strep at 1:100 dilution. Cells were grown at 30 °C with shaking at 300 RPM until saturation (∼3 days). Liquid cultures of cells containing the suppression machinery plasmid were propagated in 5 mL fresh liquid media supplemented with pen-strep in preparation of making cells electrocompetent.

#### Yeast electroporation

Cells transformed with the plasmid encoding the suppression machinery were made electrocompetent following methods described in detail elsewhere^42,65^. 4 µg of each of the two purified PCR inserts (light chain fragment and heavy chain fragment) and 1 µg of digested vector pCTCON2 were mixed together with 1 µg of the AcFRS plasmid and transformed into yeast via electroporation as described elsewhere^42,65^. The library encoded within the display plasmid was assembled by homologous recombination in electrocompetent RJY100. Electroporations were performed while keeping PCR fragments with different TAG positions and CDR-H3 loop lengths separate from one another to create 54 distinct sublibraries (6 TAG positions with 9 loop lengths). Each transformed culture was recovered in 100 mL SD-SCAA –Trp –Leu –Ura + pen-strep. A small portion of the transformed cells were plated on SD-SCAA –Trp –Leu –Ura plates in serial dilutions and allowed to grow for ∼3 days at 30 °C until colonies were formed. Colonies were counted to determine the transformation efficiency of each sublibrary. Upon saturation in liquid minimal media, each sublibrary was propagated into 1 L cultures and grown at 30 °C with shaking at 300 RPM until saturation. Saturated cultures were then pelleted and resuspended to a final concentration of 15% glycerol and stored in 2 mL aliquots at –80 °C each containing 10^10^ cells.

### Preparing noncanonical amino acid liquid stocks

All ncAAs were prepared as working stocks at 50 mM concentration. Stocks of sterile filtered 50 mM concentration of the L-isomer of ncAAs were prepared for each experiment. DI water was added to dissolve the ncAA powder with mixing by vortexing and some ncAAs required titrations with 6 N sodium hydroxide to dissolve fully. Dissolved solutions were sterile filtered through 0.2 µm filters and used immediately or stored at 4 °C for up to one week. Photosensitive ncAAs such as AzF were covered in aluminum foil and stored in the dark. In the case of *O*-methyl-L-tyrosine (OmeY), the solution required the addition of NaOH to dissolve it, and then the solution was then made pH neutral with the addition of hydrochloric acid (HCl). The solutions were then sterile filtered through a 0.2 µm filter and were used immediately or stored at 4 °C for up to one week.

### Yeast induction in preparation for flow cytometry

To propagate yeast cultures and prepare them for induction, cells were either thawed from glycerol stocks stored at –80 °C or passaged from fridge cultures. Cultures propagated in liquid SD-SCAA –Trp –Leu –Ura media to saturation overnight. Following saturation, the cultures were diluted to an OD_600_ of 1 in fresh media and grown at 30 °C until mid-log phase (OD_600_ of 2-5), approximately 4-8 hours. Cells were pelleted and resuspended to an OD_600_ of 1 in induction media (SG-SCAA –Trp –Leu –Ura). For site-specific incorporation, ncAAs were added to the media at 1 mM final concentration, and samples were incubated at 20 °C with shaking for 16-24 hours.

### Flow cytometry data collection and analysis

Induced samples, prepared as above, were labelled in 1.7 mL Eppendorf microcentrifuge tubes or V-bottom 96-well plates. 2×10^6^ cells per sample were transferred to microcentrifuge tubes and pelleted for 30 s at 14,000 rcf (or 5 min at 2,400 g). The samples were washed three times in 1 mL PBSA (or 200 µL for plates). Primary antibody labelling was conducted at room temperature for 30 min on a rotary wheel (or on an orbital shaker at 150 RPM for plates). Cells were pelleted with all remaining steps being performed on ice and with ice-cold PBSA. Cells were washed twice, and secondary labelling was performed for 15 min on ice in the dark. Following secondary labelling, the cells were washed once with ice-cold PBSA and left on ice as pellets until ready to analyze in the cytometer. Immediately prior to running each sample, the cells were resuspended in 1 mL ice-cold PBSA (or 200 µL for plates). Analytical flow cytometry was performed on an Attune NxT flow cytometer in the Tufts University Science and Technology Center using methods described in previous work^37,42^. Labelling reagents, concentrations, and volumes used for primary and secondary labelling were dependent on the type of experiment and are listed in SI Table 4. All labelling reagents were prepared in PBSA unless otherwise noted. Results from flow cytometry experiments were analyzed using FlowJo v10.8.1 software.

For flow cytometry characterization of sublibraries, the data analysis process involved several steps using FlowJo and Microsoft Excel. The general procedure followed these steps:

1. Gating: The data was initially gated to isolate the single cell populations, ensuring accurate analysis of individual cells.

a. Truncated Population Gating: The populations that were induced without a ncAA were gated to isolate the induced populations which corresponded to display events with positive HA detection only. The gate was applied to all populations.
b. Full-length Population Gating: An additional gate was drawn to isolate the induced populations that displayed full-length clones, which corresponded to display events with positive HA and c-Myc detection. The gate was applied to all populations.
2. Retrieval of frequency data: Data for the frequency of clones in each gate was extracted for HA and c-Myc detection for induced populations in the absence and presence of a ncAA. This allowed for the quantification of clones lacking TAG codons and truncation levels.
3. Data analysis in Excel: Microsoft Excel was used to calculate the percent full-length clones vs overall displayed in the absence of a ncAA (SI Figure 2). Truncation levels were calculated by the percent truncated clones vs overall displayed in the presence of a ncAA.

### DNA sequencing

Each individual sublibrary was propagated, and plasmid DNA was extracted using the Zymoprep DNA isolation kit. The DNA was transformed into *Escherichia coli* and plated in LB + 50 µg ampicillin plates. Plates were incubated at 37 °C for 16 hours for colonies to grow. Single *E. coli* colonies were inoculated, and sequence verified by Sanger sequencing using sequencing primers CON2seqFwd and CON2seqRev (SI Table 2). All sequencing in this work was submitted to Quintara Biosciences (Cambridge, MA). The diversification scheme as well as presence of the TAG codon at the intended locations were analyzed. Six colonies were sent for sequencing from each sublibrary, and a total of 317 clones were sequenced. The sequences obtained for all the individual sublibraries were analyzed using Geneious v11.1.1. Sequences were aligned, extracted, and exported into Excel for further analysis (SI Table 1).

### Library pooling

To pool the sublibraries to create the billion-member library, sequence-verified sublibraries were propagated from the freezer stocks into 100 SD-SCAA –Trp –Leu – Ura + pen-strep at 30 °C. To fully cover library diversity, 10× the amount of transformants was pooled from each sublibrary into the same 1 L culture, and in turn the pooled library was propagated to saturation. Sublibraries that twice did not propagate well after thawing were excluded from the final library. Upon saturation, 10× the number of transformants (summed from the number of transformants of each sublibrary) were spun down at 2,000 rcf for 5 min and resuspended in 8 L SD-SCAA –Trp –Leu –Ura + pen-strep. Cultures were grown to saturation overnight. All saturated cultures were spun down, resuspended in 15% glycerol, and stored in 2 aliquots as before at –80 °C with each aliquot containing ∼10^10^ cells. A final characterization of the full-length display and truncation in the pooled library following induction in the presence of each of six ncAAs was done using flow cytometry (SI Figure 6).

### Bead Depletion

Prior to conducting enrichments with the library, the pooled library was depleted for binders to bare beads and biotin-streptavidin complexes. Biotin Binder Dynabeads® (ThermoFisher Scientific) were washed and prepared as described previously^65^. Bare beads were prepared by washing 150 µL beads on a DynaMag-2 magnet in 1 mL PBSA twice. Biotin-coated beads were prepared by incubating 50 µL washed bare beads with 1 µL 100 mM D-biotin on a rotary wheel for 2 hours at 4 °C (final biotin concentration 100 µM). Following incubation, beads were washed twice on a DynaMag-2 magnet in 1 mL ice-cold PBSA.

Two vials of glycerol stocks, covering 20× the pooled library diversity, were thawed and propagated in 1 L SD-SCAA –Trp –Leu –Ura + pen-strep to saturation. Yeast was then diluted and grown until mid-log phase as described above. The culture was subsequently induced in the presence of 1 mM AzF at 20 °C with shaking at 300 RPM for 16 hours in the dark. Following induction, 10×10^9^ cells were spun down in conical tubes for 5 min at 2,400 g at room temperature and washed three times in 50 mL PBSA. The pellets were subsequently resuspended in 15 mL ice-cold PBSA. 50 µL Biotin Binder beads were added to the cells and incubated on a rotary wheel at 4 °C for two hours. Following this first bead incubation, cells were placed on a magnet (DynaMag-15) for 2 min. Cells were then removed from the tube with a serological pipette, without moving the beads and were placed in a new tube, and the process was repeated twice more. After a total of 3 incubations on the magnet, the first depletion was complete. Another 50 µL bare beads was added to the cells and they were incubated on a rotary wheel at 4 °C for two hours. Following this second bead incubation, cells were once again depleted as described above. Finally, 50 µL Biotin Binder beads that had been treated with biotin (incubated with 1 µL 100 mM D-biotin, final concentration 100 µM, for 2 h at 4 °C on a rotary wheel) were added to the cells, and they were incubated on a rotary wheel at 4 °C for two hours. Following this third bead incubation, cells were again depleted as above. Depleted cells were propagated to saturation, and frozen stocks were prepared in 15% glycerol and stored at –80 °C.

### CuAAC click chemistry

All small-scale click chemistry reactions were performed in microcentrifuge tubes using previously determined reaction conditions^42,46^. Biotin-alkyne or biotin-azide was dissolved in DMSO to a 100 mM concentration and diluted appropriately to 20 mM stock concentrations for reactions. CuAAC was performed using the general protocol of Hong et al. (2009)^66^. Except for the small molecule concentrations, all other aspects of the protocol were followed, including the order of reagent addition. Cell samples freshly induced in the presence of either AzF or OPG were pelleted and washed thrice with PBSA prior to a final resuspension in PBS before the reactions. For the two-step detection of reaction products, CuAAC reactions were performed with the alkyne- or azide- containing molecule of interest at a final 1 mM concentration for 4 h, followed by washing three times in ice-cold PBSA and resuspension in PBS. A second reaction using the previously mentioned biotin-alkyne or -azide reagent was performed on the once-clicked samples.

#### Small scale reactions (2×106 cells in 250 μL)

First step of the reaction was CuAAC with a small molecule. Cells were resuspended in 220 μL 1× PBS. The following volumes were added to the resuspended cells for the reaction in order with vortexing between each addition:

i. 2.5 μL 100 mM cargo-azide/alkyne (final concentration 1 mM)
ii. 3.8 μL 1:2 volume mixture of 20 nM CuSO_4_: 50 mM THPTA.
iii. 12.5 μL 100 mM aminoguanidine.
iv. 12.5 μL 100 mM sodium ascorbate

The reaction was left running for 4 h before termination. Termination of reaction was achieved by washing cells three times with PBSA. Following washes, cells were resuspended in 220 μL 1× PBS. Biotin click reaction was conducted by adding the following volumes to the cells for the reaction in order with vortexing between each addition:

i. 1.25 μL 20 mM biotin-azide/alkyne (final concentration 100 µM)
ii. 3.8 μL 1:2 volume mixture of 20 nM CuSO_4_: 50 mM THPTA
iii. 12.5 μL 100 mM aminoguanidine
iv. 12.5 μL 100 mM sodium ascorbate

The reaction was allowed to proceed for 15 min. Termination of reaction was achieved by washing cells three times with PBSA. Following washes, cells were labelled for flow cytometry detection of biotin and full-length display (SI Table 4).

#### Large scale reaction (1×10^10^ cells in 50 mL)

First step of the reaction was CuAAC with small molecule. Cells were resuspended in 43.74 mL 1× PBS. The following volumes were added to the reaction in order with vortexing between each addition:

i. 500 μL 100 mM cargo-alkyne or cargo-azide (final concentration of 1 mM)
ii. 760 μL 1:2 1:2 volume mixture of 20 nM CuSO_4_: 50 mM THPTA
iii. 2.5 mL 100 mM aminoguanidine
iv. 2.5 mL 100 mM sodium ascorbate

The reaction was left running for 4 h before termination. Termination of reaction was achieved by washing cells three times with PBSA. Following termination 2×10^6^ cells were washed three with PBSA. Cells were resuspended in 220 μL 1× PBS. Biotin click reaction was conducted by adding the following volumes to the cells for the reaction in order with vortexing between each addition:

i. 1.25 μL 20 mM biotin-azide/alkyne (final concentration of 100 µM)
ii. 3.8 μL 1:2 1:2 volume mixture of 20 nM CuSO_4_: 50 mM THPTA
iii. 12.5 μL 100 mM aminoguanidine
iv. 12.5 μL 100 mM sodium ascorbate

The reaction was allowed to proceed for 15 min. Termination of reaction was achieved by washing cells three times with PBSA. Cells were then labelled for flow cytometry detection of biotin and full-length display (SI Table 4).

### PTP1B preparation

The vector to express the catalytic domain of PTP1B (pET-28 His-TEV-PTP1B-cat)^67^ was a generous gift from the Shah Lab at Columbia University. The PTP1B expression plasmid was transformed into chemically competent BL21(DE3) cells (generously gifted by the Nair Lab at Tufts University) and streaked on LB plates containing 50 µg/mL kanamycin. Plate(s) were incubated at 37 °C for 16 hours until colonies formed. Individual colonies were inoculated in 5 mL LB containing 50 µg/mL kanamycin and incubated at 37 °C for 16 hours shaking at 300 RPM for 16 hours until saturated. Following saturation, 5 mL cell suspension was added to 1 L TB supplemented with 50 µg/mL kanamycin in a plastic baffled flask. TB culture(s) were grown at 37 °C for 2-4 hours until reaching an OD_600_ of 0.5. Cells were placed on ice until cooled, and 500 µL 1 M isopropyl *β*-D-1-thiogalactopyranoside (IPTG; sterile filtered before use) was added to induce PTP1B expression. Induced cells were placed at 16 °C shaking at 300 RPM for 16-24 hours. Following induction, cells were pelleted at 3200 rcf for 30 minutes at 4 °C. Supernatant was decanted and the cells from a 1 L induced culture were resuspended in 20-40 mL of ice-cold lysis buffer (50 mM tris, pH 8.0, 500 mM NaCl, 20 mM imidazole, 10% glycerol (2 mM 2-mercaptoethanol (BME) and protease inhibitor tablet (ThermoFisher Scientific) added immediately before use). Cells suspended in lysis buffer were flash frozen in liquid nitrogen and stored at –80 °C for a minimum of 24 hours. Frozen cell stocks were thawed in a water bath. Thawed cells were subjected to 13×10 second ultrasonic pulses in a water bath at room temperature with 20 second pauses in between. Cells were then centrifuged for 1 hour at 12,000 rcf at 4 °C, and supernatant was sterilized using a syringe filter while cell pellets were discarded. Supernatant was purified using Ni-NTA resin (GenScript) as described previously^30^. The eluant was then buffer exchanged into 1× PBS pH 7.4 using Amicon Ultra-15 centrifugal filter units (10 kDa MW cut-off; Millipore Sigma). Protein concentrations were measured via absorbance at 280 nm on a NanoDrop^TM^ Microvolume UV-Vis Spectrophotometer (ThermoFisher Scientific). SDS-PAGE was run to confirm the molecular weight of the purified proteins using 4-12% Bis-Tris mini gels and SimplyBlue Safestain. Within 3 days of purification, purified protein was stored by diluting to 1 mg/mL, mixing 1:1 by volume with glycerol, aliquoting 200 µL amounts to microcentrifuge tubes, flash freezing in liquid nitrogen, and storing at –80 °C. PTP1B was recovered from glycerol stocks by buffer exchanging into 1× PBS pH 7.4 using Amicon Ultra-0.5 centrifugal filter units (10 kDa MW cut-off; Millipore Sigma). PTP1B stocks were stored exclusively at 4 °C. Frozen glycerol stocks were buffer exchanged for sorting less than 24 hours before the start of enrichment and used for binding analysis within 3 days of buffer exchange to ensure enzyme integrity.

### Antigen biotinylation

Donkey IgG (Jackson Immuno-Research) and PTP1B (performed within 2 days of purification as described above; timing ensured frozen stocks could be prepared by day 3 post-purification) were diluted from stock concentrations to 1-2 mg/mL in ice cold 1× PBS. 100 mM EZ-Link NHS-LC biotin solution was prepared in dimethylformamide and mixed extensively to dissolve. Biotinylation reactions were carried out by incubating 100 µL, 200 µL, or 1 mL aliquots of protein stocks with 2-6 fold molar excess of 100 mM EZ-Link NHS-LC biotin solution at room temperature for 30 min. Reactions were quenched with 0.5 M tris, 0.02% NaN_3_, pH 7.4 using a volume equal to 10% of the reaction volume. Upon completion of the reaction, proteins were desalted twice using Zeba Spin Desalting Columns (Thermo Fisher; 7 kDa MW cutoff) according to the manufacturer’s specifications. The concentration of the resulting protein was measured via the absorbance of 280 nm light on a Nano Drop One instrument (ThermoFisher Scientific). The extent of biotinylation was estimated using the HABA-Avidin Premix according to the manufacturer’s protocol (ThermoFisher Scientific, catalog number 28005). Only protein samples that achieved a biotinylation ratio of 1-2 biotin per protein were used in enrichments to reduce the chances of enriching the library for biotinylated antigen. For biotinylated PTP1B, within 1 day of biotinylation, purified protein was stored by diluting to 1 mg/mL, mixing 1:1 by volume with glycerol, aliquoting 200 µL amounts to microcentrifuge tubes, flash freezing in liquid nitrogen, and storing at –80 °C. Biotinylated PTP1B was recovered from glycerol stocks by buffer exchanging into 1× PBS pH 7.4 using Amicon Ultra-0.5 centrifugal filter units (10 kDa MW cut-off; Millipore Sigma). Biotinylated PTP1B stocks were stored exclusively at 4 °C. Frozen glycerol stocks were buffer exchanged for sorting less than 24 hours before the start of enrichment and used for binding analysis within 3 days of buffer exchange to ensure enzyme integrity. Enzyme activity was validated before every sort (see below). Biotinylated donkey IgG stocks were stored exclusively at 4 °C and were prepared from unmodified donkey IgG stocks less than 24 hours before the start of enrichment and used for binding analysis within 3 days of preparation.

### Bead-based enrichment and FACS

#### Donkey IgG

Two cryo vials of the depleted library were thawed at room temperature and inoculated into 1 L SD-SCAA –Trp –Leu –Ura + pen-strep. The culture was grown to saturation at 30 °C with shaking overnight. The library was spun down and resuspended at an OD_600_ of 1 and grown until mid-log phase (OD_600_ between 2-5; duration 4-8 hours). The remainder of the culture was placed at 4 °C as a short-term stock. Cells were pelleted (5 min at 2,400 rcf and 4 °C) and resuspended at an OD_600_ of 1 in 1 L induction media (SG-SCAA –Trp –Leu –Ura + pen-strep at a 1:100 dilution) containing 1 mM ncAA and grown for 16 hours at 20 °C with shaking.

100 μL Biotin Binder Dynabeads were prepared by washing three times on the magnet in 1 mL ice-cold PBSA. Washes were done by incubation on a DynaMag-2 (ThermoFisher Scientific) for 2 min followed by removal of supernatant. After washing, the beads were resuspended in 100 μL ice-cold PBSA. Beads were aliquoted in 50 μL volumes and incubated with biotinylated donkey IgG (33 pmol antigen/10 μL beads) on a rotary wheel at 4 °C for 2-16 hours. Each 50 μL aliquot was used in a different sorting track described below.

For the first round of sorting, 2×10^10^ induced cells per sorting track were pelleted in at 2,400 rcf for 5 min, washed and resuspended in 15 mL ice-cold PBSA. For each sort, prepared beads were washed and 50 μL were added to the cells, then placed on a rotary wheel (i) at 4 °C for 2 hours, and (ii) at 4 °C for 24 hours. After this incubation, the cells were placed on a DynaMag-15 for 2 min and the supernatants were transferred to a new 15 mL conical tube. The remaining cells from both tracks were resuspended in 0.5 M tris-HCl pH 4.1 for 1 h at 4 °C on a rotary wheel. After the incubation with the denaturant, the cells were placed on the DynaMag-15 for 2 min and the supernatants were transferred to a new 15 mL conical tube. The remaining cells from both tracks were then washed twice more on the magnet by removing the cells after incubation on the magnet and resuspending the beads in PBSA. Cells that were attached to the beads were rescued in 100 mL SD-SCAA –Trp –Leu –Ura + pen-strep for overnight growth until saturation. After the initial sort (Round 1), subsequent rounds of sorting (Rounds 2 and 3) were conducted using 1 billion cells and 10 μL of antigen-coated magnetic beads, while keeping the conditions consistent for each track.

Following four rounds of bead-based enrichments, each of the populations were sorted from the third and fourth rounds using fluorescence-activated cell sorting (FACS) on a BioRad S3e cell sorter. To prepare samples for sorting, 2×10^7^ induced cells from each population were washed three times in 1 mL PBSA. Cells were incubated with 100 μL of 200 nM biotinylated donkey IgG on a rotary wheel (i) at 4 °C for 2 hours, and (ii) at 4 °C for 24 hours. After the incubation, cells were washed three times with 1 mL ice-cold PBSA. The remaining cells were resuspended in 0.5 M tris-HCl pH 4.1 for 1 h at 4 °C on a rotary wheel. After the incubation with the denaturant, the cells were washed three times with 1 mL ice-cold PBSA and prepared for labelling. For primary labeling, cells were resuspended in 100 μL chicken anti-cMyc at a dilution of 1:500 in PBSA and were incubated for 30 min on a rotary wheel at 300 RPM at RT. Following primary labelling cells were washed twice with 1 mL ice-cold PBSA. During secondary labeling, cells were incubated in the dark on ice with 100 μL mixture of goat anti-chicken Alexa Fluor 647 and streptavidin Alexa Fluor 488 at a dilution of 1:500 in PBSA. For each sort, between 50,000 and 10 million events were sorted by FACS and approximately 300– 10,000 c-Myc-positive, antigen binding-positive events were collected. Sorted cells were rescued in a final 5 mL SD-SCAA –Trp –Leu –Ura + pen-strep at 30 °C with shaking at 300 RPM; populations were grown to saturation prior to subsequent evaluations.

Analytical flow cytometry was used to evaluate the binding properties of enriched populations. Cells from each population were induced and prepared for flow cytometry as described above. Briefly, for each round of sorting after the first round of positive enrichment, induced cells were evaluated for binding to donkey IgG by flow cytometry. Cells were subjected to the same wash conditions as described. Samples were labeled with 200 nM biotinylated antigen and chicken anti-cMyc at a 1:500 dilution in PBSA. For secondary labeling, cells were labeled with goat anti-chicken Alexa Fluor 647 and streptavidin Alexa Fluor 488 a 1:500 dilution.

#### PTP1B

Two cryo vials of the depleted library were thawed at room temperature and inoculated into 1 L SD-SCAA –Trp –Leu –Ura + pen-strep. The culture was grown to saturation at 30 °C with shaking overnight. The library was spun down and resuspended at an OD_600_ of 1 and grown until mid-log phase (OD_600_ between 2-5; duration 4-8 hours).

The remainder of the culture was placed at 4 °C as a short-term stock. Cells were pelleted (5 min at 2,400 rcf and 4 °C) and resuspended at an OD_600_ of 1 in 1 L induction media (SG-SCAA –Trp –Leu –Ura + pen-strep at a 1:100 dilution) containing 1 mM OBeY and grown for 16 hours at 20 °C with shaking.

60 μL Biotin Binder Dynabeads were prepared by washing three times on the magnet in 1 mL ice-cold PBSA. Washes were performed by incubation on a DynaMag-2 (Thermo Fisher Scientific) for 2 min followed by removal of supernatant. After washing, the beads were resuspended in 600 μL ice-cold PBSA and incubated with biotinylated PTP1B (33 pmol antigen/10 μL beads) on a rotary wheel at 4 °C for 2-16 hours. PTP1B used for sorting was always recovered from glycerol stocks on the day before the enrichment to ensure freshness of stock.

For the first round of sorting, 1×10^10^ induced cells were pelleted and washed three times with 50 mL PBSA. Beads incubated with biotinylated PTP1B were washed three times with ice-cold PBSA as described above and resuspended in 60 µL cold PBSA. 10 µL beads were used in a plate reader assay to ensure PTP1B was still active on the beads before enrichment. For activity assay, beads were aliquoted to select wells in a black-walled 96-well plate and diluted to 20 µL with 10 µL cold PBSA. 180 µL 5 mM 6,8-difluoro-4-methylumbelliferyl phosphate (DiFMUP) was added to each well, and fluorescence (ex. 360 nm, em. 460 nm) was measured over 10 minutes on a SpectraMax i3X plate reader (Molecular Devices). 20 µL of 200 nM PTP1B, 10 µL washed, bare beads (diluted to 20 µL), and 5 mM DiFMUP alone were used as controls. Once PTP1B was determined to be active on the beads, the induced, washed cells were resuspended in 10 mL 1× PBS and incubated with the remaining 50 µL PTP1B-coated beads for 2 hours at 4 °C on a rotary wheel. Following incubation, the cell + bead mixture was incubated on the DynaMag-15 for 5 minutes. Cells were carefully removed without disturbing beads and discarded. Beads were washed once with 15 mL ice-cold PBSA, and then rescued in 15 mL SD-SCAA –Trp –Leu –Ura + pen-strep for overnight growth at 30 °C with shaking until saturation. Once cells were saturated, they were pelleted at 2400 rcf for 5 minutes and resuspended in 1 mL SD-SCAA –Trp –Leu – Ura + pen-strep in a 1.7 mL tube. Cell suspension was placed on the DynaMag-2 for 2 minutes, and cells were recovered in 100 mL SD-SCAA –Trp –Leu –Ura + pen-strep, without the beads. Cells were grown again overnight to saturation at 30 °C with shaking, and then either used immediately or stored at 4 °C for future rounds of enrichment or analysis.

The second round of enrichment started by growing a 100 mL overnight culture of the Round 1 cells to saturation. Cells were diluted and induced as described above, and a 250 mL induction volume was used to ensure there would be 1 billion induced cells for each new enrichment track. Biotin binder beads were prepared again as described above, using 50 µL beads instead of 60 µL, and PTP1B activity was again confirmed on the beads before use. Induced cells were washed three times with 50 mL PBSA, and then split in to five aliquots of 1 billion cells. For <u>Track A</u>, 1 billion cells were resuspended in 1 mL 1× PBS and incubated with 10 µL PTP1B-coated beads for 2 hours on the rotary wheel at 4 °C, and binding cells were recovered as in round 1. For <u>Track B</u>, 1 billion cells were resuspended in 1 mL 1× PBS and incubated with 10 µL PTP1B-coated beads for 24 hours on the rotary wheel at 4 °C. After 24 hours incubation, cells + beads mixture was placed on the DynaMag-2 for 2 minutes, and non-bound cell suspension was removed. Beads were washed once with ice-cold PBSA without disturbing the magnetic beads. Beads were then resuspended in 1 mL 0.5 M tris-HCl, pH 4.1, and placed back on the rotary wheel at 4 °C for 24 hours. Following incubation, cells that remained bound to the beads were recovered as in round 1. For <u>Track C</u>, the conditions were the same as in Track B, except incubation with PTP1B-coated beads occurred at room temperature instead of 4 °C. For <u>Track D</u>, all the conditions were the same as Track C, except the denaturing step used 0.1 M urea for 30 minutes at room temperature instead of 0.5 M tris-HCl for 24 hours. For <u>Track E</u>, 1 billion induced and washed cells were resuspended in 1 mL 50 nM PTP1B and incubated at room temperature for 24 hours on a rotary wheel. Following overnight incubation, cells were pelleted and washed twice with 1 mL PBSA. Cells were resuspended in 1 mL PBSA and incubated on the rotary wheel for 10 minutes at room temperature. Pelleting and incubation steps were repeated twice more for a total of three 10-minute incubations with PBSA as a “gentle” denaturing step. Following washes, cells were incubated with 10 µL washed Biotin Binder beads for 2 hours at 4 °C on the rotary wheel to capture cells bound to biotinylated PTP1B. Following incubation, cells were recovered as in round 1. Round 3 of bead enrichment proceeded the same way as round 2.

Following the 3^rd^ round of enrichment, cells were depleted of the enriched wildtype populations. Each track of cells was passaged, diluted, and induced as described before, however induction was in the absence of any ncAA. 10 million induced cells from each track were washed and labelled for HA and c-Myc detection as described above and sorted on an S3e cell sorter (BioRad) to isolate cells that show HA detection, but not c-Myc detection, representing the desired population that contains a stop codon. 1.5 million cells were recovered from each track in 5 mL SD-SCAA –Trp – Leu –Ura + pen-strep.

Following depletion (populations termed “WTd”), cells were subjected to positive enrichments using FACS. To prepare samples for sorting, 2×10^7^ induced cells from each track were washed three times in 1 mL PBSA. Cells were incubated with 200 μL of 250 nM biotinylated PTP1B on a rotary wheel at 4 °C for 24 hours. After the incubation, cells were washed two times with 1 mL ice-cold PBSA. The remaining cells were resuspended in 0.5 M tris-HCl pH 4.1 for 24 h at 4 °C on a rotary wheel. After the incubation with the denaturant, the cells were washed three times with 1 mL ice-cold PBSA and prepared for labelling. For primary labeling, cells were resuspended in 200 μL chicken anti-cMyc at a dilution of 1:500 in PBSA and were incubated for 30 min on a rotary wheel at 300 RPM at RT. Following primary labelling cells were washed twice with 1 mL ice-cold PBSA. During secondary labeling, cells were incubated in the dark on ice with 200 μL mixture of goat anti-chicken Alexa Fluor 647 and anti-biotin PE at a dilution of 1:500 in PBSA. For each sort, approximately 50,000 to 10 million events were sorted by FACS and approximately 300–10,000 c-Myc-positive, antigen binding-positive events were collected. Sorted cells were rescued in a final 5 mL SD-SCAA –Trp –Leu – Ura + pen-strep at 30 °C with shaking at 300 RPM; populations were grown to saturation prior to subsequent evaluations.

Analytical flow cytometry was used to evaluate the binding properties of enriched populations (including pre-enrichment). Cells from each population were induced and prepared for flow cytometry as described above. Samples were incubated with 250 nM biotinylated PTP1B, followed by primary labelling with chicken anti-cMyc at a 1:500 dilution in PBSA. For secondary labeling, cells were labeled with goat anti-chicken Alexa Fluor 647 and streptavidin Alexa Fluor 488 or anti-biotin PE a 1:500 dilution.

### Binder characterization

Plasmid DNA was isolated from desired populations sorted against donkey IgG and PTP1B and transformed into *E. coli* to prepare single clones for sequence analysis. 13 clones were sequenced from the sorted population against donkey IgG (OBeY-24 B4F2) as well as 38 total clones from populations of PTP1B binders (Tracks A and C). Unique clones were transformed into RJY100, plated on solid SD-SCAA –Trp –Leu – Ura media, and grown at 30 °C until colonies appeared (2–3 days). Colonies were inoculated in 5 mL of SD-SCAA –Trp –Leu –Ura + pen-strep and allowed to grow to saturation at 30 °C with shaking (300 RPM) for 2-3 days. Following saturation, the cultures were diluted to an OD_600_ of 1 in fresh SD-SCAA –Trp –Leu –Ura + pen-strep and grown at 30 °C until mid-log phase (OD_600_ 2-5; duration 4-8 hours). Cells were pelleted (5 min at 2,000 rcf and 4 °C) and resuspended to an OD_600_ of 1 in SG-SCAA – Trp –Leu –Ura + pen-strep with 1 mM desired ncAA. Cells were prepared for flow cytometry analysis as described above – see individual experiments for detailed parameters.

For binding only conditions (cells bound with antigen followed by flow cytometry labelling; no attempt to disrupt noncovalent interactions with denaturing conditions), following induction cells were washed three times in PBSA and incubated with either with 200 nM unconjugated antigen and chicken anti-cMyc at a 1:500 dilution in PBSA, or 50-250 nM biotinylated antigen. Labeling and subsequent data collection was performed as outlined above.

For binding and denaturing conditions (cells bound with antigen followed by incubation under denaturing conditions followed by flow cytometry labelling), following induction cells were washed three times in PBS and incubated with biotinylated antigen. Cells were incubated with antigen for various times at either 4 °C, room temperature, or 37 °C (see individual experimental details in main text or figure captions). Following incubation, cells were washed three times in PBSA and the pellets were incubated with 8 M urea (200 mM EDTA, 25 mM tris, pH 8.4) at room temperature for specified time. Subsequently, cells were washed three times in PBSA labelled for flow cytometry as described above.

### Binding affinity (K_D_) data collection and analysis

1.5 million freshly induced RJY100 cells displaying OPG-substituted scFvs (Donkey 1.1, DXB4.2.4, 4.2.19 and 4.2.24) were aliquoted in 1.7 mL microcentrifuge tubes, pelleted, washed three times with 1× PBSA, and then resuspended in 1 mL PBSA. 300 μL washed cells were then combined with 0.3 μL of chicken anti-cMyc, vortexing gently to mix. To prepare for binding titration, donkey IgG was added to 96-well V-bottom plates starting at 1 μM final concentration with seven subsequent 4-fold dilutions and a final PBSA blank (no IgG). To each of these 8 IgG concentrations and one PBSA blank, 10 μL cells incubated with chicken anti-cMyc were added, resulting in approximately 15,000 cells per sample, and incubated on the orbital shaker at 150 RPM at room temperature overnight (final chicken anti-cMyc dilution of 1:1000; wild-type Donkey1.1 and DXB4.2.4, 4.2.19, 4.2.24 were added to wells containing the 8 different concentrations of biotinylated donkey IgG and to wells containing the PBSA blank). After the overnight incubation, samples were diluted and washed 3-5 times with ice-cold PBSA. Secondary labeling and flow cytometry data collection was performed as described for scFvs above, and 3,000 events were collected. Reagents used for secondary labeling are listed in SI Table 4. All titrations were performed in technical triplicates. Data analysis to obtain binding affinities (K_D_) were done using steps similar to those outlined in the “Flow cytometry data collection and analysis” section above, with a few modifications described in our previous work^30^. Data was exported and analyzed on GraphPad Prism. The background corrected data for biotinylated antigen detection were normalized between 0 and 1, then using the “Receptor binding – Saturation binding” model and “One site --Specific binding” equation, the data was fitted to a curve. K_D_ was estimated including 95% confidence interval.

### Production and purification of soluble scFvs

#### Donkey IgG Binders

DNA encoding clones DXB4.2.4 and DXB4.2.19 were amplified via PCR, with the pRS-scFv-Fwd and pRS-scFv-Rev primers, (SI Table 5) and then cloned into pRS314-GAL-4420-his6 (Trp marker)^68,69^ vector digested at the NheI and BamHI restriction sites using Gibson Assembly. The assemblies were transformed into chemically competent *E. coli* DH5*α*Z1, plated on selective media (LB + 50 µg/mL ampicillin) and grown at 37°C overnight. Resulting colonies were inoculated and grown to saturation in 5 mL selective liquid media (LB + 50 µg/mL ampicillin). Plasmid DNA was then isolated from these cultures using *E. coli* plasmid DNA mini-prep kits. The cloned plasmids were digested at the NheI and BamHI sites to confirm the presence of correctly sized inserts by performing gel electrophoresis. Finally, the plasmids were sequence verified using Sanger sequencing (Quintara Biosciences).

Plasmid constructs were cotransformed into Zymo-competent RJY100 cells with the OTS construct pRS315-KanRmod-AcFRS, plated on selective media (–Trp –Leu – Ura) and grown at 30 °C until colonies appeared (∼3 days). Colonies were inoculated from each plate into 5 mL liquid cultures, which were grown to saturation. The saturated cultures were passaged once, diluted into 50 mL minimal media, and grown to saturation. Saturated cultures were pelleted and resuspended in 100 mL YPG + pen-strep with 1 mM ncAA (OBeY or OPG) and incubated for 4 days at 20 °C with shaking at 300 RPM. At the end of induction, the cultures were pelleted for 35 min at 3,214 rcf and filtered using a 0.2 μm filter. Filtrates for scFv secretions were purified using Ni-NTA resin (GenScript) as described previously^30^. The eluant was then buffer exchanged into 1× PBS pH 7.4 using Amicon Ultra-15 centrifugal filter units (10 kDa MW cut-off; Millipore Sigma). Protein concentrations were measured via absorbance at 280 nm on a NanoDrop^TM^ Microvolume UV-Vis Spectrophotometer (ThermoFisher Scientific). SDS-PAGE was run to confirm the molecular weight of the purified proteins 4-12% Bis-Tris mini gels and SimplyBlue Safestain. All soluble proteins were mixed in a 1:1 volumetric ratio with glycerol, aliquoted in 200 μL volumes, flash frozen with liquid nitrogen, and stored at −80 °C for long-term storage.

#### PTP1B Binders

DNA encoding clones PXA.1 and PXC.4 were amplified *via* PCR, with the pCHA-scFv-NheI-Fwd and pCHA-scFv-XmaI-Rev primers, (SI Table 5) and then cloned into the pCHA-FcSup-TAA^30^ (Trp marker) vector digested at the NheI and XmaI restriction sites using Gibson Assembly. The assemblies were transformed into chemically competent *E. coli* DH5*α*Z1, plated on selective media (LB + 50 µg/mL ampicillin) and grown at 37 °C overnight. Resulting colonies were inoculated and grown to saturation in 5 mL selective liquid media (LB + 50 µg/mL ampicillin). Plasmid DNA was then isolated from these cultures using *E. coli* plasmid DNA mini-prep kits. The cloned plasmids were digested at the NheI and XmaI sites to confirm the presence of correctly sized inserts by performing gel electrophoresis. Finally, the plasmids were sequence verified using Sanger sequencing (Quintara Biosciences).

Plasmid DNA were cotransformed into Zymo-competent RJY100 cells with the OTS construct pRS315-KanRmod-AcFRS, plated on selective media (–Trp –Leu –Ura) and grown at 30 °C until colonies appeared (∼3 days). Colonies were inoculated from each plate into 5 mL liquid cultures, which were grown to saturation. The saturated cultures were passaged once, diluted into 50 mL minimal media for overnight saturation, and diluted again in 500 mL minimal media for overnight saturation. 500 mL saturated cultures pelleted and induced in 1 L YPG + pen-strep with 1 mM ncAA (OBeY or OmeY) for 4 days at 20 °C with shaking at 300 RPM. At the end of induction, the cultures were pelleted for 35 min at 3,214 rcf and filtered using a 0.2 μm filter. Filtrates for scFv-Fc secretions were purified using protein A resin (GenScript) as described previously^30,64^. The eluant was then buffer exchanged into 1× PBS pH 7.4 using Amicon Ultra-15 centrifugal filter units (10 kDa MW cut-off; Millipore Sigma). Protein concentrations were measured via absorbance at 280 nm on a NanoDrop^TM^ Microvolume UV-Vis Spectrophotometer (ThermoFisher Scientific). SDS-PAGE was run to confirm the molecular weight of the purified proteins using 4-12% Bis-Tris mini gels and SimplyBlue Safestain.

### In-Solution Bead-Based Pulldown

100 μL magnetic beads were washed on a magnet and coated with biotinylated donkey IgG as described above (see “Bead-based enrichment and FACS”). Following washes, beads were aliquoted into 5 separate tubes (20 μL original bead volume/tube). OBeY- and OPG-substituted purified scFvs were added to the beads at 10× molar excess. The solution was then incubated for 6 h at 37 °C on a rotary wheel (300 RPM). Following incubation, the beads were placed on the magnets and subjected to washing, then each sample was split in two, containing 10 μL each. Half of the samples were stored at 4 °C as pellets until ready to be used, and the other half were incubated with 1 mL urea buffer (8 M urea, 200 mM EDTA, 25 mM tris, pH 8.4) for 24 h at room temperature on a rotary wheel (300 RPM). Following denaturation with urea, beads were washed on the magnet with PBSA. All samples were then labeled for full length scFv with the primary antibody chicken anti-cMyc using previously described protocols. Secondary labeling was performed with Alexa Fluor goat anti-chicken 647 and Streptavidin 488 (see SI Table 4 for labelling conditions). Samples were run on the flow cytometer and analyzed as previously described.

### In-solution PTP1B Binding

60 µL Biotin Binder beads were washed with the DynaMag-2 as described above. Beads were incubated with PTP1B overnight (33 pmol/10 µL beads), and then washed again with ice-cold PBSA. PTP1B-coated beads were separated to 6 different containers, representing 10 uL original bead volume per tube. Beads were resuspended in 5-fold molar excess of either PXA.1-OBeY, PXA.1-OmeY, PXC.4-OBeY, PXC.4- OmeY, or FAP-Fc^59^ (non-binding control) and left to incubate on a rotary wheel at 4 °C for 3 hours. Following incubation, beads were washed on the magnet with PBSA, including a 10 uL bare bead control for flow cytometry. All samples were then labeled for full length scFv-Fc with the primary antibody mouse anti-HA using previously described protocols. Secondary labeling was done with Alexa Fluor goat anti-mouse 647 and Streptavidin 488. Samples were run on the flow cytometer and analyzed as previously described; see SI Table 4 for labelling conditions.

### In-solution PTP1B Activity Assay

100 µL solutions were prepared containing 50 nM PTP1B and 1 µM binding scFv-Fcs (20-fold molar excess), with an additional PTP1B-only control (no scFv-Fc added). Solutions were incubated for 30 minutes at room temperature on the rotary wheel. Following binding, 20 µL from each sample was aliquoted in triplicate to a black 96-well plate. 180 µL 5 mM 6,8-difluoro-4-methylumbelliferyl phosphate (DiFMUP) was added to each well, and fluorescence (ex. 360 nm, em. 460 nm) was measured over 10 minutes on a SpectraMax i3X plate reader (Molecular Devices). 5 mM DiFMUP alone was used as a control. The enzymatic fractional activity was determined by comparing the slope of the fluorescence over time plots in each sample to the PTP1B only control, as described previously^47^.

## Acknowledgments

The construction, validation, and screening of the chemically expanded antibody library against Donkey IgG and all subsequent clone characterization was supported by a grant from the National Cancer Institute of the National Institutes of Health (R21CA214239) as part of the Innovative Molecular and Cellular Analysis Technologies program. Screening of the chemically expanded antibody library against PTP1B and subsequent clone characterization was supported by a grant from the National Institute of General Medical Sciences of the National Institutes of Health (R35GM133471). The content is solely the responsibility of the authors and does not necessarily represent the official views of the National Institutes of Health. We thank the laboratory of Matthew Bogyo for providing N_3_-PEG_4_-vinylsulfone and N_3_-bAGbA-vinylsulfone. We thank the laboratory of Neel Shah for providing the plasmid pET-28 His-TEV-PTP1B-cat. We thank the laboratory of Nikhil Nair for providing BL21(DE3) cells.

## Notes

### Competing Interest Statement

The authors have declared no competing interest.

