## Supporting Information for "Design, Construction, and Validation of a Yeast-Displayed Chemically Expanded Antibody Library"

**Authors/Affiliations**

Arlinda Rezhdo[†], Rebecca L. Hershman[†], and James A. Van Deventer[*, †, ‡]

[†]Chemical and Biological Engineering Department, Tufts University, Medford, Massachusetts 02155, USA

[‡]Biomedical Engineering Department, Tufts University, Medford, Massachusetts 02155, USA

*Corresponding author

#### Supporting Information

**Acceptable Quality of Sublibraries for Incorporation into Pooled Library**

*Characterization of full-length display in the absence and presence of ncAA*

The first criterion for acceptable sublibrary quality was the evaluation of full-length display in the absence of a ncAA and in the presence of AzF and OPG. To quantify the presence of the encoded stop codon in the scFv sequence, full-length display in the absence of a ncAA was evaluated. A population that displays full-length clones under such conditions indicates absence of a stop codon. A second attempt at making a sublibrary was undertaken for each case that showed full-length clones ≥1% of the displayed clones. All sublibraries that had <1% full-length clones were moved forward for further evaluation. The pooled library yielded 0.2% of displayed clones to be full length in the absence of a ncAA. This value is well below the 1% threshold set forth in the quality check criteria, supporting the assumption that a stop codon is present in >99% of the library and can be used for the incorporation of a ncAA. To evaluate the display in the presence of AzF and OPG, flow cytometry was used to validate the ability of the sublibraries to incorporate the desired ncAA when they were induced in the presence of AzF or OPG.

*Truncation analysis*

The second criterion for acceptable sublibrary quality was evaluation of instances of truncated clones. To that end, we calculated the percentage of yeast populations displaying truncated scFvs in the presence of a ncAA. We defined truncation as an scFv that is not displaying full length due to termination from a stop codon in the encoding sequence. The XYZ diversification scheme allows TAG stop codons in the CDR-H3 sequence, and in the presence of a ncAA and suppression machinery, the incorporation of the ncAA will depend on the efficiency of the orthogonal translation system (OTS) and the surrounding amino acid sequence. Therefore, truncation levels will in part depend on the probability of a TAG codon being encoded in the CDR-H3 sequence, and efficiency of the OTS to incorporate a ncAA in the sequence. Experimental values, normalized to the OTS efficiency (see *Material and Methods*), were compared to theoretical truncation based only on the CDR-H3 diversification scheme. Truncation levels obtained experimentally increased in longer loop lengths, consistent with the increase in probability of TAG presence in the loop and in agreement with the calculated theoretical truncation.

*Library validation via DNA sequencing*

The third and final criterion for acceptable sublibrary quality was validation of amino acid sequence diversity. We validated the design sequence of each sublibrary by sequencing a sample of approximately six clones, totaling to 317 clones for the whole library. Three characteristics were deemed necessary for a sequence to pass quality check: (1) sequence of the CDR-H3 region is unique and matches the designed “XYZ” diversification scheme, (2) TAG codon is present and in the intended location, and (3) there are no frameshift mutations in the rest of the sequence. A sublibrary was considered “verified” if at least 80% of the sequenced clones had all three characteristics. Deletions or insertions in the CDR-H3 or frameshift mutations were also evaluated, and sequences that had such errors were cause for sublibraries failing quality check. Out of 54 sub-libraries, 13 failed initial sequencing quality check and were remade. The new sublibraries were re-characterized as previously, and were sequence verified. Five of the new sublibraries did not pass quality check a second time, and they were excluded from further experiments. The sequences obtained from all the individual sublibraries were analyzed and compared to theoretical sequence expectations calculated from the diversification scheme. Obtained sequences matched the overall diversification scheme of the CDR-H3 region both at the nucleotide and the amino acid level. Furthermore, the observed amino acid frequencies matched the design frequencies for individual loop lengths. Each sequence was checked for the presence of the TAG codon, and only 2 sequences out of 317 did not contain an encoded stop codon. Finally, one sublibrary (L50 loop length 9) did not regrow from glycerol stocks and was omitted from the library despite passing initial quality check.

**CDR-H3 Loop Length**

| TAG Position | 9 | 10 | 11 | 12 | 13 | 14 | 15 | 16 | 17 |
|---|---|---|---|---|---|---|---|---|---|
| L1 | | 1.00 | | 0.85 | 1.25 | 1.20 | 2.05 | 1.15 | 3.05 |
| L28 | 1.00 | 1.30 | 2.20 | 2.90 | 1.00 | 6.05 | 4.45 | 0.85 | 2.25 |
| L50 | | 2.65 | 2.65 | 1.15 | 3.00 | 1.55 | 1.70 | 0.60 | 0.25 |
| L93 | 0.80 | 1.65 | | 2.65 | 0.95 | 3.60 | 3.75 | 3.70 | 1.17 |
| H31 | | 1.15 | 2.30 | 2.00 | 3.90 | 3.35 | 6.10 | 1.50 | 1.45 |
| H54 | | 2.60 | 4.00 | 1.95 | 1.75 | 5.15 | 2.40 | 0.95 | 1.50 |

6.10
5.00
4.00
2.00
0.60

Number of transformants ($\times 10^7$)

**SI Figure 1.** Number of transformants for each separately constructed sublibrary. Total number of transformants: $1.1\times10^9$.

### Calculations of full-length clones in absence of a ncAA

##### Absence of ncAA

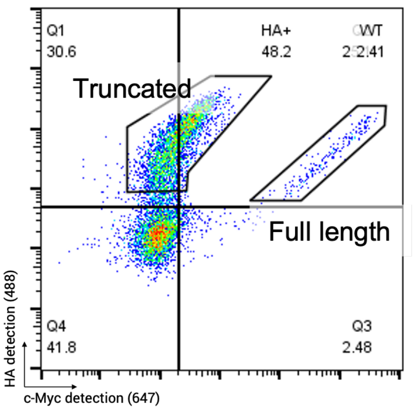


##### Full-length analysis and comparison

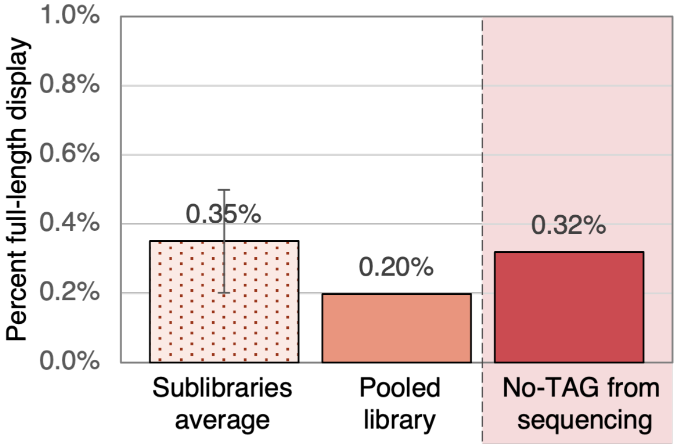


$$\% \text{ Full length} = \frac{\text{Full length}}{\text{Truncated} + \text{Full length}}$$

**CDR-H3 Loop length**

| TAG Positions | 9 | 10 | 11 | 12 | 13 | 14 | 15 | 16 | 17 |
|---|---|---|---|---|---|---|---|---|---|
| L1 | | 0.49 | | 0.94 | 0.99 | 0.76 | 0.98 | 0.94 | 0.61 |
| L28 | 0.04 | 0.15 | 0.16 | 0.11 | 0.10 | 0.07 | 0.10 | 0.05 | 0.26 |
| L50 | | 0.06 | 0.05 | 0.08 | 0.27 | 0.19 | 0.11 | 0.02 | 0.26 |
| L93 | 0.82 | 0.39 | | 0.66 | 0.16 | 0.15 | 0.14 | 0.20 | 0.27 |
| H31 | | 0.38 | 0.20 | 0.23 | 0.13 | 0.59 | 0.33 | 0.80 | 0.15 |
| H54 | | 0.35 | 0.31 | 0.47 | 0.71 | 0.27 | 0.17 | 0.95 | 0.29 |

*Values are %

**SI Figure 2**. Calculations of full-length clones in the absence of a ncAA; flow cytometry evaluation of “truncated” and “full length” clones in the absence of a ncAA. Percent full length clones were calculated using the formula shown. The analysis was performed for all sublibraries and full-length analysis comparison was done as an average of all sublibraries, the pooled library, and from sequencing. This shows that clones without TAG codons are present in the library at approximately 0.35%.

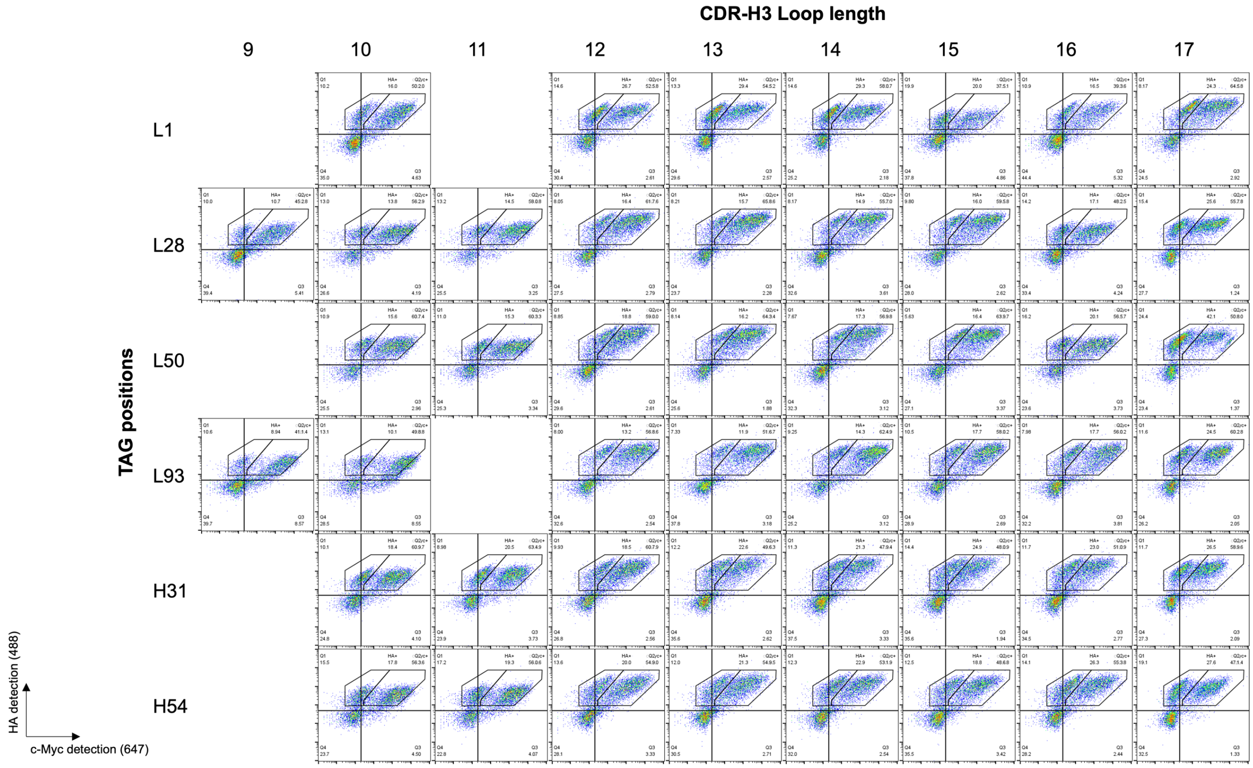


**SI Figure 3.** Evaluation of AzF incorporation in each sublibrary using flow cytometry shows consistent incorporation and display levels for all sublibraries.

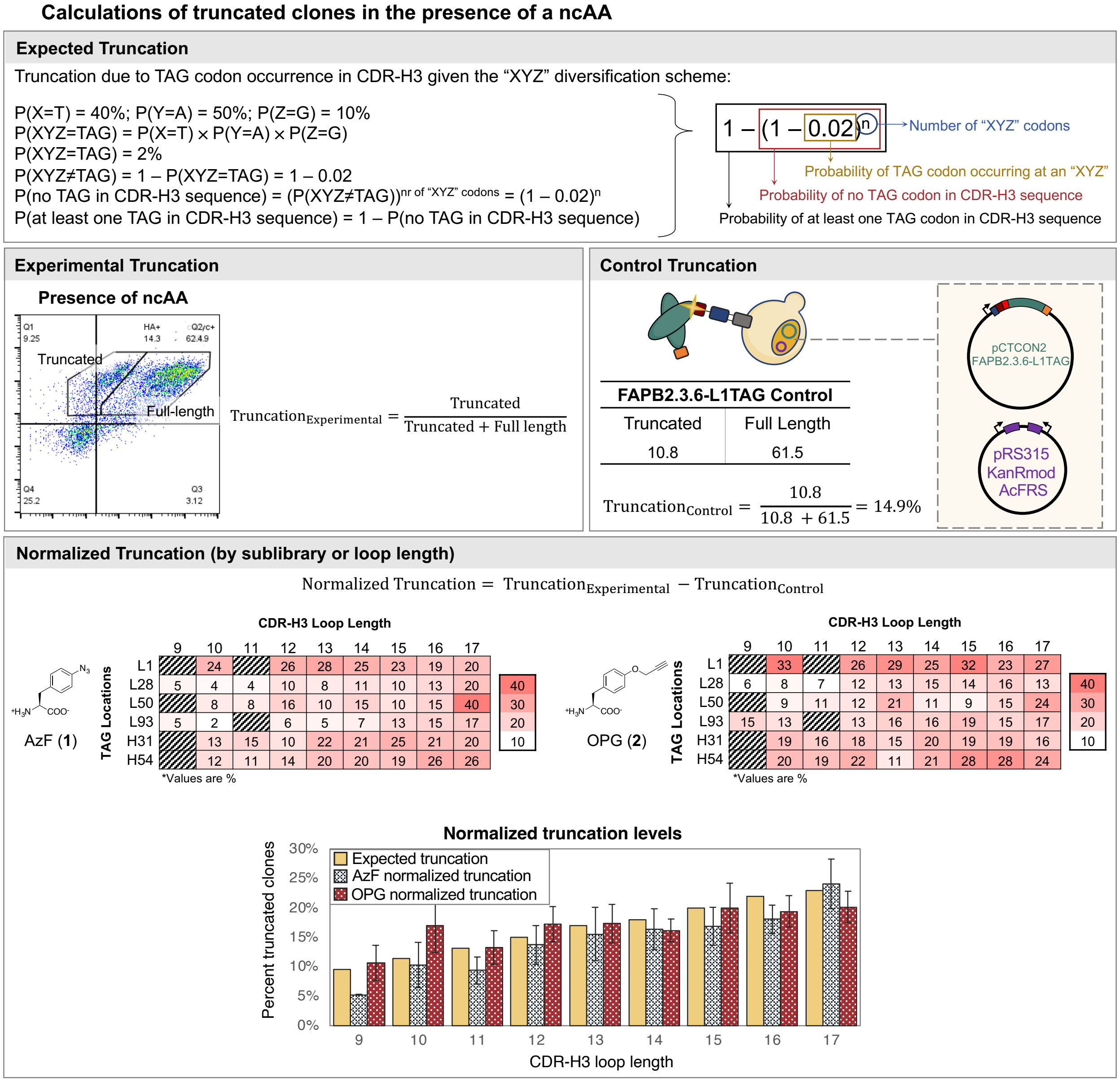


AzF (1) — CDR-H3 Loop Length (*Values are %)

| TAG Locations | 9 | 10 | 11 | 12 | 13 | 14 | 15 | 16 | 17 |
|---|---|---|---|---|---|---|---|---|---|
| L1 | | 24 | | 26 | 28 | 25 | 23 | 19 | 20 |
| L28 | 5 | 4 | 4 | 10 | 8 | 11 | 10 | 13 | 20 |
| L50 | | 8 | 8 | 16 | 10 | 15 | 10 | 15 | 40 |
| L93 | 5 | 2 | | 6 | 5 | 7 | 13 | 15 | 17 |
| H31 | | 13 | 15 | 10 | 22 | 21 | 25 | 21 | 20 |
| H54 | | 12 | 11 | 14 | 20 | 20 | 19 | 26 | 26 |

OPG (2) — CDR-H3 Loop Length (*Values are %)

| TAG Locations | 9 | 10 | 11 | 12 | 13 | 14 | 15 | 16 | 17 |
|---|---|---|---|---|---|---|---|---|---|
| L1 | | 33 | | 26 | 29 | 25 | 32 | 23 | 27 |
| L28 | 6 | 8 | 7 | 12 | 13 | 15 | 14 | 16 | 13 |
| L50 | | 9 | 11 | 12 | 21 | 11 | 9 | 15 | 24 |
| L93 | 15 | 13 | | 13 | 16 | 16 | 19 | 15 | 17 |
| H31 | | 19 | 16 | 18 | 15 | 20 | 19 | 19 | 16 |
| H54 | | 20 | 19 | 22 | 11 | 21 | 28 | 28 | 24 |

**SI Figure 4.** Calculation of truncated clones in the presence of a ncAA and comparisons between expected and normalized experimental truncation. Expected truncation (top) is calculated based on the probability of at least one TAG codon being present in CDR-H3 sequence according to the design. The experimental truncation (middle) is calculated based on the populations that fall in the truncated and full-length gates. The normalized truncation (bottom) is calculated in each sublibrary based on the experimental truncation and control truncation. A comparison of the expected and normalized truncations when the sublibraries are induced with AzF and OPG show expected trends in truncated clones.

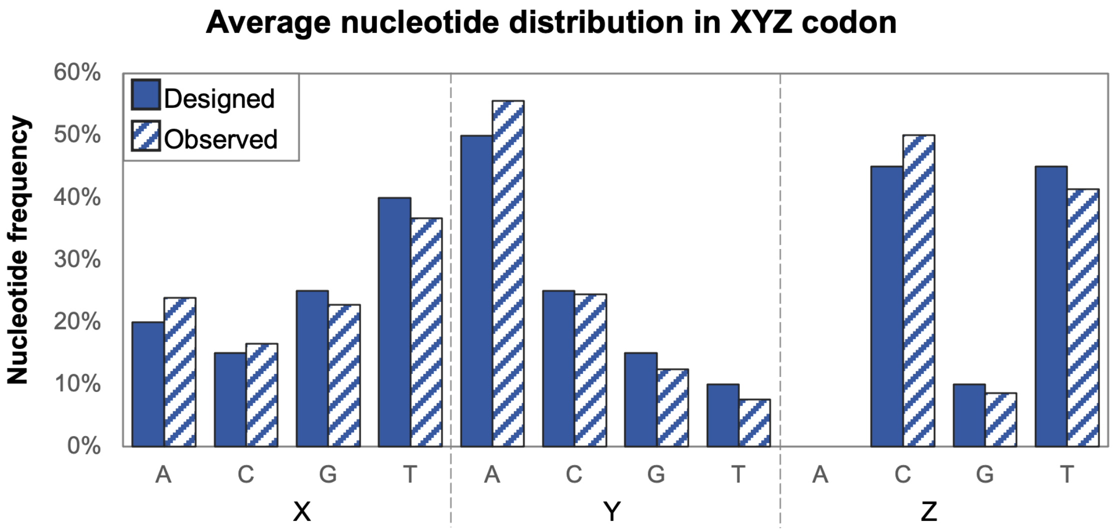


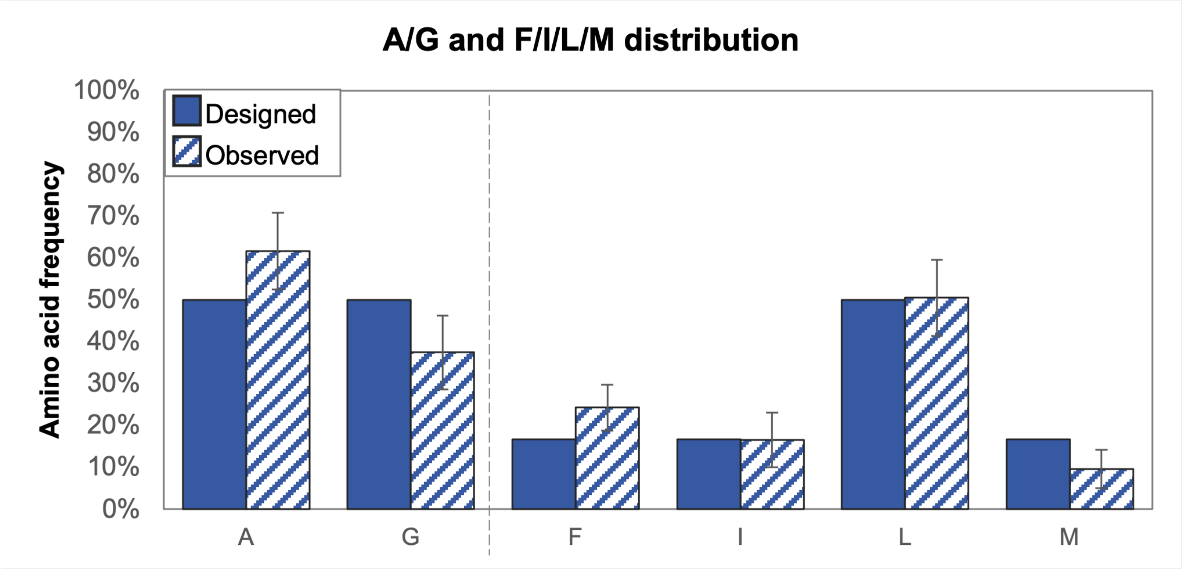


**SI Figure 5.** Sequence analysis of library clones showed the observed average nucleotide distribution in XYZ codon followed the design (top). Amino acid analysis from the sequence verified clones showed the observed frequencies of A/G and F/I/L/M distribution to also follow the design (bottom).

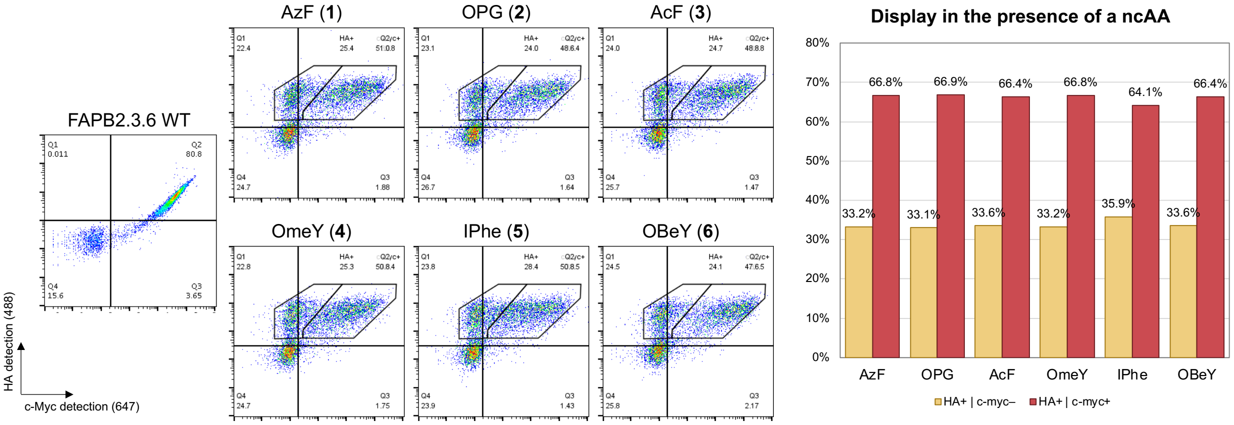


**SI Figure 6.** Pooled library exhibits similar incorporation of all six ncAAs, as observed in the flow cytometry plots. The fraction of truncated (yellow) and full-length scFv (red), quantified from the flow cytometry data, indicates efficient incorporation of the six ncAAs.

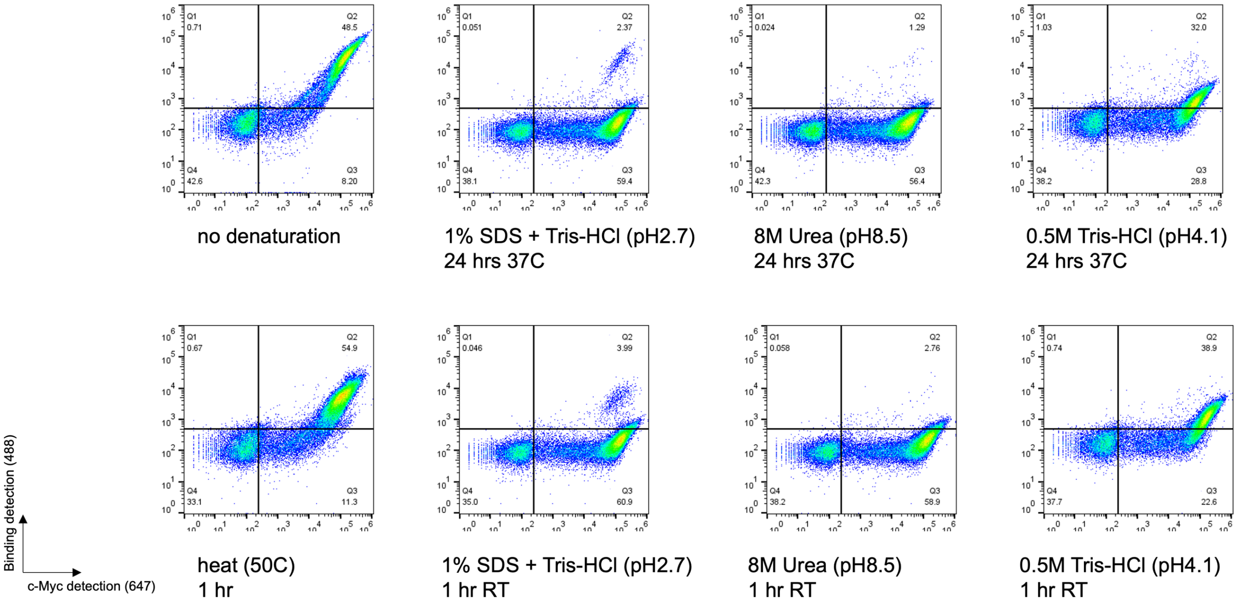


**SI Figure 7.** Flow cytometry results of donkey IgG binding retention to the clone Donkey1.1 following incubation with 200 nM donkey IgG and denaturation with various denaturing conditions and buffers as listed in the figure.

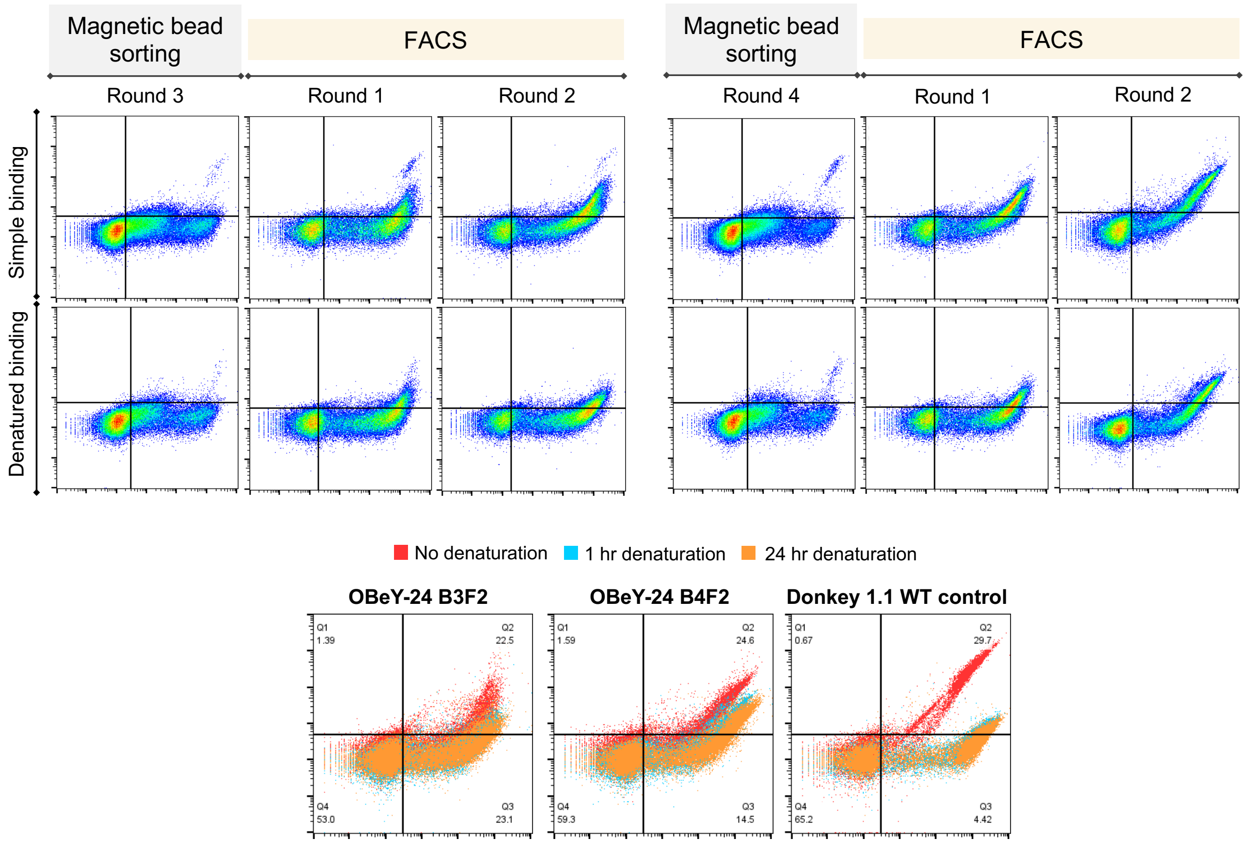


**SI Figure 8.** Flow cytometry analysis of retained binding to OBeY-substituted OBeY-24 populations with 200 nM biotinylated donkey IgG under simple binding conditions (4 hours binding at 37 °C; no denaturation) or following denaturation (denaturation with 8 M urea, 200 mM EDTA, 25 mM tris, pH 8.4 for 1 hour).

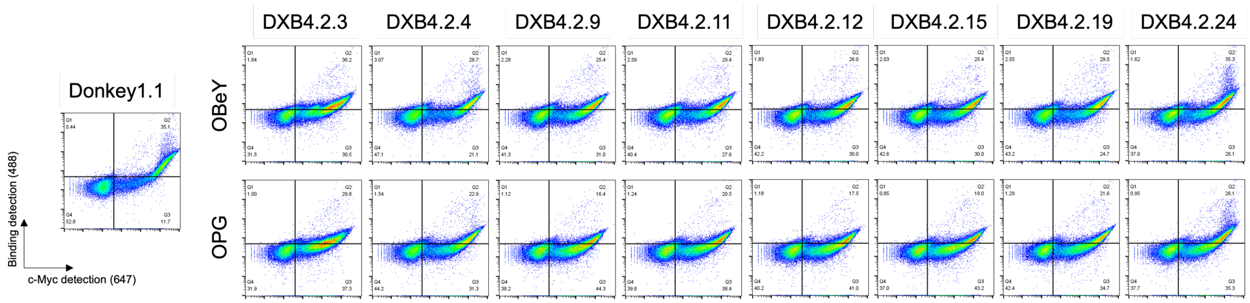


**SI Figure 9.** Flow cytometry plots of single clone binding to 200 nM native form of donkey IgG for 30 minutes at room temperature.

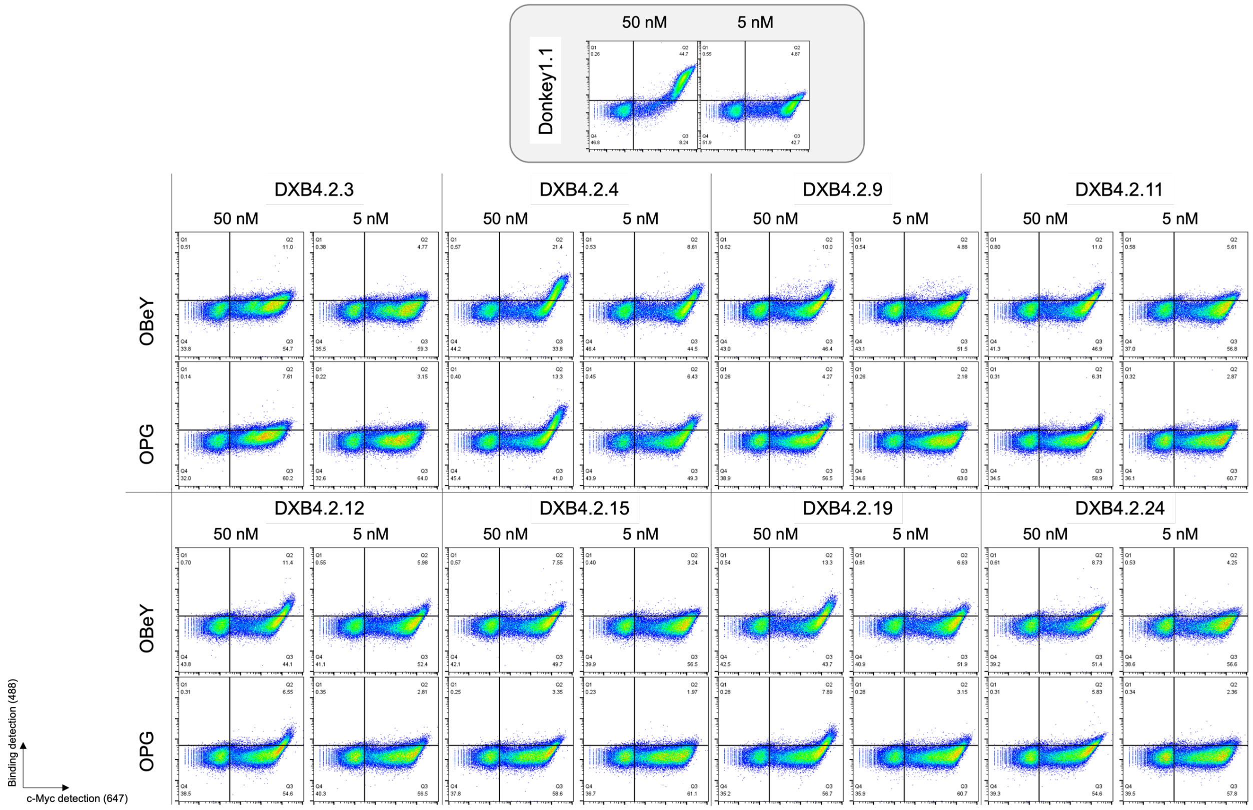


**SI Figure 10.** Flow cytometry plots of single clones binding to 50 nM and 5 nM biotinylated donkey IgG for 30 minutes at room temperature.

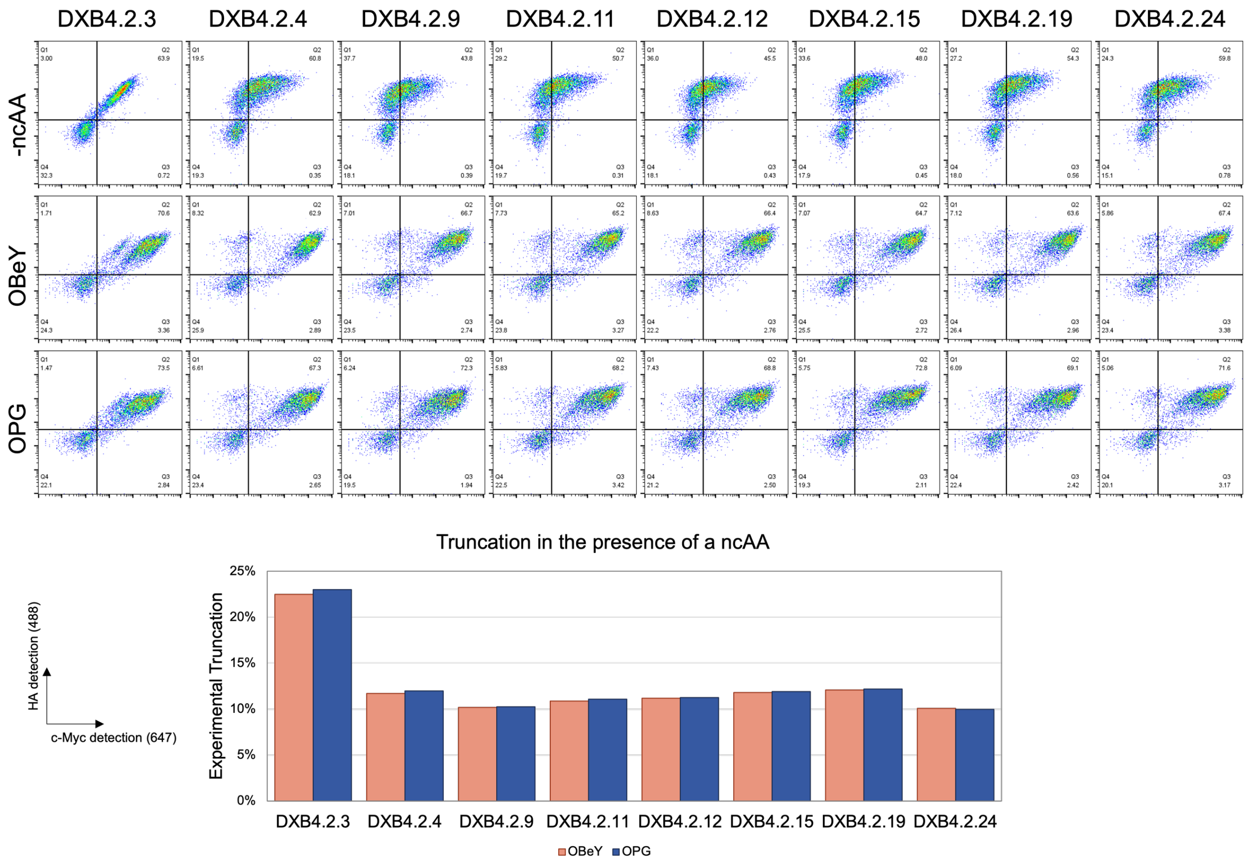


**SI Figure 11.** Full-length display in the absence and presence of a ncAA for the single clones, and truncation analysis in the presence of each ncAA.

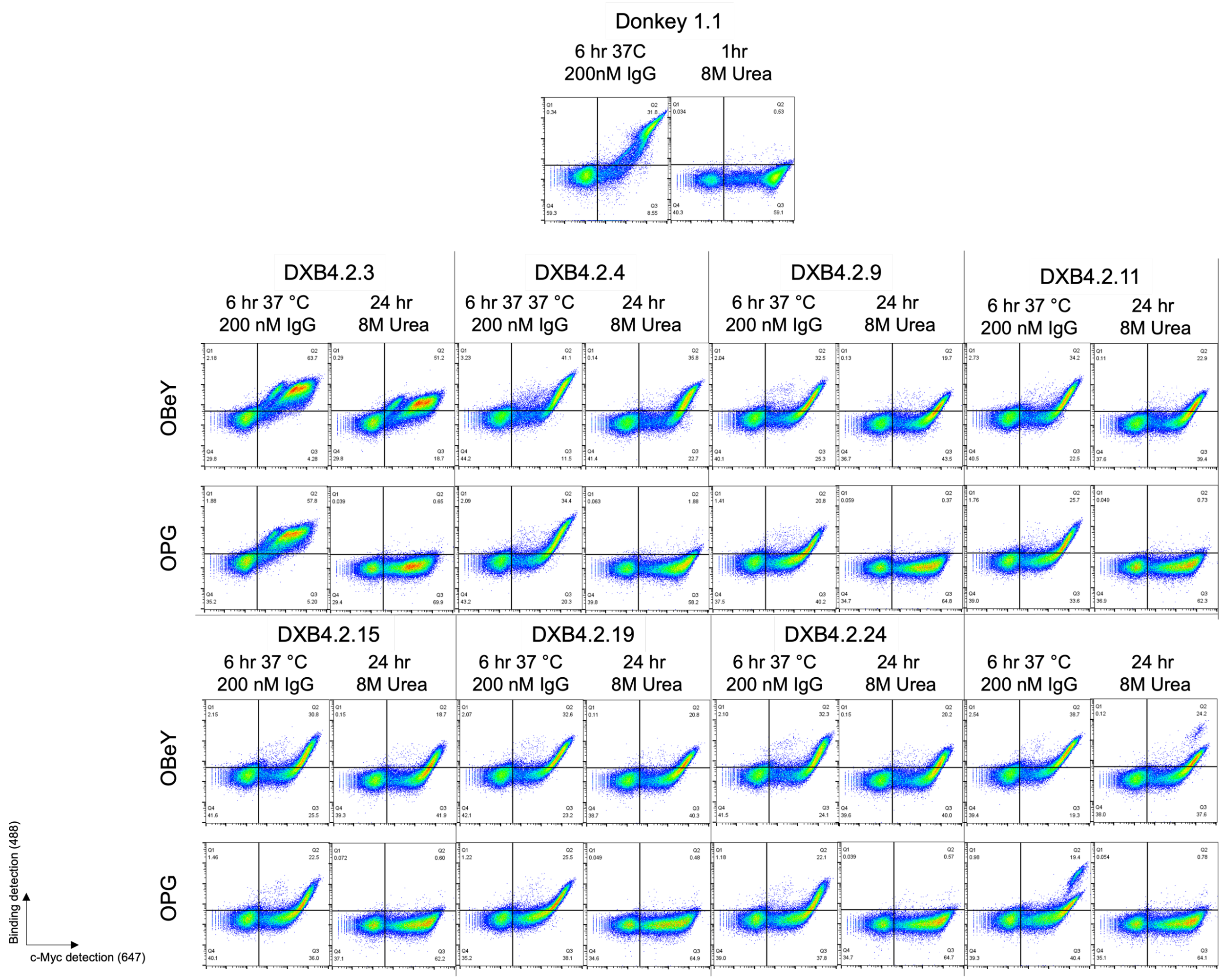


**SI Figure 12.** Flow cytometry plots corresponding to data shown in main text figure 3.4A.

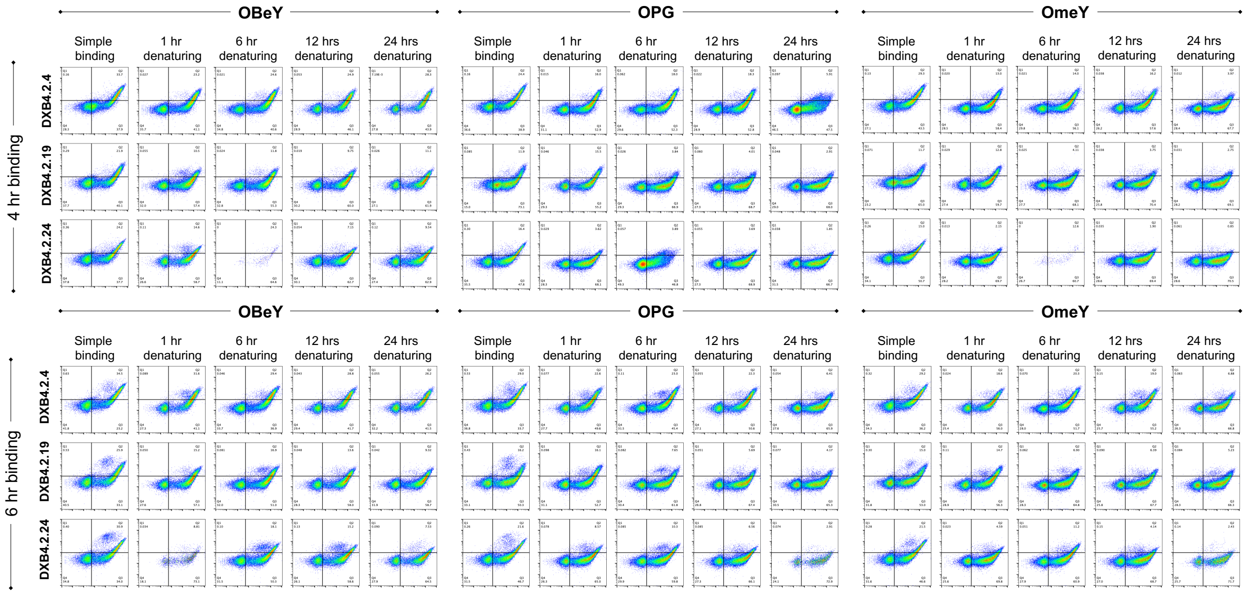


**SI Figure 13.** Flow cytometry analysis of single clones binding to 200 nM biotinylated donkey IgG under simple binding conditions (no denaturation) or under various denaturation time points with 8 M urea, 200 mM EDTA, 25 mM tris, pH 8.4.

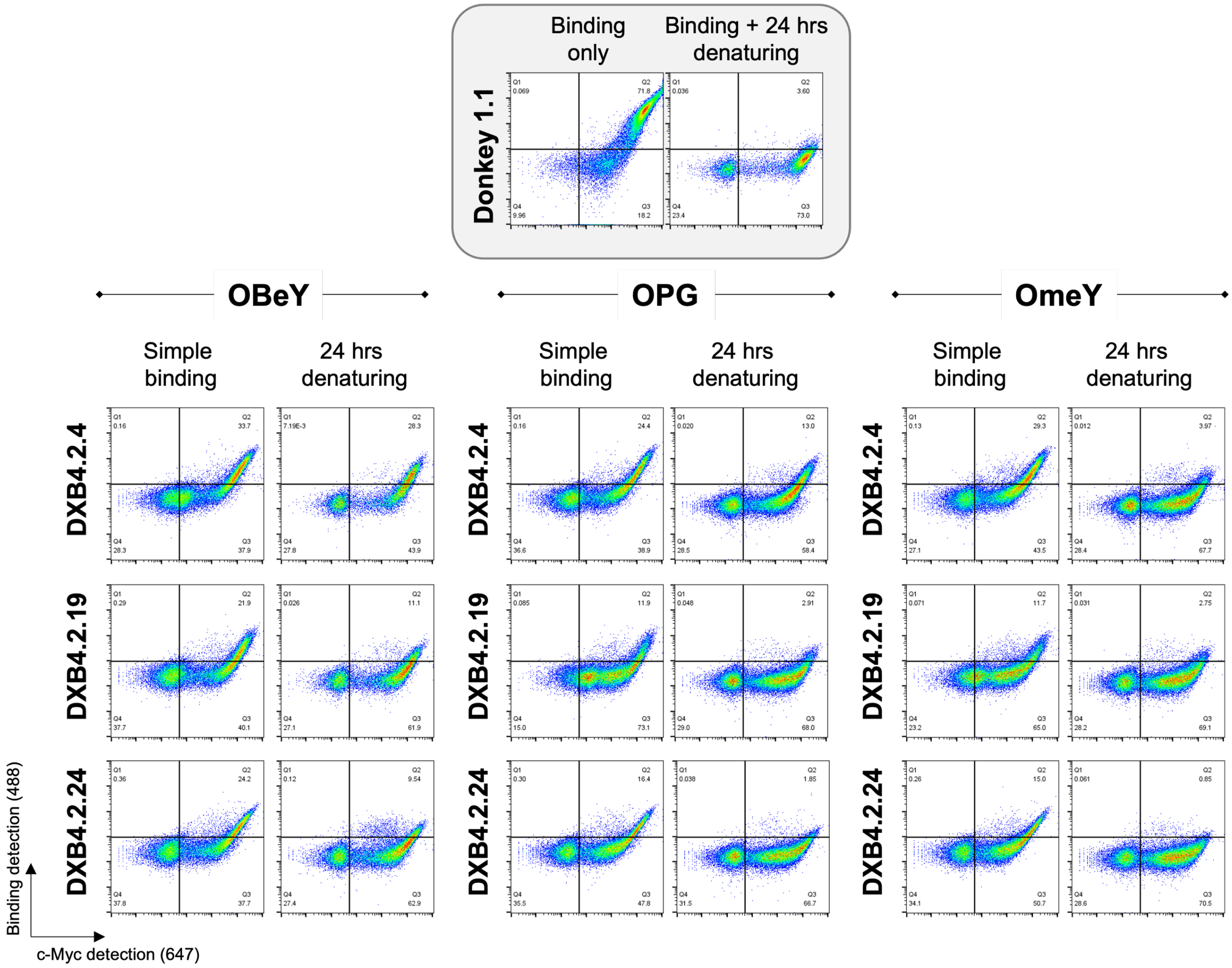


**SI Figure 14.** Flow cytometry plots of single clones binding to 200 nM biotinylated donkey IgG for 6 hours at 37 °C followed by 24 h denaturing with 8 M urea (pH 8.4, EDTA 200 mM, tris 25 mM).

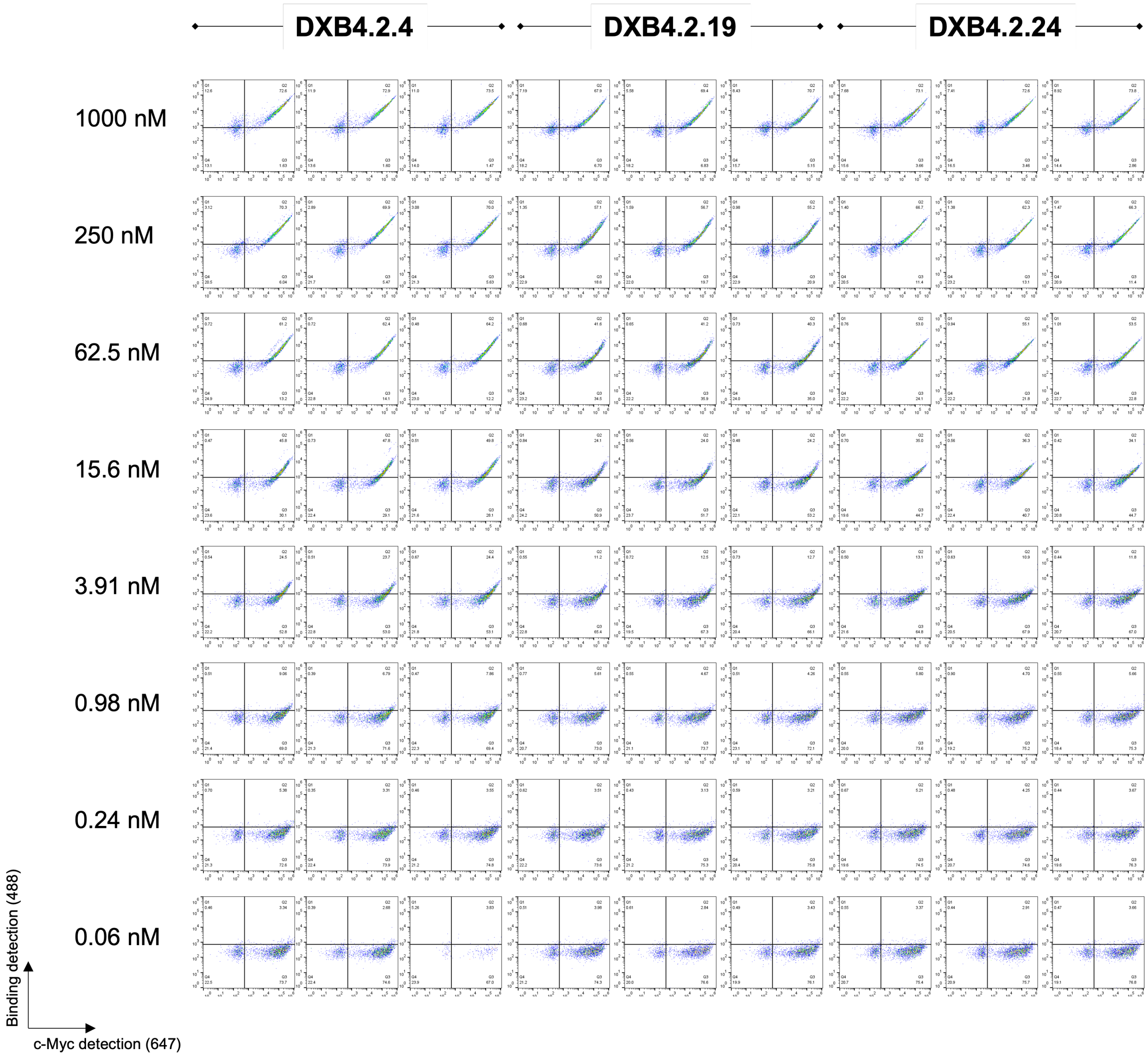


**SI Figure 15.** Flow cytometry data used to calculate $K_D$ values for single clone binding to titrated concentration of biotinylated donkey IgG.

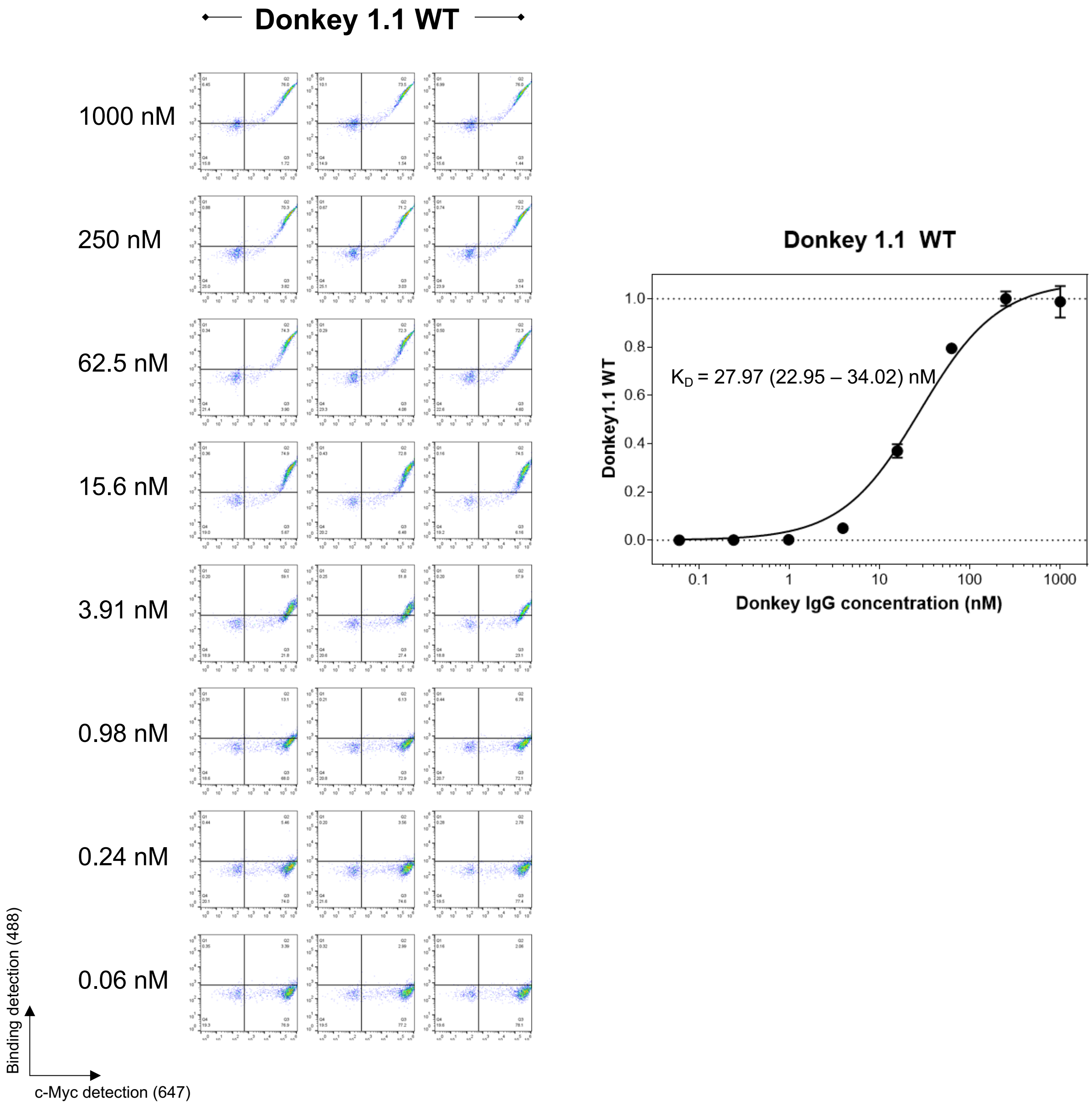


**SI Figure 16.** Flow cytometry data used to calculate $K_D$ values for Donkey1.1 WT binding to titrated concentration of biotinylated donkey IgG.

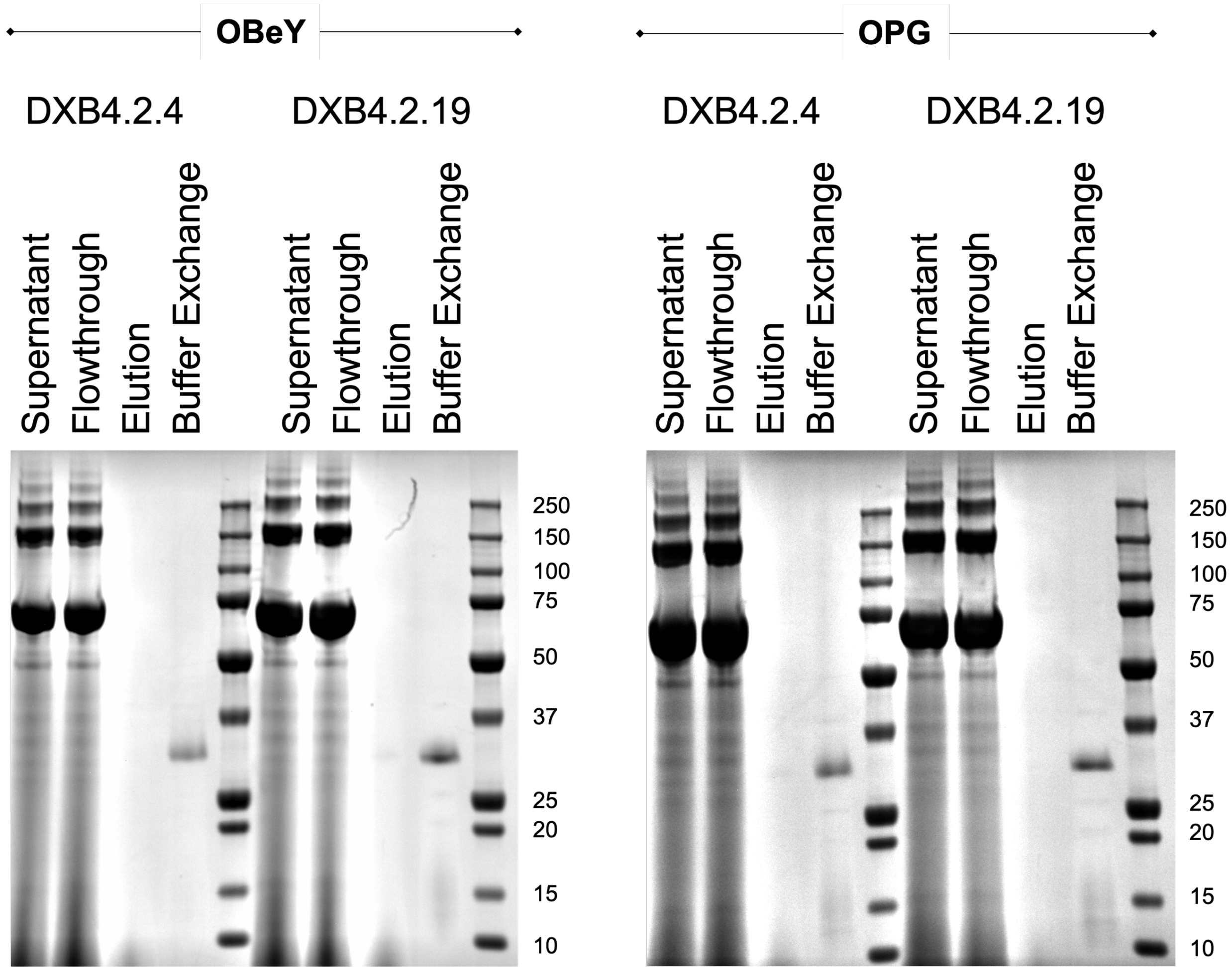


**SI Figure 17.** SDS-PAGE validation of purified scFv proteins with OBeY and OPG substituted positions.

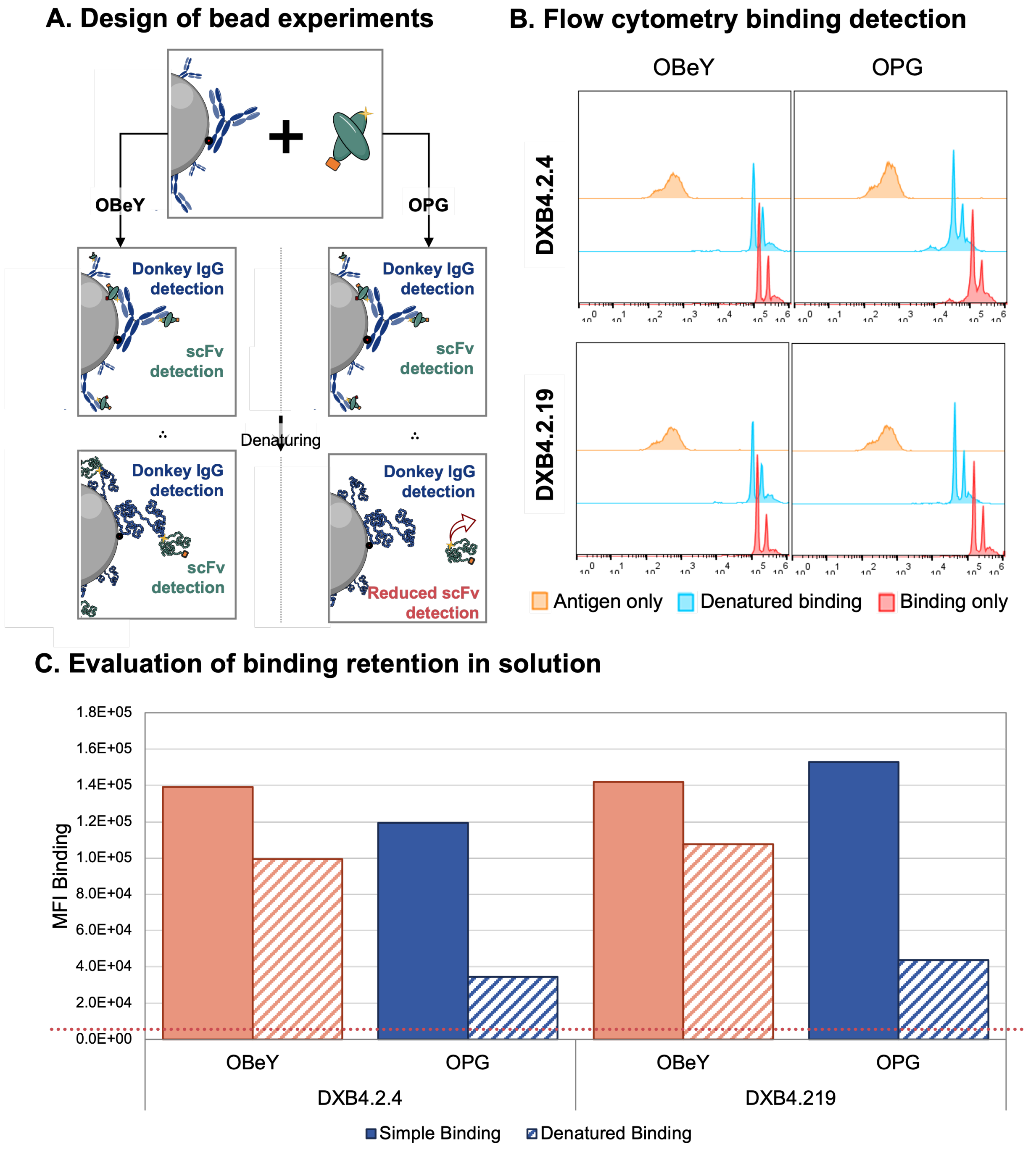


**SI Figure 18.** Binding evaluation and crosslinking of single clones in solution. (A) Experimental setup for the evaluation of irreversible bond formation via flow cytometry involves antigen coated beads and soluble ncAA-containing scFvs. (B) Flow cytometry detection of binders to the IgG coated beads under binding only and denaturing conditions. (C) MFI evaluation of data shown in panel B. Dotted line represents background MFI levels.

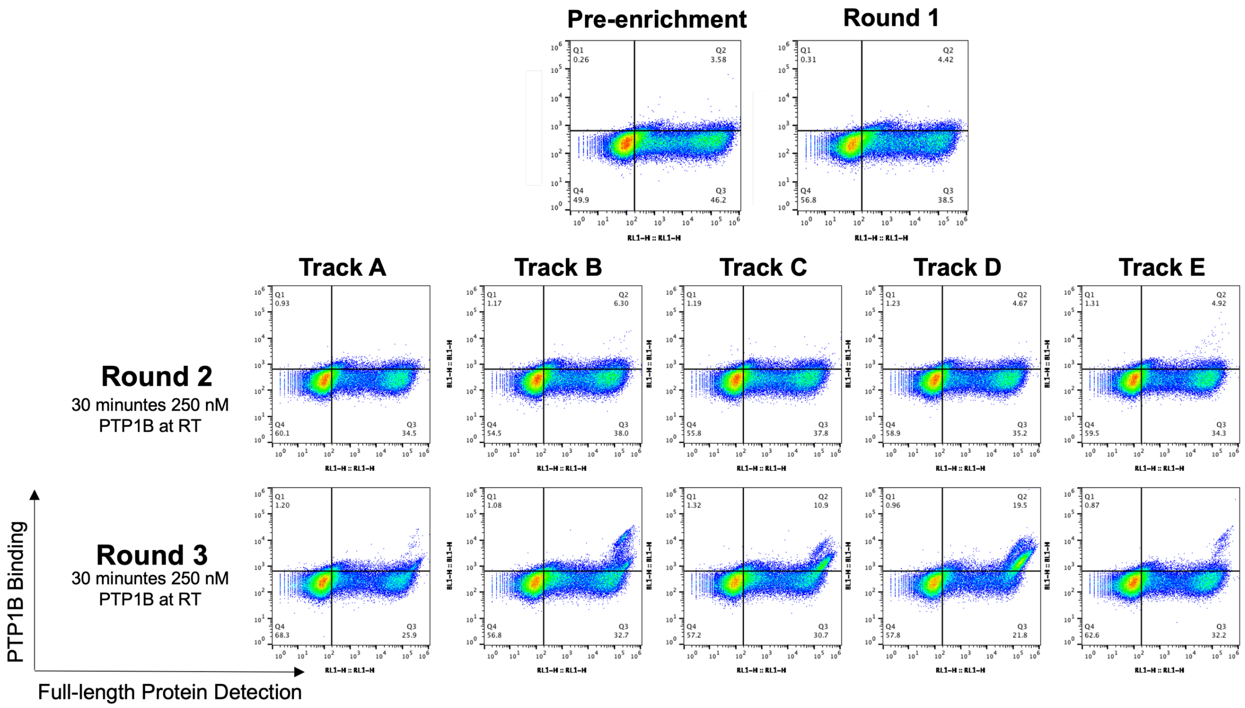


**SI Figure 19.** Flow cytometry plots monitoring PTP1B binding for each round of enrichment. Samples were treated with 250 nM PTP1B for 30 minutes at room temperature.

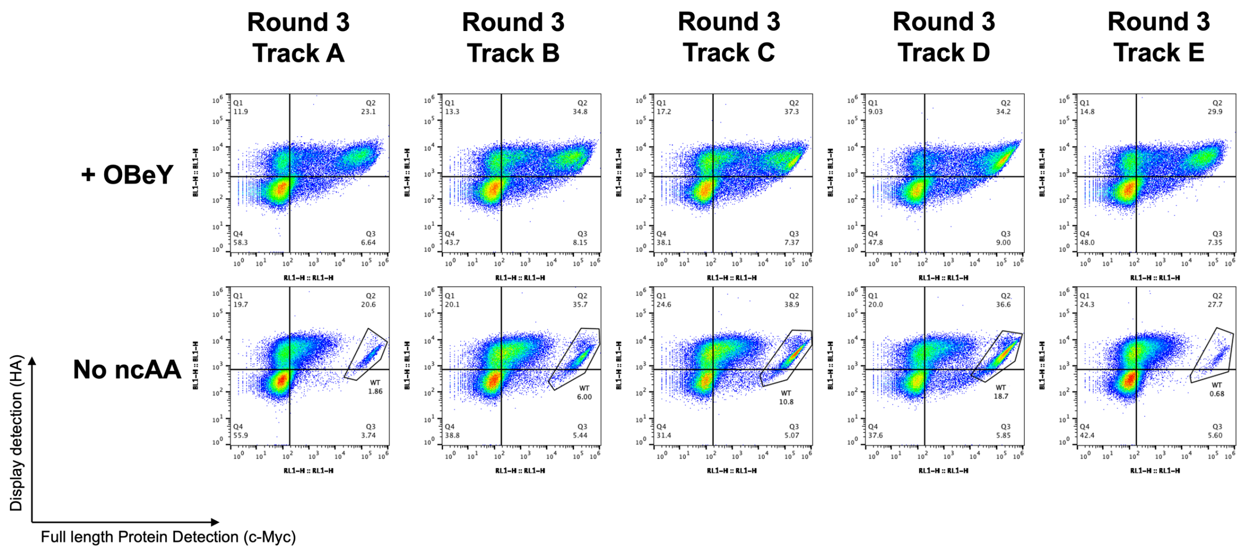


**SI Figure 20.** Each track of round 3 PTP1B bead enrichments was induced in the presence of 1 mM OBeY as well as in the absence of any ncAA to determine if constructs lacking TAG codons have been enriched. Cells were labelled for HA (display) and c-Myc (full length protein) detection for analysis. The gate labelled “WT” indicates the percent of the population showing full length protein readthrough in the absence of ncAA, and thus the apparent percentage of “wildtype” constructs in each population.

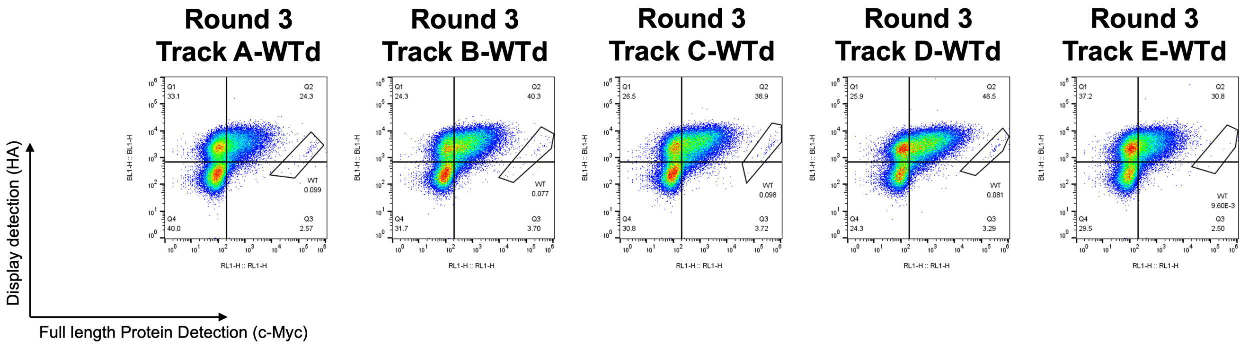


**SI Figure 21.** Each track of round 3 PTP1B enrichments following wildtype depletion (WTd) was induced in the absence of any ncAA to determine if wildtype constructs have been eliminated. Cells were labelled for HA (display) and c-Myc (full length protein) detection for analysis. The gate labelled “WT” indicates the percent of the population showing full length protein in the absence of ncAA incorporation, and thus the apparent percentage of wildtype constructs in each population.

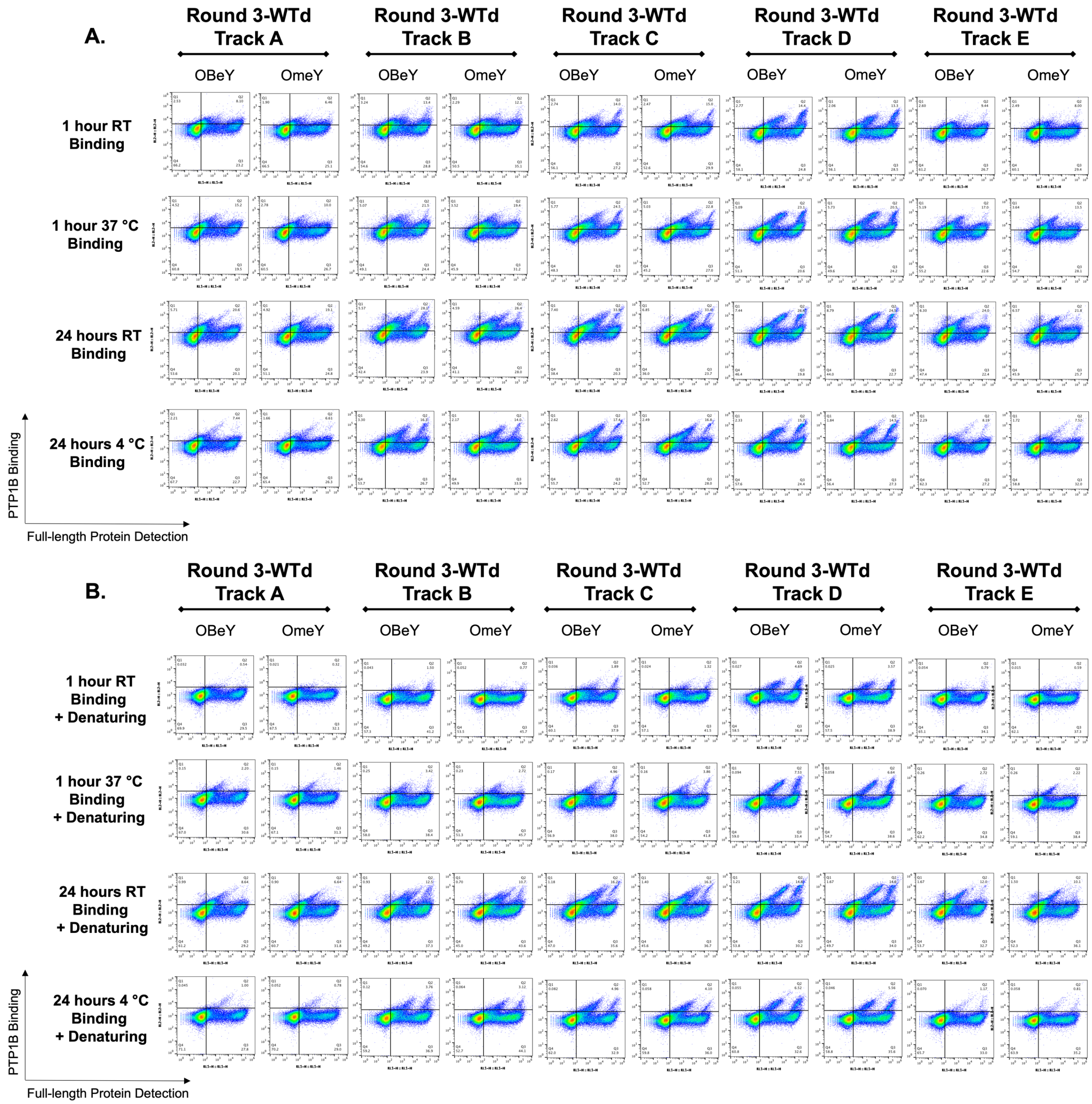


**SI Figure 22.** A) Binding analysis of wildtype-depleted PTP1B enrichment populations at indicated temperatures for either 24 hours or 1 hour. Samples were incubated with 250 nM PTP1B for each condition. B) Samples from A subjected to denaturing conditions: 1 hour room temperature incubation with 8 M urea, 200 mM EDTA, 25 mM tris HCl, pH 8.5.

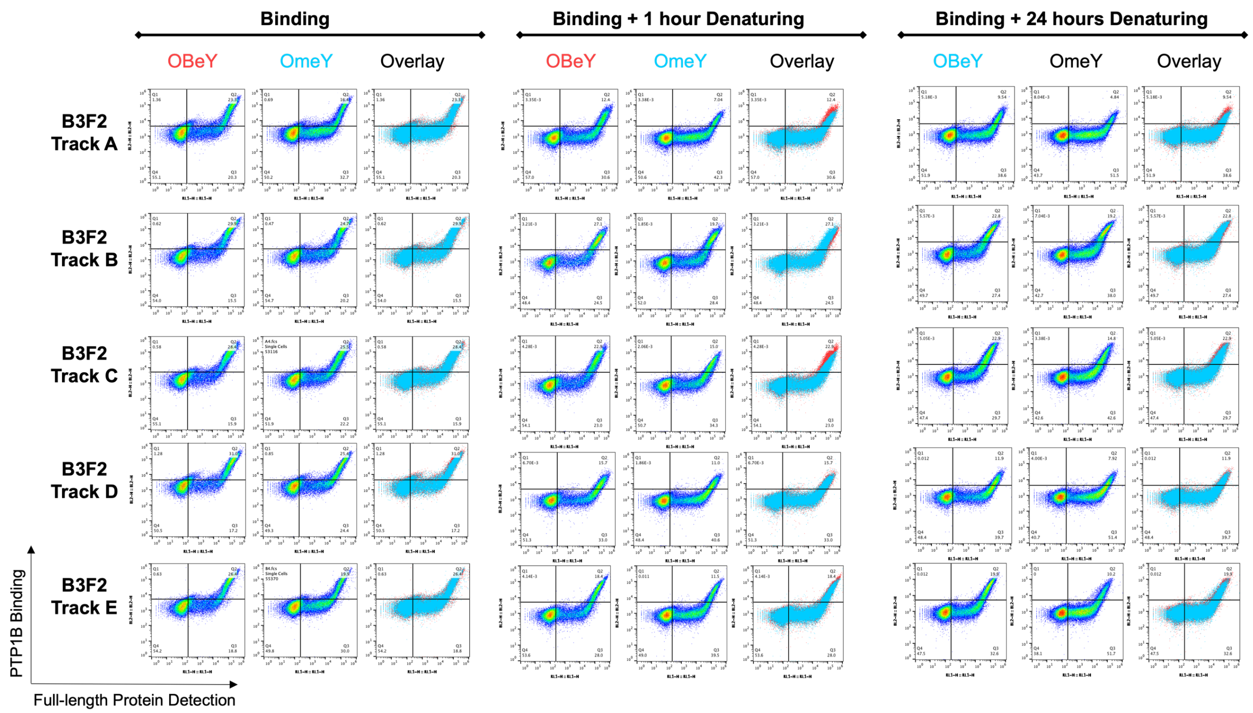


**SI Figure 23.** Analysis of B3F2 sorting tracks (3 rounds of bead sorting and 2 rounds of FACS). Samples were incubated with 250 nM PTP1B for 24 hours at 4 °C and subjected to either 1 hour or 24 hours of denaturing with 8 M urea, 200 mM EDTA, 25 mM tris-HCl, pH 8.5.

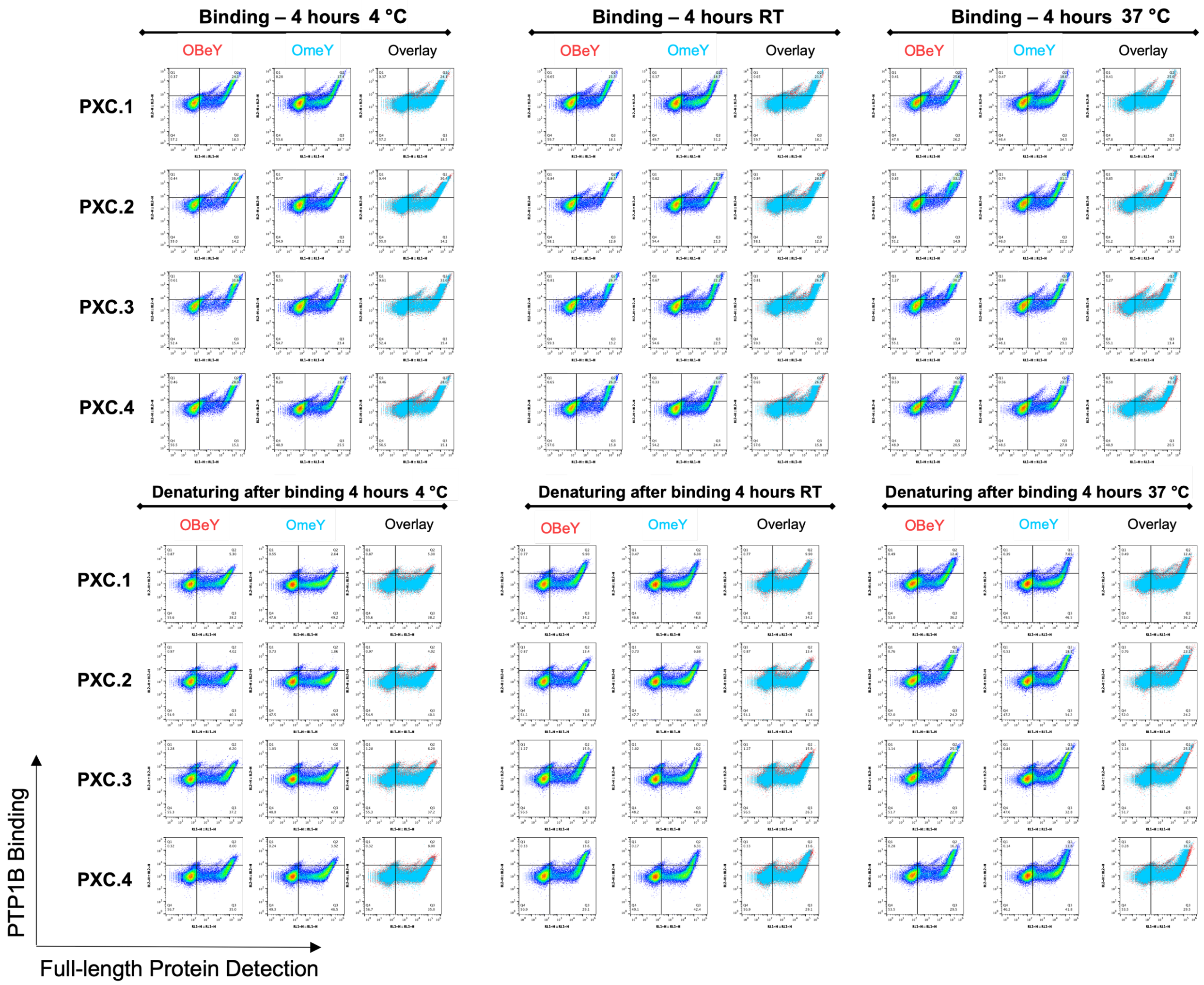


**SI Figure 24.** Individual clones isolated from library populations were subjected to analysis with 250 nM PTP1B for 4 hours at 4 °C, room temperature, and 37 °C (top). Following binding, samples were also subjected to denaturing for analysis with 8 M urea, 200 mM EDTA, 25 mM tris-HCl, pH 8.4 overnight (bottom).

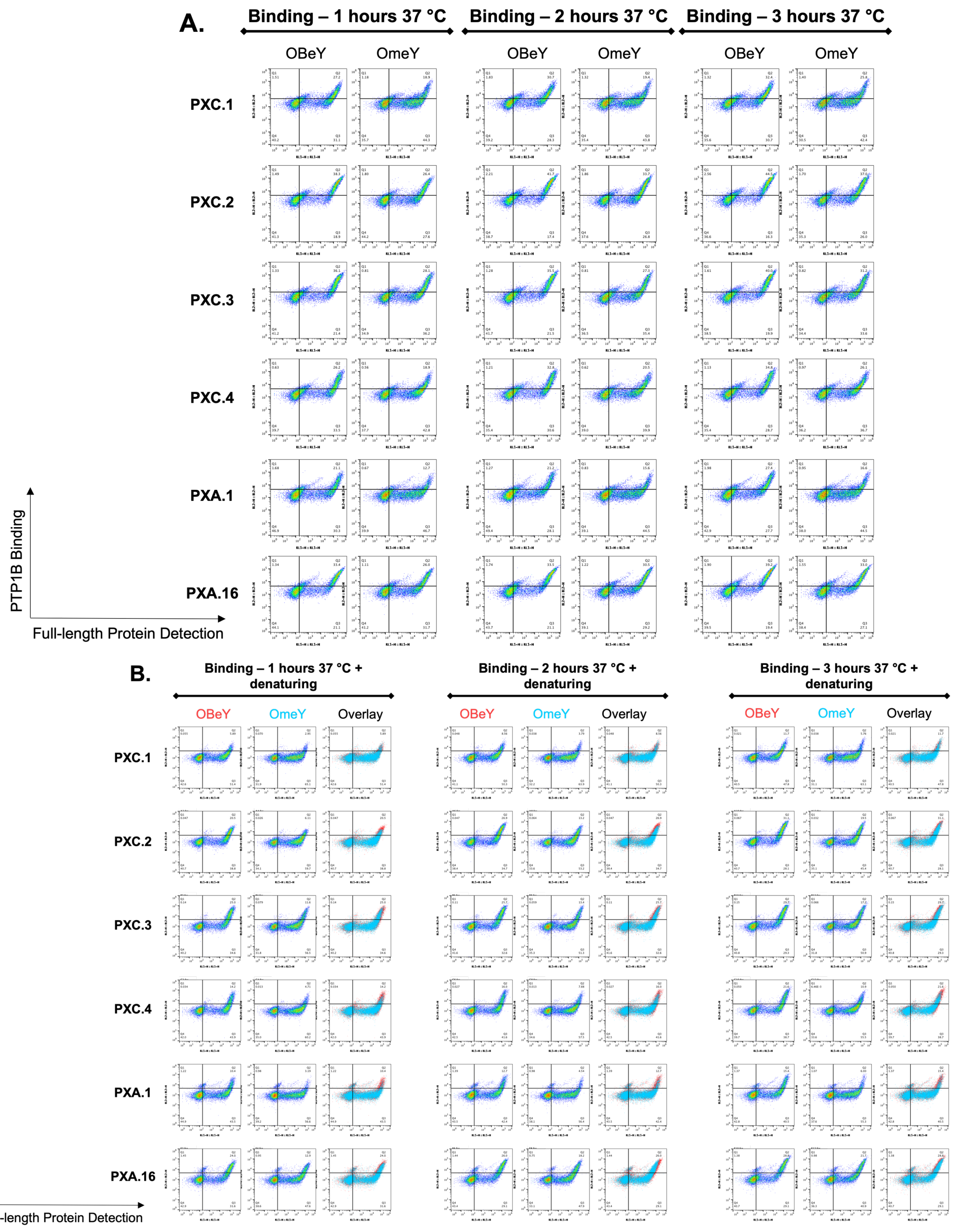


**SI Figure 25**. A) Individual PT1B binding clones incubated at 37 °C with 250 nM PTP1B for indicated time. B) Samples from A subjected to denaturing conditions: 24-hour room temperature incubation with 8 M urea, 200 mM EDTA, 25 mM tris HCl, pH 8.5.

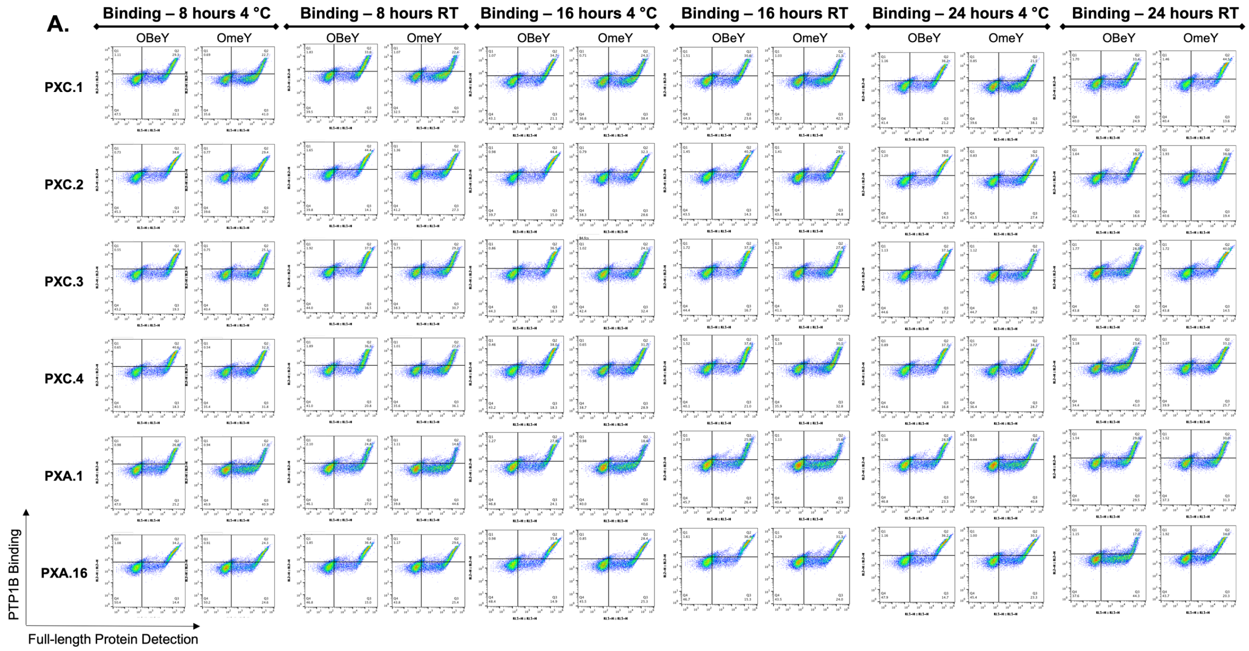


**SI Figure 26**. A) Individual PTP1B binding clones incubated for 8, 16, and 24 hours at either 4 °C or room temperature with 250 nM PTP1B. B) Samples from panel A subjected to denaturing conditions: 24-hour room temperature incubation with 8 M urea, 200 mM EDTA, 25 mM tris HCl, pH 8.5 (data on page below).

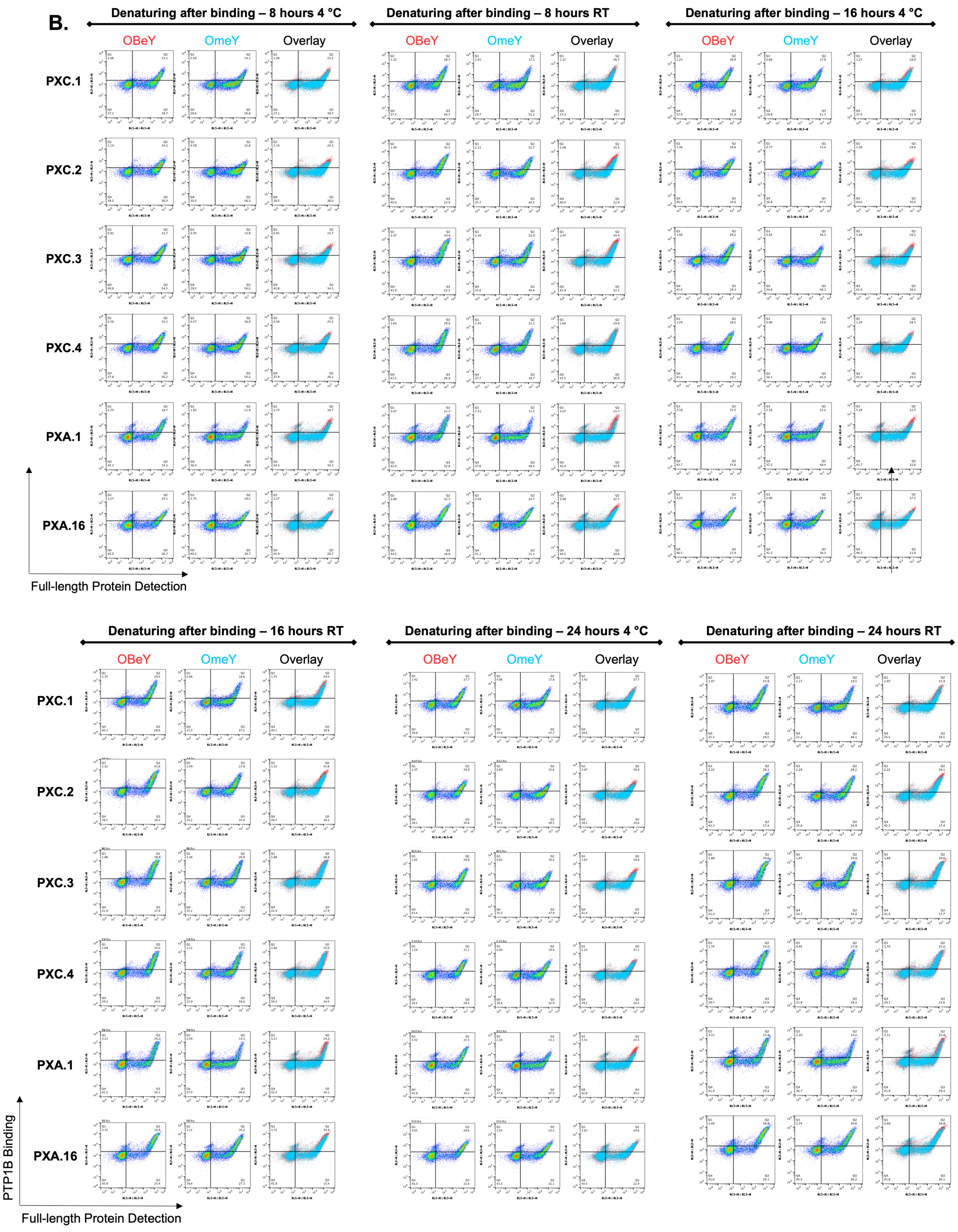


**SI Figure 26** continued.

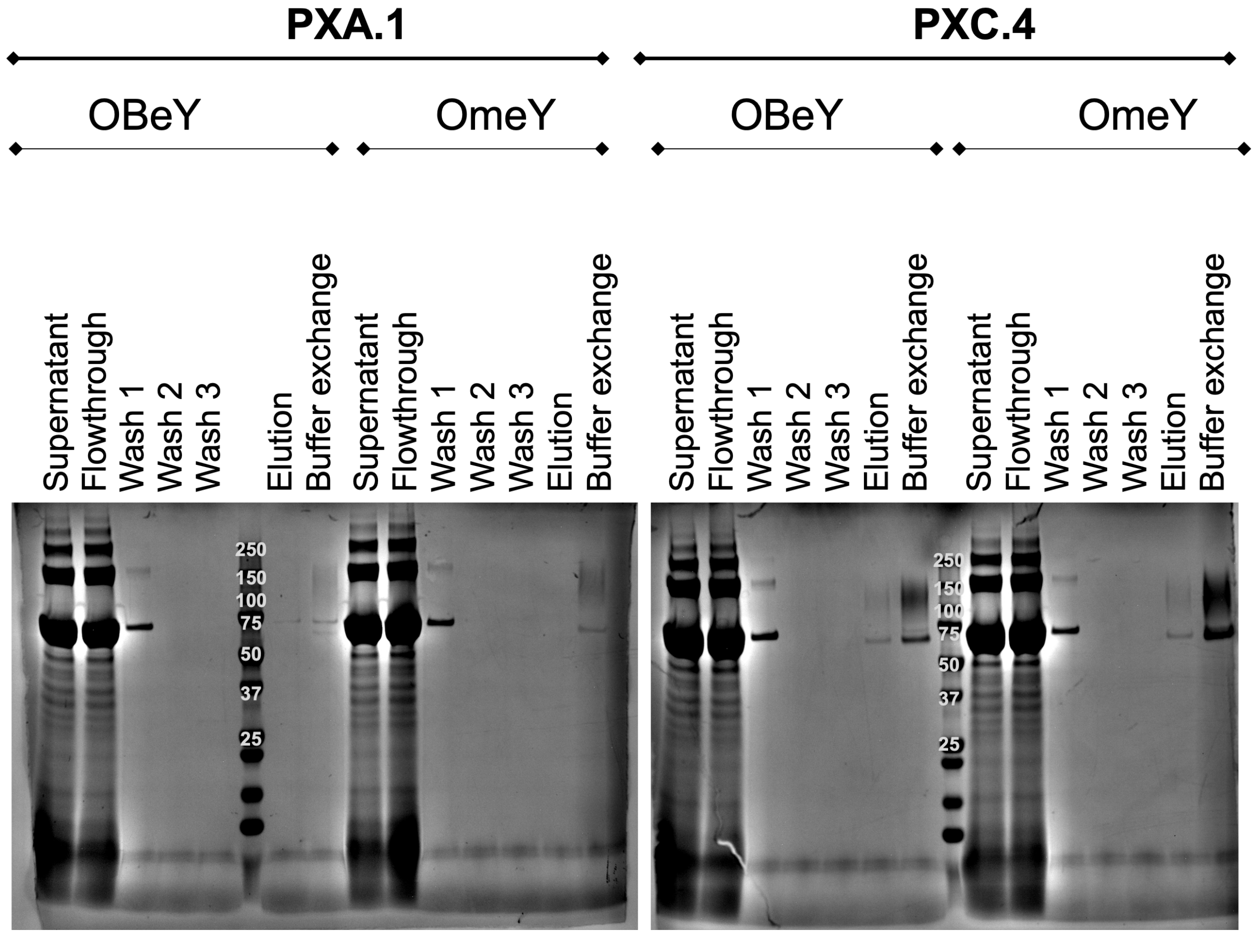


**SI Figure 27.** SDS-PAGE validation of purified PTP1B binding scFv-Fc constructs with OBeY or OmeY substitutions.

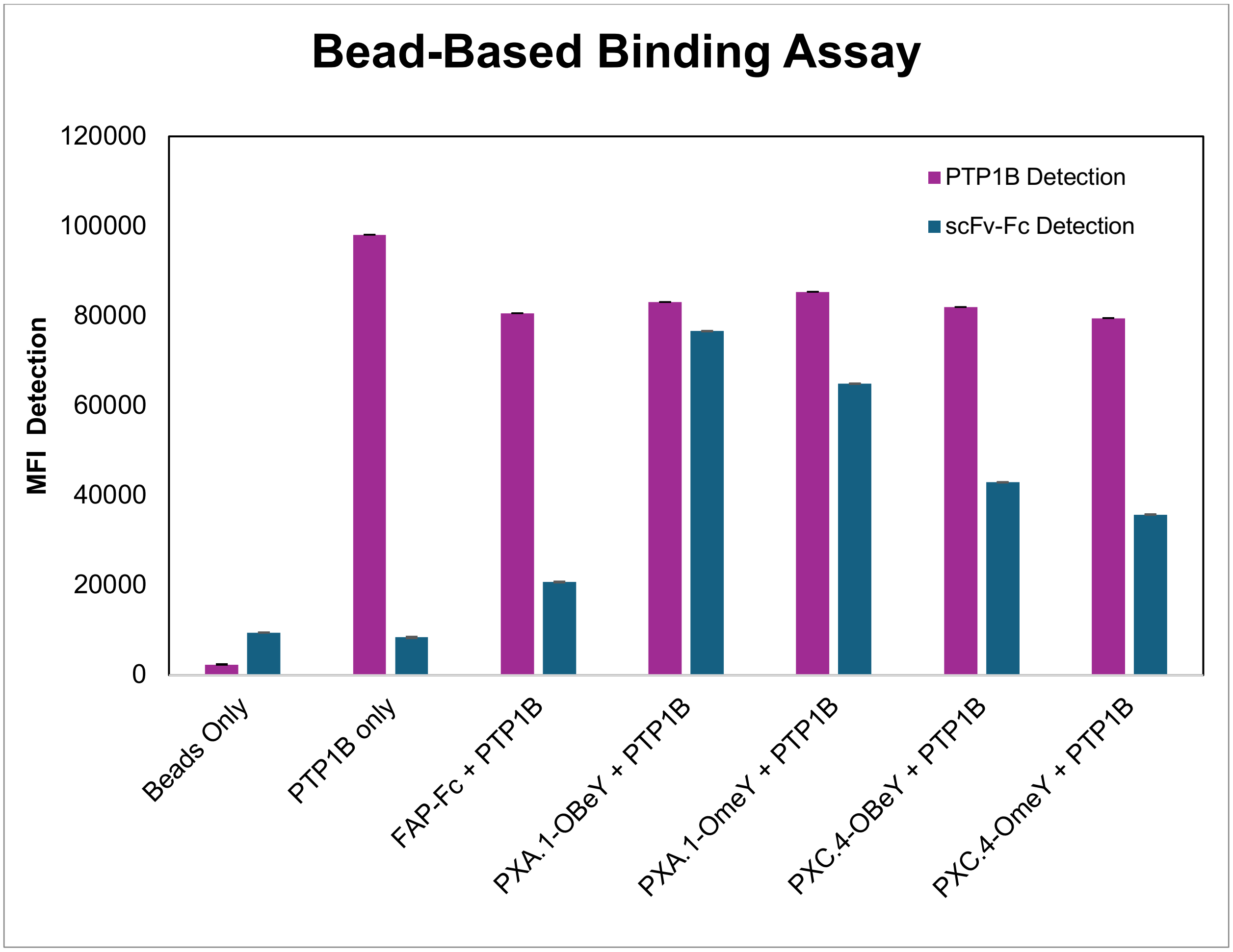


**SI Figure 28.** Results of PTP1B bead-based capture assay. Biotinylated PTP1B was immobilized on streptavidin coated beads and subsequently incubated with 150 nM scFv-Fc for 3 hours at 4 °C. Beads were labelled for biotin detection (PTP1B) and HA detection (scFv-Fcs) to show in solution binding of PXA.1 and PXC.4 to PTP1B.

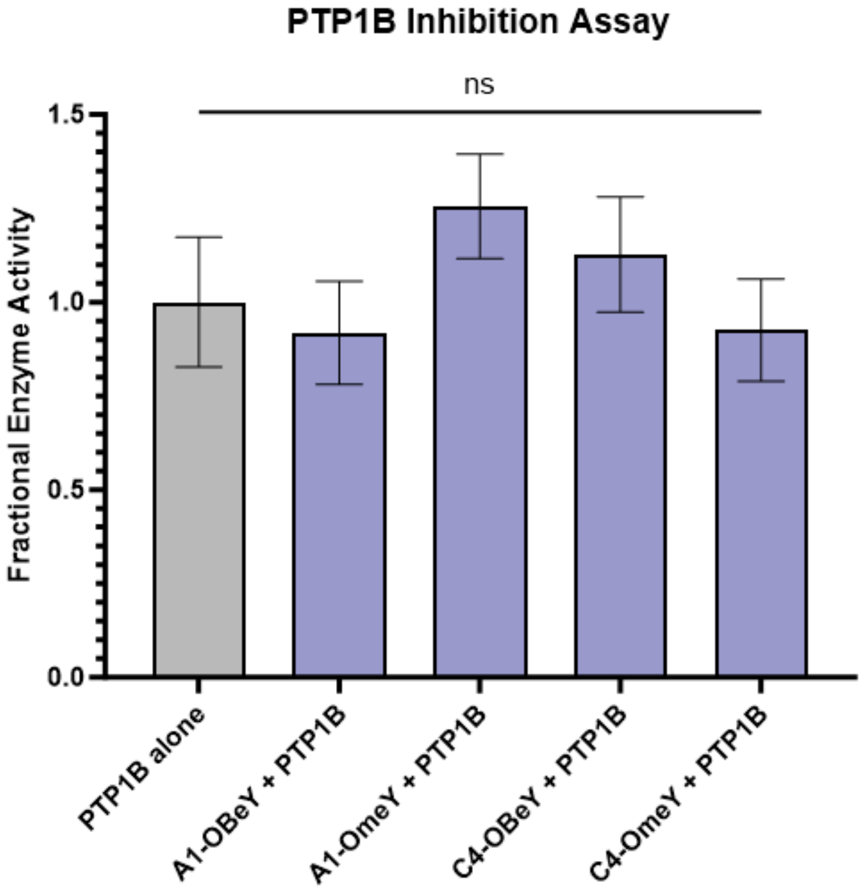


**SI Figure 29.** 50 nM PTP1B was incubated with 1 µM each scFv-Fc for 30 minutes at room temperature, and an additional sample of 50 nM PTP1B alone was used as a control. PTP1B activity was measured in all samples following incubation, and the fractional enzyme activity is compared in the graph shown here. A one-way analysis of variance (ANOVA) was used to determine statistical significance.

**SI Table 1.** Sequence validation of sublibraries.

| | Sublibrary by looplength | Clones sequenced | Unique clones | Clones with TAG in CDR-H3 | Unique clones with correct design | Clones with indels | Frameshift mutations | Clones with missing TAG |
|---|---|---|---|---|---|---|---|---|
| L1 | 9 | 5 | 4 | 0 | 1 | 2 | 1 | 1 |
| | 10 | 6 | 6 | 0 | 6 | 0 | 0 | 0 |
| | 11 | 6 | 6 | 2 | 3 | 1 | 2 | 0 |
| | 12 | 7 | 7 | 0 | 7 | 0 | 0 | 0 |
| | 13 | 6 | 6 | 0 | 5 | 0 | 1 | 0 |
| | 14 | 7 | 7 | 2 | 6 | 1 | 1 | 0 |
| | 15 | 7 | 7 | 3 | 6 | 1 | 1 | 0 |
| | 16 | 6 | 6 | 3 | 5 | 0 | 1 | 0 |
| | 17 | 6 | 6 | 1 | 5 | 0 | 1 | 1 |
| L28 | 9 | 6 | 6 | 0 | 5 | 1 | 0 | 0 |
| | 10 | 5 | 5 | 0 | 5 | 0 | 0 | 0 |
| | 11 | 5 | 5 | 0 | 5 | 0 | 0 | 0 |
| | 12 | 5 | 5 | 0 | 5 | 0 | 0 | 0 |
| | 13 | 5 | 5 | 0 | 5 | 0 | 0 | 0 |
| | 14 | 5 | 5 | 2 | 5 | 0 | 0 | 0 |
| | 15 | 8 | 8 | 0 | 7 | 1 | 0 | 0 |
| | 16 | 6 | 6 | 0 | 5 | 1 | 1 | 0 |
| | 17 | 7 | 7 | 0 | 6 | 0 | 0 | 0 |
| L50 | 9 | 6 | 6 | 0 | 5 | 1 | 0 | 0 |
| | 10 | 5 | 5 | 0 | 5 | 0 | 0 | 0 |
| | 11 | 5 | 5 | 1 | 5 | 0 | 0 | 0 |
| | 12 | 5 | 5 | 0 | 5 | 0 | 0 | 0 |
| | 13 | 5 | 5 | 1 | 5 | 0 | 0 | 0 |
| | 14 | 6 | 6 | 2 | 5 | 0 | 1 | 0 |
| | 15 | 7 | 7 | 1 | 6 | 1 | 0 | 0 |
| | 16 | 6 | 6 | 0 | 5 | 1 | 0 | 0 |
| | 17 | 6 | 6 | 0 | 6 | 0 | 0 | 0 |
| L93 | 9 | 5 | 5 | 0 | 5 | 0 | 0 | 0 |
| | 10 | 6 | 6 | 0 | 6 | 0 | 0 | 0 |
| | 11 | 6 | 5 | 0 | 4 | 1 | 0 | 0 |
| | 12 | 6 | 6 | 1 | 6 | 0 | 0 | 0 |
| | 13 | 5 | 5 | 0 | 5 | 0 | 0 | 0 |
| | 14 | 6 | 6 | 1 | 5 | 1 | 0 | 0 |
| | 15 | 6 | 6 | 2 | 5 | 1 | 0 | 0 |
| | 16 | 6 | 6 | 1 | 5 | 0 | 0 | 0 |
| | 17 | 5 | 5 | 2 | 5 | 0 | 0 | 0 |
| H31 | 9 | Failed at flow cytometry check | | | | | | |
| | 10 | 5 | 5 | 1 | 5 | 0 | 0 | 0 |
| | 11 | 7 | 7 | 2 | 6 | 1 | 0 | 0 |
| | 12 | 8 | 8 | 2 | 7 | 0 | 1 | 0 |
| | 13 | 6 | 6 | 0 | 5 | 0 | 1 | 0 |
| | 14 | 6 | 6 | 0 | 5 | 1 | 0 | 0 |
| | 15 | 5 | 5 | 0 | 5 | 0 | 0 | 0 |
| | 16 | 7 | 7 | 1 | 6 | 1 | 0 | 0 |
| | 17 | 6 | 6 | 2 | 5 | 0 | 1 | 0 |
| H54 | 9 | Failed at flow cytometry check | | | | | | |
| | 10 | 5 | 5 | 0 | 5 | 0 | 0 | 0 |
| | 11 | 7 | 7 | 2 | 7 | 0 | 0 | 0 |
| | 12 | 10 | 10 | 2 | 10 | 0 | 0 | 0 |
| | 13 | 7 | 7 | 2 | 6 | 0 | 1 | 0 |
| | 14 | 8 | 8 | 1 | 7 | 1 | 0 | 0 |
| | 15 | 7 | 7 | 0 | 7 | 0 | 0 | 0 |
| | 16 | 5 | 5 | 2 | 5 | 0 | 0 | 0 |
| | 17 | 8 | 8 | 1 | 8 | 0 | 0 | 0 |
| Total: | | 317 | 315 | 43 | 284 | 18 | 14 | 2 |

**SI Table 2.** Primers used for library construction.

| Primer Name | Sequence (5' to 3') |
|---|---|
| SyntheticAmpFwd | GGAGGCGGTAGCGGAGGCGGAGGGTCGGCTAGCGACATACAGATGACTCAAAGTCCCAG |
| SyntheticAmpRev | GTCCTCTTCAGAAATAAGCTTTTGTTCGGATCCTGAGGAGACGGTGACCAGGGTTCC |
| SidLinkFwd | GGTACTACTGCCGCTAGTGGTAGTAGTGGTG |
| SidLinkRev | GGCACCACTGCTACTGCCACCACTACTACCAC |
| CDR9-17Rev | CAGGGTTCCTTGGCCCCAGTAGTCSADASC(z'y'x')5-13<br>CTTAGCACAGTAGTAAACAGCAGTATCTTC |
| CON2seqFwd | GTTCCAGACTACGCTCTGCAGG |
| CON2seqRev | GATTTTGTTACATCTACACTGTTG |
| L1TAGFwd | GCGGTAGCGGAGGCGGAGGGTCGGCTAGCTAGATACAGATGACTCAAAGTCCCAGTTCACTA |
| L1TAGRev | TAGTGAACTGGGACTTTGAGTCATCTGTATCTAGCTAGCCGACCCTCCGCCTCCGCTACCGC |
| L28TAGFwd | GTCACCATTACGTGTAGAGCTTCTCAGTAGATTAGCTCGTACTTGAATTGGTATCAACAG |
| L28TAGRev | CTGTTGATACCAATTCAAGTACGAGCTAATCTACTGAGAAGCTCTACACGTAATGGTGAC |
| L50TAGFwd | GGGAAAGCTCCAAAGTTGCTGATCTATTAGGCATCTAGCTTACAAAGTGGTGTACCTTCC |
| L50TAGRev | GGAAGGTACACCACTTTGTAAGCTAGATGCCTAATAGATCAGCAACTTTGGAGCTTTCCC |
| L93TAGFwd | TTCGCCACATATTACTGCCAACAATCCTACTAGACTCCACCTACATTTGGTGGTGGCACTAAA |
| L93TAGRev | TTTAGTGCCACCACCAAATGTAGGTGGAGTCTAGTAGGATTGTTGGCAGTAATATGTGGCGAA |
| H31TAGFwd | TCTTGTGCTGCTAGTGGATTCACGTTTAGTTAGTATGCCATGTCATGGGTTAGACAAGCTCCAG |
| H31TAGRev | CTGGAGCTTGTCTAACCCATGACATGGCATACTAACTAAACGTGAATCCACTAGCAGCACAAGA |
| H54TAGFwd | GGCTTAGAATGGGTTTCTGCGATATCTGGATAGGGTGGGTCAACTTACTATGCAGATTCCGTC |
| H54TAGRev | GACGGAATCTGCATAGTAAGTTGACCCACCCTATCCAGATATCGCAGAAACCCATTCTAAGCC |

**SI Table 3.** Germline sequences used in this study. Restriction enzyme sites highlighted in yellow.

| Domain | Sequence |
|---|---|
| **VL**: IGKV1-39-JK4 | GCTAGCGACATACAGATGACTCAAAGTCCCAGTTCACTATCTGCGTC<br>TGTTGGTGATAGAGTCACCATTACGTGTAGAGCTTCTCAGTCGATTA<br>GCTCGTACTTGAATTGGTATCAACAGAAACCAGGGAAAGCTCCAAA<br>GTTGCTGATCTATGCAGCATCTAGCTTACAAAGTGGTGTACCTTCC<br>AGGTTTTCAGGCTCAGGATCTGGAACTGATTTCACACTTACCATAT<br>CATCCTTACAACCGGAAGATTTCGCCACATATTACTGCCAACAATC<br>CTACTCTACTCCACCTACATTTGGTGGTGGCACTAAAGTGGAGATT<br>AAGGGTACTACTGCCGCTAGTGGTAGTAGTGGTGGCAGTAGCAGT<br>GGTGCC |
| **VH**: IGHV3-23 | GGTACTACTGCCGCTAGTGGTAGTAGTGGTGGCAGTAGCAGTGGT<br>GCCGAGGTGCAATTGCTAGAATCAGGAGGTGGTTTGGTACAACCT<br>GGTGGTAGCTTAAGGTTGTCTTGTGCTGCTAGTGGATTCACGTTTA<br>GTAGCTATGCCATGTCATGGGTTAGACAAGCTCCAGGTAAAGGCT<br>TAGAATGGGTTTCTGCGATATCTGGATCTGGTGGGTCAACTTACT<br>ATGCAGATTCCGTCAAAGGCAGATTTACCATTTCCAGAGACAATT<br>CGAAGAATACACTGTACCTTCAGATGAACTCGTTACGTGCAGAAG<br>ATACTGCTGTTTACTACTGTGCTAAG |
| **VH**: JH4 | GACTACTGGGGCCAAGGAACCCTGGTCACCGTCTCCTCAGGATCC |

**SI Table 4.** Flow cytometry labelling conditions used in this study.

| Flow cytometry labelling for full-length characterization | | |
|---|---|---|
| **Epitope (detected feature)** | **Primary Label (dilution)** | **Secondary Label (dilution)** |
| HA epitope tag (Display levels) | Mouse anti-HA (1:500) | Goat anti-mouse Alexa Fluor 488 (1:500) |
| c-Myc epitope tag (Full length protein) | Chicken anti-c-Myc (1:500) | Goat anti-mouse Alexa Fluor 647 (1:500) |

| CuAAC reactions | | | |
|---|---|---|---|
| **ncAA** | **Reaction** | **First step molecule** | **Second step molecule** |
| **AzF** | Small molecule click | Small molecule-alkyne | Biotin-alkyne |
| | Biotin click | N/A | Biotin-alkyne |
| **OPG** | Small molecule | Small molecule-azide | Biotin-azide |
| | Biotin click | N/A | Biotin-azide |

| Flow cytometry labelling for CuAAC characterization | | |
|---|---|---|
| **Epitope (detected feature)** | **Primary Label (dilution)** | **Secondary Label (dilution)** |
| c-Myc epitope tag (Full length protein) | Chicken anti-c-Myc (1:500) | Goat anti-mouse Alexa Fluor 647 (1:500) |
| Biotin (clicked molecules) | N/A | Streptavidin Alexa Fluor 488 (1:500) |

| Flow cytometry labelling for IgG binding characterization | | |
|---|---|---|
| **Epitope (detected feature)** | **Primary Label (dilution)** | **Secondary Label (dilution)** |
| c-Myc epitope tag (Full length protein) | Chicken anti-c-Myc (1:500, 50 µL) | Goat anti-mouse Alexa Fluor 647 (1:500) |
| Biotinylated antigen | N/A | Streptavidin Alexa Fluor 488 (1:500) -OR- Mouse anti-biotin PE (1:500) |
| Donkey IgG native epitope (Donkey IgG protein) | N/A | Rabbit anti-donkey DyLight 488 (1:500) |

**SI Table 5.** Primers used to construct and sequence secretion vectors.

| Primer Name | Sequence (5' to 3') |
|---|---|
| pRS314-scFv-fwd | TTGGACAAGAGAGAAGCTCGGCCGGCTAGCGACATACAGA TGACTCAAAGTCCCAGTTCA |
| pRS314-scFv-rev | TTCTTCAGAAATAAGCTTTTGTTCGGATCCTGAGGAGACGG TGACCAGGGTTCCTTGGCC |
| pRS314-SeqPrimer-fwd | GTTGGCTATCTTCGCTGC |
| pRS314-SeqPrimer-rev | GTACAGTGGGAACAAAGTC |
| pCHA-scFv-NheI-fwd | CCATACGACGTTCCAGACTACGCTGCTAGCGACATACAGAT GACTCAAAGTCCC |
| pCHA-scFv-XmaI-rev | TTTGTCGGAACTTTTAGGTTCTACCCCGGGTGAGGAGACGG TGACCAGGGTTCCTTG |
| scFvFc-seq-Fwd | TGCCATTGGCCTTAGCTCAACCGG |
| scFv-Fc-seq-Rev | CGGCTTTGGCGGGAACAAAAAGACG |
